# Maize root system stiffness is determined by the size and distribution of the below-ground root system

**DOI:** 10.64898/2026.02.05.704061

**Authors:** Ashley N. Hostetler, Emilia Pierce, Jonathan W. Reneau, David Griffin, Erin E. Sparks

## Abstract

Root system stiffness is a measurement that describes the resistance of a root system to rotation. Previous work has shown that genotypes with a higher root system stiffness are also more susceptible to root lodging. In this study, we deconvolve the contribution of root architecture, individual root-level mechanics, and root-soil interaction to root system stiffness in maize inbred lines. Collectively, larger above- and below-ground root systems and stiffer individual brace roots contribute to a higher root system stiffness. When considering all traits in predictive models, the below-ground root architecture primarily drives the prediction of root system stiffness categories. These below-ground traits describe the size and distribution of the root system. Analysis of a *roothairless3* mutant revealed a reduction in root system stiffness that could not be decoupled from a reduction in root size. Together these results link root system stiffness to the size and distribution of the below-ground root system and highlight the importance of this measurement for both root lodging resistance and root phenotyping.

**HIGHLIGHT:** The size and spatial distribution of below-ground roots are the strongest predictors of maize root system stiffness, providing new insight into plant mechanics and the relationship with root lodging susceptibility.

## INTRODUCTION

Plant root systems perform essential functions, including acquiring water and nutrients, facilitating symbiotic interactions, and providing mechanical anchorage to maintain plant stability. A failure of anchorage is called root lodging, and has devastating impacts on agricultural crop production (Berry et al., 2004; Rajkumara, 2008). In maize, root lodging can cause between 5-43% yield losses depending on the plant growth stage, with root lodging at flowering causing the greatest impacts (Lindsey et al., 2021). Severe weather events, such as the 2020 Derecho that swept through Iowa USA, caused losses exceeding 80% (Barten et al., 2022). Despite the devastating impacts, root lodging resistance is difficult to study due to the multiple interacting factors involved.

Root lodging occurs when the external bending moment exceeds the resistance of the root system to rotation, resulting in mechanical failure of the root system through breaking, buckling, or pulling from the soil. The external bending moment is determined by the interaction of wind with the height of the stalk, the mechanics of the stalk, and the drag characteristics of the stalk (Stubbs et al., 2023; Hostetler et al., 2025). Conversely, root system resistance to rotation is determined by mechanical aspects of the root system. Root system mechanics can be divided into two sequential deformation modes: elastic deformation, which is non-destructive and recoverable, and plastic deformation, which is irreversible and may involve structural damage (Ikiriko et al., 2025)). Elastic deformation is measured as root system stiffness, which describes the relationship between displacement and force. Plastic deformation is measured as root system strength, which reports the force at failure. Prior work has associated root architectural traits with differences in anchorage strength and root lodging resistance (Stamp and Kiel, 1992; Sanguineti et al., 1998; Liu et al., 2012; Reneau et al., 2020; Hostetler et al., 2022b; Ren et al., 2022). These associations have identified both below-(crown root system) and above-ground (brace root system) root systems as important factors for lodging-resistance in maize. A genome-wide association analysis expanded upon these findings and showed that root architecture explained 45% of the variation in root lodging after the 2020 Derecho (Zheng et al., 2023). However, this analysis also highlighted the potential for the cell wall to contribute to root lodging (Zheng et al., 2023).

The cell wall properties will modify mechanics at the individual root-level. However, the assessment of individual root-level mechanics in maize has been limited (Ikiriko et al., 2025). Foundationally, mechanics is assessed by applying a displacement and measuring a force. Displacements can be applied in tension (Beck et al., 1988; Ennos et al., 1993; Chimungu et al., 2015), compression (not reported for maize due to the difficulty of measurement), or bending (Ennos et al., 1993; Goodman and Ennos, 1998; Hostetler et al., 2022a). Among these, bending is considered the easiest and most high-throughput characteristic to quantify (Ikiriko et al., 2025). The most common type of bending test is 3-point bending, where the root sits on two lower anvils and an upper anvil induces displacement at the mid-point of the root. From this assay, the structural stiffness (*K*) reports the initial relationship between force and displacement (Niklas and Spatz, 2012). *K* is a combination of the second moment of area (*I*), which describes the shape of the root, and the bending modulus (*E*), which describes the underlying material properties of the root. Modifications in the cell wall would change the *K* through changes in *E*. Beyond these measurements, root strength describes the force at failure (breaking). However, there have been limited attempts to link root-level mechanics with overall root system mechanics. Among prior studies, one showed a positive association of root tensile strength with uprooting resistance (Beck et al., 1988). However, this study did not attempt to deconvolve the contribution of root shape from root material properties. These results suggest that individual root-level mechanics play an important role in anchorage, but this role has not been fully explored.

We recently reported a tool that non-destructively measures the root system stiffness of maize plants and showed that maize hybrids with a higher root system stiffness were more susceptible to root lodging (Hostetler et al., 2025). This outcome is consistent with a stiffer root system being more brittle and reaching failure strength at lower displacements. However, the underlying contributions of root architecture, individual root-level mechanics, and root-soil interactions that determine root system stiffness remain unknown.

In this study, we address this gap to determine the factors that contribute to root system stiffness. Using a small diversity panel of maize inbred lines, we first showed that the results from maize hybrids were translatable to maize inbred genotypes, with a higher root system stiffness associated with increased root lodging susceptibility. We then assessed phenotypic and mechanical characteristics of the root systems. These included root crown excavation and characterization of below-ground root architecture, integration of previously published brace root architectures (Hostetler et al., 2022b), and 3-point bend mechanical testing of brace roots. For each of these analyses, larger root architectures (above- and below-ground) and stiffer brace roots were associated with a higher root system stiffness. When considering all significant traits in random forest models, below-ground root architectural traits primarily drove the prediction of root system stiffness. Further analysis identified below-ground traits describing the size (e.g. maximum diameter, steep angle frequency, and surface area) and distribution (solidity) of the root system as top predictors. To assess the potential for rhizosphere interactions playing a role in anchorage, we quantified the root system stiffness of *roothairless3* mutants. While these mutants showed a reduction in root system stiffness compared to wild-type siblings, this was attributed to a smaller root system. Collectively, this work shows that the below-ground root architecture is the major contributor to root system stiffness in maize.

## MATERIALS AND METHODS

### Plant Material & Field Management

Forty-three maize (*Zea mays*) inbred genotypes were selected as a small diversity panel (**Table S1**). Twenty-one were founder lines from the Nested Association Mapping population (McMullen et al., 2009) and the remaining 22 were lines selected to capture additional genetic variation (Hostetler et al., 2022b). Inbred experiments were conducted in Newark, DE USA (39°40’ N and 75°45’ W) in 2017, 2018, 2019, 2020, 2021, and 2022. Seeds were planted on 18 May 2017, 30 May 2018, 22 May 2019, 1 June 2020, 5 May 2021, and 23 May 2022.

Delaware fields were treated with pre-emergence (Lexar at 3.5 quart per acre; Simazine at 1.2 quart per acre) and post-emergence (Accent at 0.67 ounces per acre) herbicide, soil insecticide (COUNTER 20 G at 5.5 lbs per acre), and fertilizer (Ammonium Sulphate 21-0-0 at 90 lbs per acre; 30% urea ammonium nitrate 40 gallons per acre). Daily weather data can be located at the Delaware Environmental Observing System (https://www.deos.udel.edu) and selecting the Newark, DE-Ag Farm Station.

In 2025, the *roothairless3* (*rth3*) mutant (Wen and Schnable, 1994; Hochholdinger et al., 2008) was grown in Columbia, MO USA (38°91’ N and 92°28’ W). The *rth3* mutant seeds were obtained from the Maize Genetics Coop (Woodhouse et al., 2021) and propagated through self-fertilization. Seeds were sterilized in a 50% bleach solution for 10 min, rinsed with sterile water, and then placed in pre-dampened A4 germination paper. Germination rolls were placed in a beaker of water inside a benchtop growth chamber for seven days with a 12-hour light-dark cycle. After one week, seedlings were checked for the presence or absence of the root hair phenotype, and four segregating lines were transplanted to the field on 9 June 2025.

The Missouri field site was treated with pre-emergence (Degree Xtra at 3.7 quarts per acre) and post-emergence (Atrazine at 1 quart per acre) herbicide. NPK 160-0-40 fertilizer was applied pre-planting at a rate of 160 lbs N per acre and 40 lbs of K per acre. Weather data for the Columbia, MO USA field site can be located at Weather Underground (https://www.wunderground.com/dashboard/pws/KMOCOLUM110).

All locations and years utilized a 12 ft plot length with 18 seeds for inbred genotypes and 8 seedlings for mutant genotypes.

### Experimental Design and Data Collection

#### Quantification of Root System Stiffness

We previously reported a tool that non-destructively measures the root system stiffness of large grain crops, called Sorghum and Maize Under Rotational Force (SMURF) (Hostetler et al., 2025). Briefly, this device was attached to plant stalk with straps. The associated software was used to start a test. The device first balanced its own weight and tared a load cell, which measures forces. The device then extended and displacement and force were recorded. The root system stiffness was extracted from the force-displacement curve.

In 2020 and 2021, root system stiffness was quantified on the inbred genotypes at reproductive maturity. In 2020 Tropical Storm Isaias caused widespread root lodging at 65 days after planting (dap). Data was collected from 13 genotypes that were not root lodged at 128 dap. In 2021, data was collected from all inbred genotypes at 130-131 dap. 2020 data collection included 1-2 plots per genotype and at least three plants per plot. 2021 data collection included 2-4 replicate plots, and at least two plants per plot. In 2025, the root system stiffness was quantified on *rth3* mutant and wild-type sibling plants at 95 dap (84 days after transplanting).

Inbred genotypes were assigned to root system stiffness categories (low, moderate, high) using a previously reported approach (Hostetler et al., 2022b). Briefly, a genotypic mean was determined for each of the genotypes from the 2021 quantification of root system stiffness. Data was then scaled and centered, and the population mean and standard deviation were determined. Genotypes that had a genotypic average greater than one standard deviation below the population mean were categorized as having a low root system stiffness, while genotypes that had a genotypic average greater than one standard deviation above the population mean were categorized as having a high root system stiffness. Genotypes with a genotypic mean within one standard deviation of the population mean were categorized as having a moderate root system stiffness.

#### Characterization of Root Lodging Susceptibility

Root lodging susceptibility data for inbred genotypes were obtained from (Hostetler et al., 2022b) based on the aftermath of the 2020 Tropical Storm. Data were subset to include the lines in this study. Genotypes were assigned into root lodging susceptibility categories (low, moderate, high) using the same approach described above for root systems stiffness categories.

#### Quantification of Brace Root Emergence

In 2021 and 2022, brace root emergence and leaf count at time of brace root emergence were recorded. Brace root emergence was determined when 50% of the plants in a plot had visible roots emerging from above-ground stem nodes.

#### Quantification of the Below-ground Root Architecture

In 2017, the below-ground root architecture of inbred genotypes was characterized at reproductive maturity. The root crowns were excavated from 1-2 plots per genotype and 1-3 plants per plot at 146 dap. After the root crowns were manually excavated, residual soil was removed by a rubber-tipped air nozzle connected to a 30-gallon air compressor. After cleaning, root crowns were placed on a dark-colored background with a scale marker and imaged with a Canon EOS Rebel T6.

Root images were manually thresholded and run through RhizoVision Explorer (Seethepalli et al., 2021). Briefly, regions of interest were defined to capture only the root system that entered the soil. The images were analyzed using the whole root analysis and a pruning setting of 10. Traits were grouped into general categories, which described the distribution, shape, extent, and size of the root system (**Table 1**). Trait descriptions can be found in (Seethepalli et al., 2021).

**Table 1.**
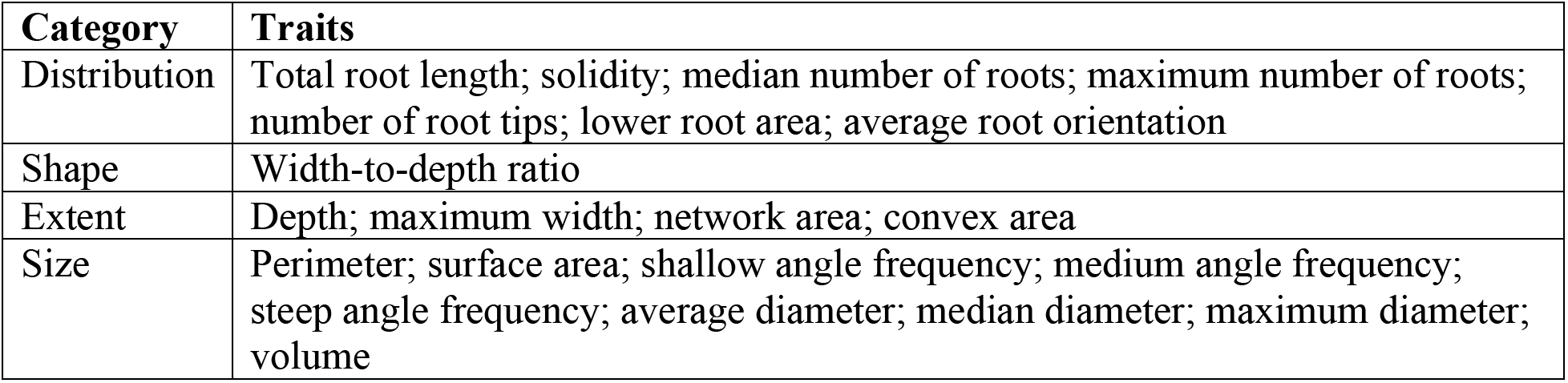
Categorization of Root Architecture Traits.

In 2025, the *rth3* mutant and wild-type sibling plants that were quantified for root system stiffness were manually excavated, washed with water, and imaged as described above. The maximum width was measured from root crown images in FIJI (Schindelin et al., 2012).

#### Quantification of the Above-ground Root Architecture

In 2019, above-ground brace root traits were quantified and previously published (Hostetler et al., 2022b). The following traits were used from this previous study: number of nodes in the soil, single brace root width, stalk width, brace root height on stalk, spread width, and root angle.

#### Quantification of Brace Root Mechanics

In 2018 and 2019, brace roots were collected to quantify individual root-level mechanics at reproductive maturity. Brace roots that entered the soil were excised with shears and stored in tassel bags until testing. Roots were trimmed to 20 mm, and 3-point bend tests were performed on the Instron 685C-05 Universal Testing Stand as previously described (Hostetler et al., 2022a; Pierce et al., 2025). Briefly, the brace root length, diameter perpendicular to the upper anvil, and diameter parallel to the upper anvil were measured using a digital caliper. The root was then placed on the lower anvils for testing. Roots were pre-loaded to 0.2 N and tested at 1 mm/min until 0.3 mm.

Data was exported from Bluehill software and run through a curation pipeline, where the slope of the force-displacement curve was extracted as the structural stiffness (*K*). Displacement data less than 0.1 mm were used to run the linear regression. The second moment of area (*I*) was calculated based on the root geometry and span length as:

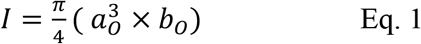

where *π* = 3.1415, *a*_0_ is the minor radius axis parallel to the plane of displacement, and *b*_0_ is the major radius axis perpendicular to the plane of displacement.

The bending modulus (*E*) was then calculated as:

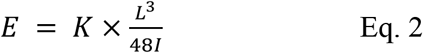

Where *K* is the structural stiffness, L is 17.5 mm (fixture span length used in this setup), and *I* is the second moment of area calculated from Eq. 1.

All scripts for processing these data were previously published in (Pierce et al., 2025) and available at https://doi.org/10.5281/zenodo.15014302.

### Statistics

All statistical analyses were performed using R ver. 4.0.2 (R Core Team). Analysis of variance (ANOVA) models were used for statistical analyses. Prior to running each ANOVA, a Shapiro-Wilk test was used to test the normality of residuals. If residuals were not drawn from a normal distribution, a Tukey’s Ladder of Powers (*rcompanion* package ver. 2.4.1) (Mangiafico, 2021) transformation was used. If a *p* value less than 0.05 was associated with the F statistic, the analyses were considered significant and a post-hoc Tukey honest significant difference (HSD) test was used to test all pairwise comparisons. All figures were generated with the ggplot2 (ver. 3.5.2) (Wickham, 2016) and the cowplot (ver. 1.1.3) (Wilke, 2015) packages.

The following models were run, displayed as “dependent variable ∼ independent variable*independent variable”. (1) root system stiffness ∼ genotype, (2) root system stiffness ∼ genotype*year, (3) root system stiffness ∼ root lodging susceptibility category, (4) below-ground root architecture traits ∼ genotype, (5) below-ground root architecture traits ∼ root system stiffness category, (6) above-ground root architecture traits ∼ root system stiffness category, (7) brace root mechanics ∼ genotype*node, and (8) brace root mechanics ∼ root system stiffness category*node. For models 4, 5, 6, 7 and 8, individual models were run for each trait within that category.

Principal component analyses (PCA) were used to reduce dimensionality, identify the most important traits within data types, and visualize clustering patterns. A PCA was run on the below-ground root architecture traits and the above-ground root architecture traits individually. The ‘prcomp’ function from the base R *stats* package (ver. 4.4.0) was used to run the PCA, and the *ggfortify* package (ver. 0.4.17) (Tang et al., 2016) was used to plot the PCA. For both PCAs, data was scaled prior to running the PCA and root system stiffness categories were overlaid on points to visualize clustering.

Supervised random forest modeling approaches were used to identify the most important traits in predicting root system stiffness categories. Two models were tested with root system stiffness category as the outcome, and (1) all significant traits from analysis variance models as predictors, and (2) all significant traits from below-ground root architecture traits as predictors. Random forest modeling was performed using the following packages: *tidymodels* (ver. 1.3.0) (Kuhn and Wickham, 2020), *ranger* (ver. 0.17.0) (Wright and Ziegler, 2017), *kkn* (ver. 1.4.0) (Schliep and Hechenbichler, 2025), and *randomForest* (ver. 4.7.1.2) (Liaw and Wiener, 2002).

Genotypic averages for each trait were used in the first random forest model and plant level data was used for each trait in the second random forest model. For both models, data was then randomly split into an 80% training and 20% testing data set stratified by the root system stiffness category. Prior to the final model, model training and hyperparameter tuning were used to identify the most appropriate model. Specifically, the random forest was configured to grow 1000 trees, and the following parameters were optimized: number of variables sampled at each split and minimum node size. The performance of the model was estimated using five-fold cross-validation and repeated three times. The final model was fit on the training and testing set and model accuracy was determined. Predictor importance was determined using the mean decrease in Gini, where the highest mean decrease in Gini was considered the most important predictors of the model (Menze et al., 2009).

To determine if root hairs contributed to the root system stiffness, two ANOVA models were tested. Specifically, genotype (*rth3* mutant or wild-type sibling) was the independent variable in both models and either root system stiffness or maximum crown width was the dependent variable. To determine if the root system stiffness of *rth3* plants was different than expected given the root system size, data was scaled across environments to compare to inbred genotypes. Specifically, a scaling factor was calculated by dividing the mean trait value of the wild-type sibling (quantified in MO) by the mean trait value of B73 (quantified in DE) for both root system stiffness and maximum crown width. Then the *rth3* mutant and wild-type sibling values were divided by the scaling factor to estimate their expected values in the same environment as the inbred genotypes.

## RESULTS

### Root system stiffness varied among inbred genotypes and is associated with root lodging susceptibility

In maize hybrids, genotypes with a higher root system stiffness are associated with greater susceptibility to root lodging (Hostetler et al., 2025). To determine if this relationship extends beyond hybrids, the root system stiffness of 43 maize inbred genotypes was measured across two years. In 2020, Tropical Storm Isaias caused extensive root lodging, and data collection was only possible for 13 of 43 genotypes. Despite the reduced genotypes, there was a significant effect of genotype on root system stiffness for the lines that were assessed (**Table S2**). In 2021, all 43 inbred genotypes were measured and there was again a significant effect of genotype on root system stiffness (**Figure 1A, Table S3**). To determine the consistency of root system stiffness between years, the data were compared for the 13 lines measured in both years and showed a high correlation (r= 0.91, **Figure S1A**). While the root system stiffness varied between years (**Figure S1B, Table S4**), the relationship between genotypes was maintained across years with high root system stiffness genotypes being high in both years (**Figure S1C**). This result is the same as reported for hybrids (Hostetler et al., 2025). Given that relationships among inbred genotypes were consistent, genotypes were grouped into three root system stiffness categories (low, moderate, or high) stiffness to enable multi-year analyses (**Figure 1A**).

**Figure 1.**
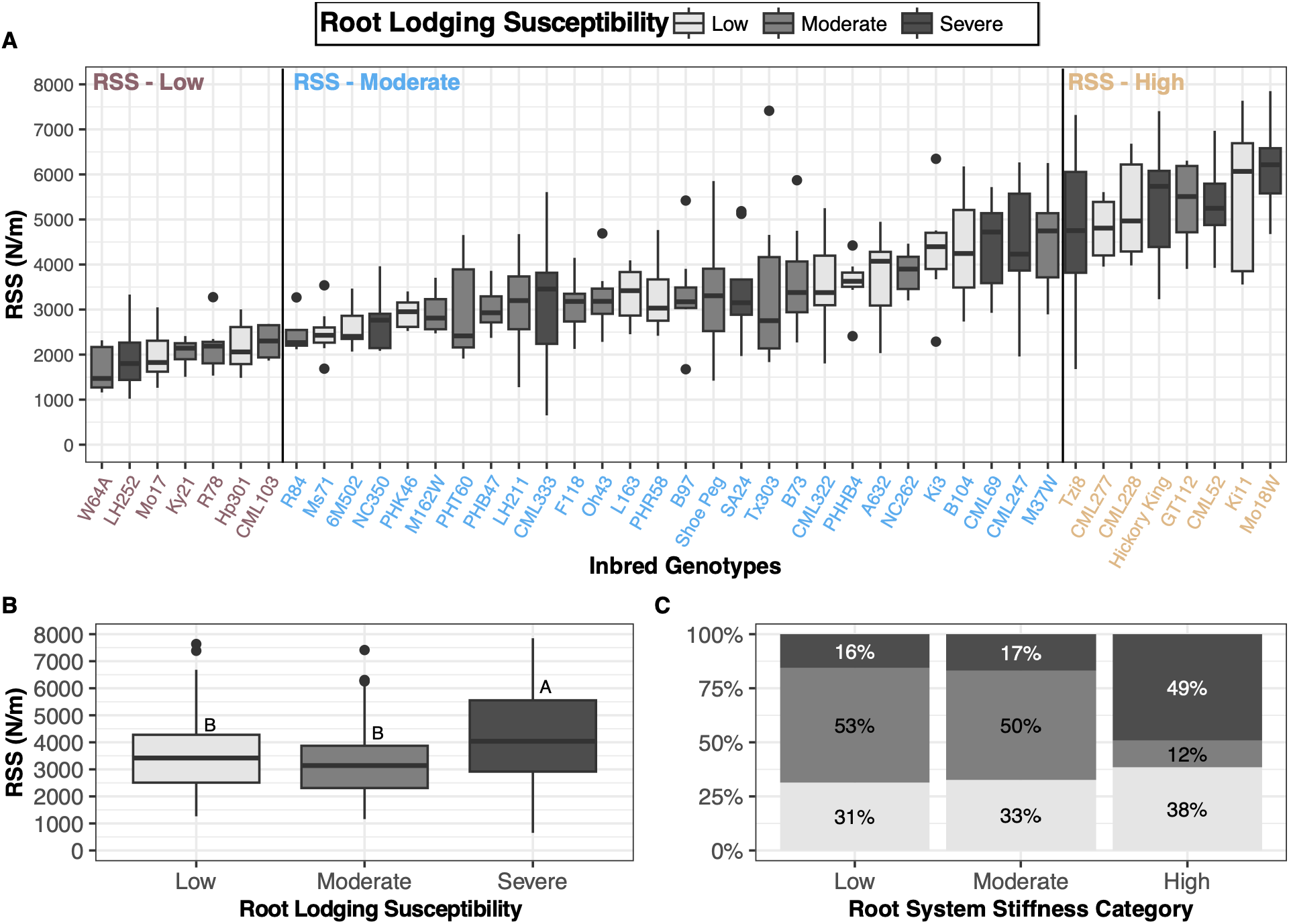
The root system stiffness varies among inbred genotypes and is associated with root lodging susceptibility. (A) Inbred genotypes used in this study have variation in root system stiffness. Inbred genotypes were categorized as having a low (maroon), moderate (blue), or high (gold) root system stiffness relative to the other inbred genotypes used in this population. Inbred genotypes were also categorized as having a low (light grey), moderate (grey), or severe (dark grey) root lodging susceptibility relative to the other genotypes used in this population. (B) Root lodging susceptibility categories significantly differed for root system stiffness, with those having a higher root system stiffness being more susceptible to root lodging and those with a lower root system stiffness being less susceptible to root lodging. Root lodging susceptibility categories that share a letter do not significantly differ from one another (Tukey HSD assessed at *P* < 0.05). (A-B) Black dots indicate outliers (C) The inbred genotypes with a high root system stiffness had 49% of the inbreds categorized as severe, 12% as moderate, and 38% as low to root lodging susceptibility. The inbred genotypes with low and moderate root system stiffness had similar distributions across severe (16% and 17%, respectively), moderate (53% and 50%, respectively), and low (31% and 33%, respectively) root lodging susceptibility. RSS = root system stiffness.

To determine if differences in root system stiffness were associated with root lodging susceptibility, genotypes were grouped into percent root lodging categories. As previously reported for hybrids (Hostetler et al., 2025), inbred genotypes with a higher root system stiffness were more susceptible to root lodging compared to those with a lower root system stiffness (**Figure 1B, Table S5**). These results expand upon the finding from hybrid genotypes to indicate that a stiffer root system will fail more frequently in maize. However, root system stiffness categories do not perfectly overlap with root lodging susceptibility categories (**Figure 1A** and **1C**). Thus, this finding is consistent with root system stiffness as only one of the factors influencing root lodging susceptibility.

### Larger below- and above-ground root systems are associated with higher root system stiffness

To quantify the link between root system stiffness and root architecture, the below-ground and above-ground root systems were characterized. First, root crowns were excavated and analyzed for 21 traits (**Table 1**), all of which were significantly affected by inbred genotype (**Figure S2, Table S6**). To determine if these traits were able to distinguish between the different root system stiffness categories, a PCA was used and revealed distinct clusters for the low and high root system stiffness categories (**Figure 2A, Figure S3**). Each trait was then evaluated for differences between the root system stiffness categories (**Figure 2B-D, Figure S4, Tables S7**). Fifteen traits, describing the distribution, extent, and size of the root system, showed differences between the root system stiffness categories (**Figure 2B-D**). For all significant traits, genotypes with a high root system stiffness had larger below-ground traits. Separation of root system stiffness categories was the greatest when comparing the extremes of root system stiffness, i.e., low versus high.

**Figure 2.**
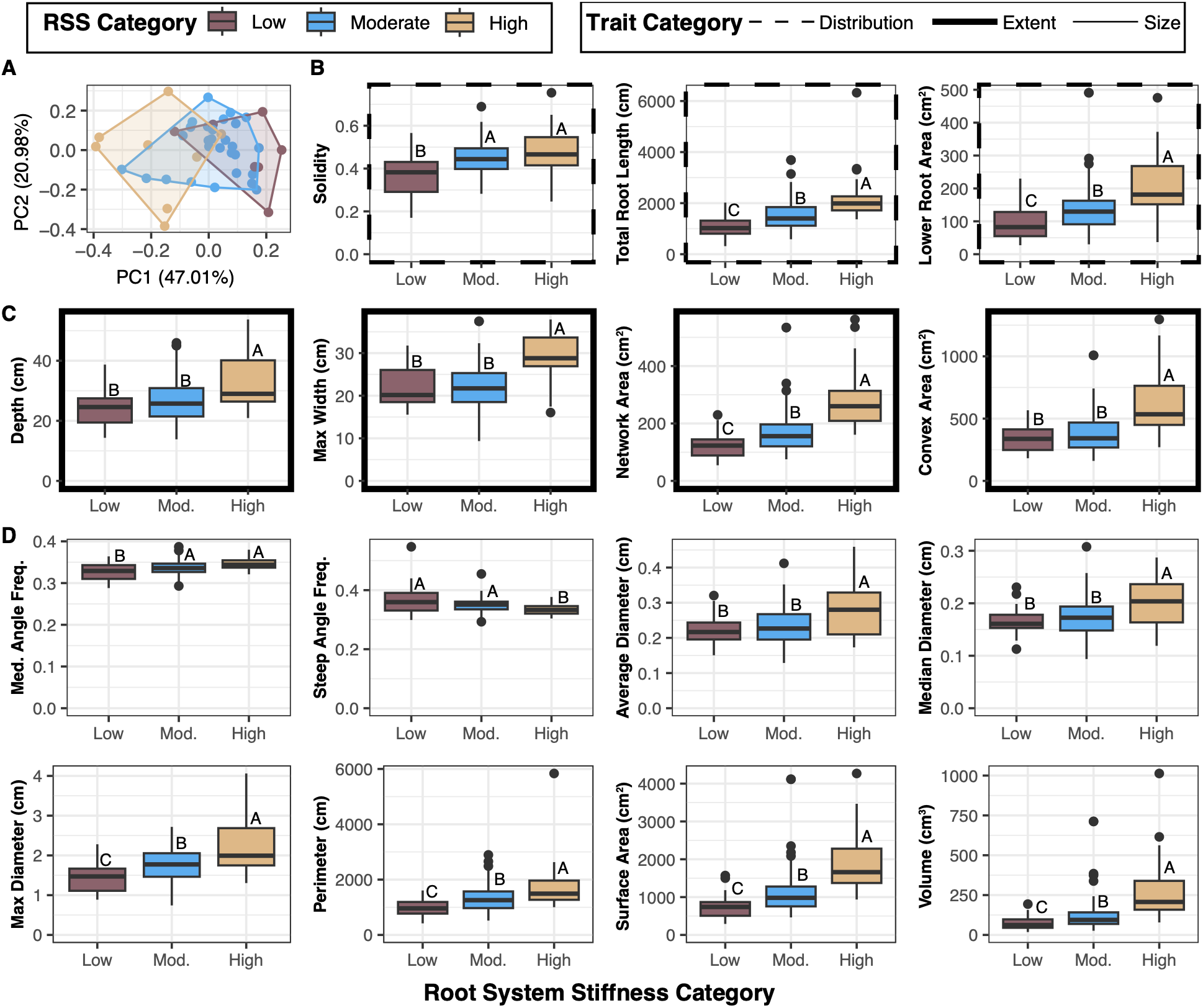
The below-ground root system varies among root system stiffness categories. (A) A PCA of below-ground root traits separates the high from the low root system stiffness category along PC1. (B) Traits describing the distribution of the root system were significantly different among root system stiffness categories. This included solidity, total root length, and lower root area. (C) Traits describing the extent of the root system were significantly different among root system stiffness categories. This included depth, max width, network area, and convex area. (D) Traits describing the size of the root system were significantly different among root system stiffness categories. This included medium angle frequency, steep angle frequency, average diameter, median diameter, maximum diameter, perimeter, surface area, and volume. Root system stiffness categories that share a letter are not significantly different from one another (Tukey HSD assessed at *P* < 0.05). Black dots indicate outliers. RSS = root system stiffness. Mod = moderate.

Above-ground brace root architectural variation for the inbred genotypes used in this study has been previously reported (Hostetler et al., 2022b). Six previously published brace root traits were analyzed here in the context of root system stiffness categories (**Figure S5**). A PCA of brace root traits revealed distinct clusters for low and high root system stiffness categories along PC1 (**Figure 3A, Figure S6**). However, separation of root system stiffness categories by brace root traits explained less of the variation (27.45%; **Figure 3A**) than the below-ground traits (47.01%; **Figure 2A**). At the individual trait level, brace root angle, brace root width, brace root spread width, brace root height on stalk, and stalk width significantly differed between the root system stiffness categories (**Figure 3B-F, Table S8**). The number of brace root nodes in the soil was not significantly different between categories (**Figure S7**). Overall, inbred genotypes with a larger above-ground root system were associated with the high root system stiffness category (**Figure 3, Figure S6**). Collectively, these data demonstrate that larger below- and above-ground root systems are associated with a higher root system stiffness.

**Figure 3.**
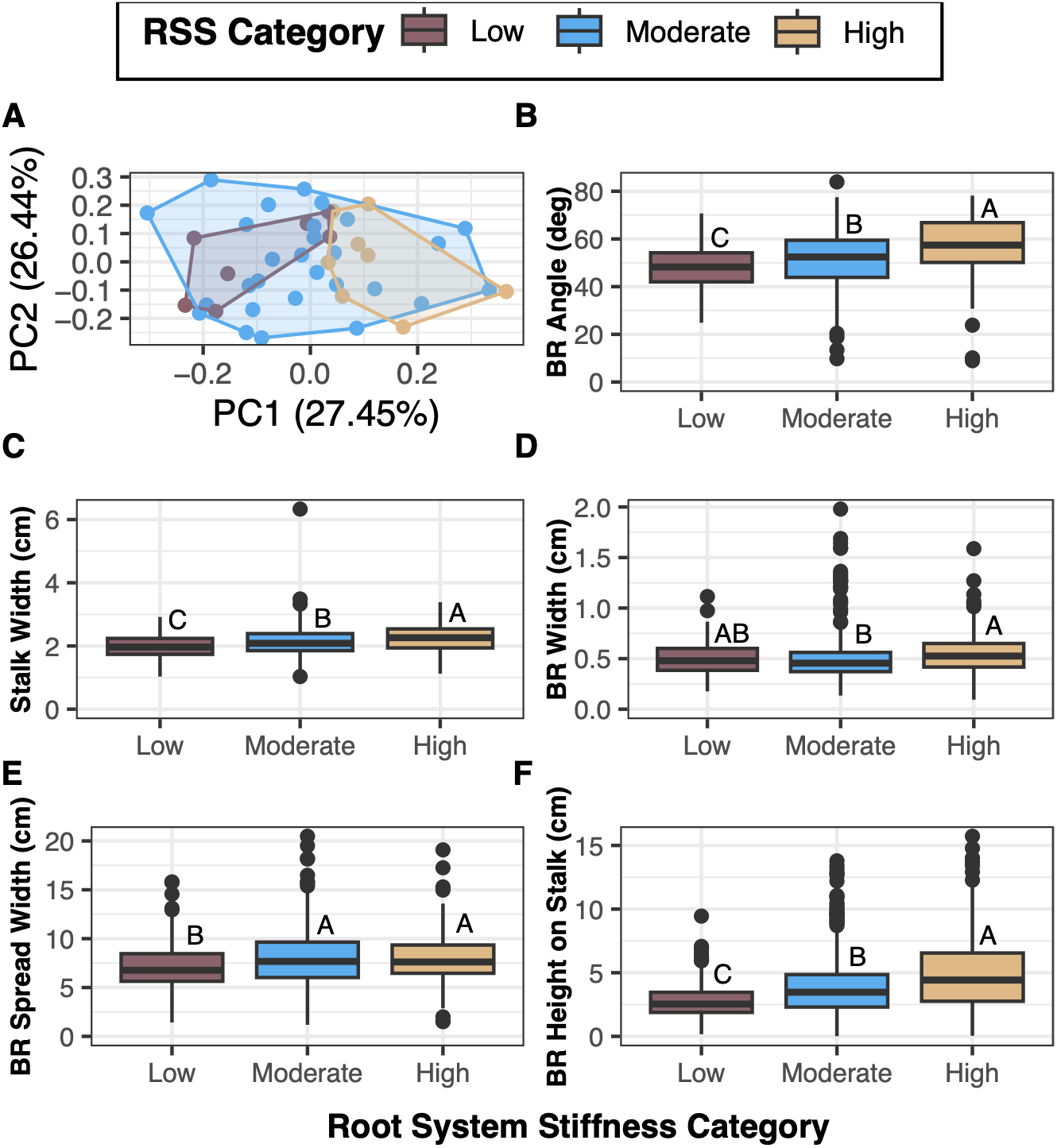
The above-ground root system varies among root system stiffness categories. (A) A PCA of brace root traits separates the high from low root system stiffness categories along PC1. (B-F) Brace root traits that are statistically different among root system stiffness categories. These included (B) brace root angle, (C) stalk width, (D) brace root width, (E) brace root spread width, and (F) brace root height on stalk. Root system stiffness categories that share a letter are not significantly different from one another (Tukey HSD assessed at *P* < 0.05). Black dots indicate outliers. BR = brace root. RSS = root system stiffness.

### Delayed brace root emergence is associated with higher root system stiffness

The timing of brace root emergence on root architecture and root system stiffness was next examined. Across two-years, brace roots emerged between 28 and 62 dap and was dependent on genotype, year, and the interaction (**Figure S8, Table S9**). When considering brace root emergence date by root system stiffness categories, there was an effect of root system stiffness category and year, but no interaction. Specifically, the higher root system stiffness category had later brace root emergence regardless of year (**Figure 4A, Table S10**). The later emergence date was also associated with emergence at a later vegetative growth stage (**Figure 4B**). Thus, differences in brace root emergence cannot be attributed to differences in plant growth rate.

**Figure 4.**
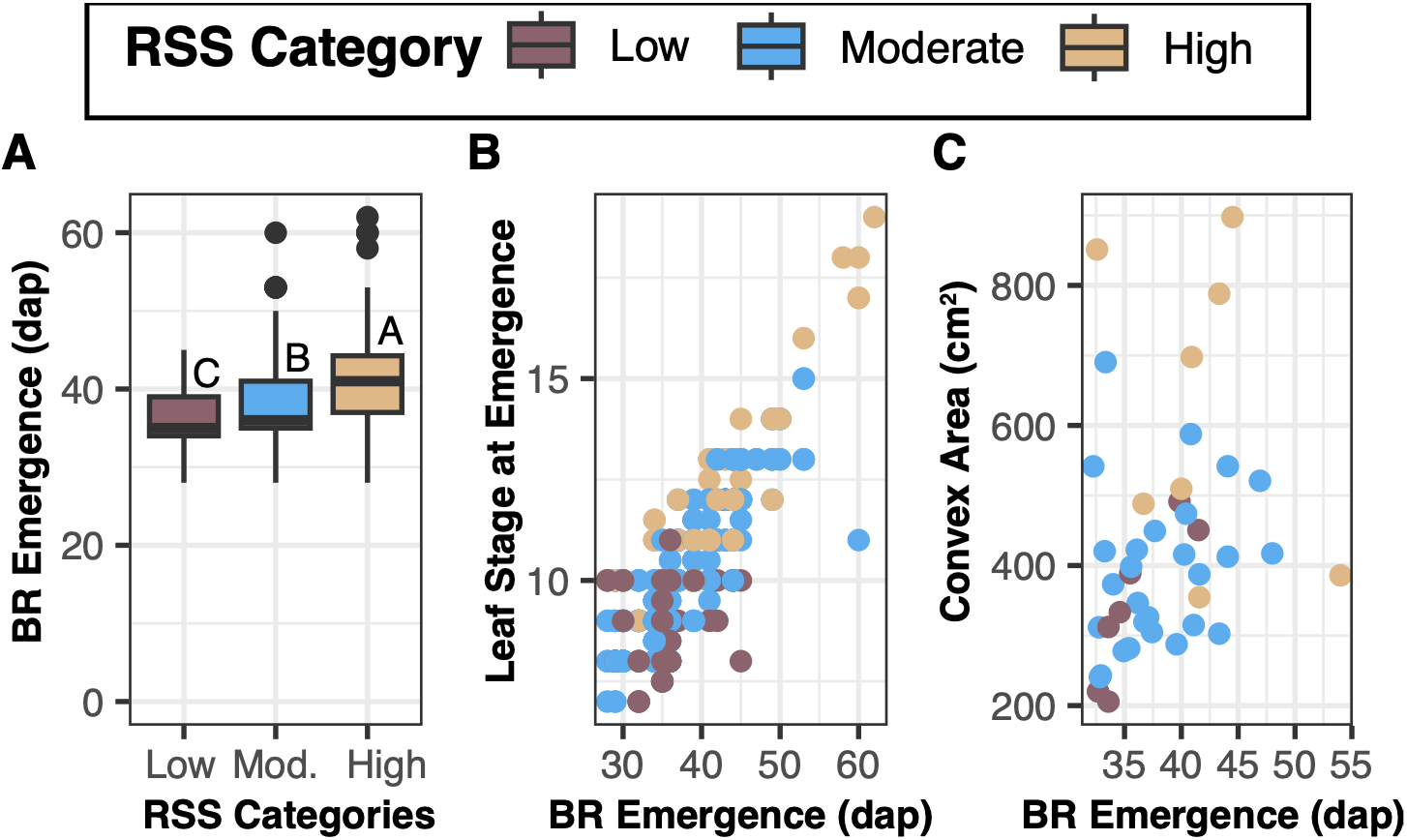
Brace root emergence is associated with root system stiffness and below-ground root system size. (A) Later brace root emergence is associated with a higher root system stiffness, whereas earlier brace root emergence is associated with a lower root system stiffness. Root system stiffness categories that share a letter are not significantly different from one another (Tukey HSD assessed at *P* < 0.05). Black dots indicate outliers. (B) Brace root emergence was positively correlated with the leaf stage at timing of brace root emergence. (C) Brace root emergence was positively correlated with below-ground root system convex area. RSS = root system stiffness. BR = brace root. Mod = moderate.

We hypothesized that later brace root emergence would enable more investment into the below-ground root system. To test this hypothesis, the relationship between brace root emergence and root crown convex area was investigated. Indeed, inbred genotypes with a later emergence date of brace roots had larger belowground convex root system area (**Figure 4C**). Thus, delayed brace root emergence is associated with a greater investment in the below-ground root system.

### Brace root structural stiffness is associated with root system stiffness and is driven by changes in geometry

In addition to the root architecture, the potential influence of individual root-level mechanics on overall root system stiffness was assessed. Because direct mechanical testing of subterranean roots is limited by their underground location, brace root mechanics were quantified with 3-point bending. Consistent with previous work (Hostetler et al., 2022a), brace root structural stiffness (*K*), second moment of area (*I*), and bending modulus (*E*) all vary by node, genotype, and their interaction (**Figure S9, Table S11**). In assessing the differences between root system stiffness categories, brace root *K*, *I* and *E* all vary by node and genotype, but not their interaction (**Figure S10, Table S12-S13**). However, *K* and *I* varied between the high and low root system stiffness categories while *E* did not. Plants with a high root system stiffness had brace roots with a higher *K* (**Figure 5A**) and a larger *I* (**Figure 5B**) compared to those with a low root system stiffness. Since *E* did not vary between the high and low root system stiffness categories, these data demonstrate that the differences in brace root mechanics associated with root system stiffness categories are due to the larger root size, as opposed to changes in the composition of the roots. Thus, capturing root architectural variation will also capture the root mechanical contribution to root system stiffness.

**Figure 5.**
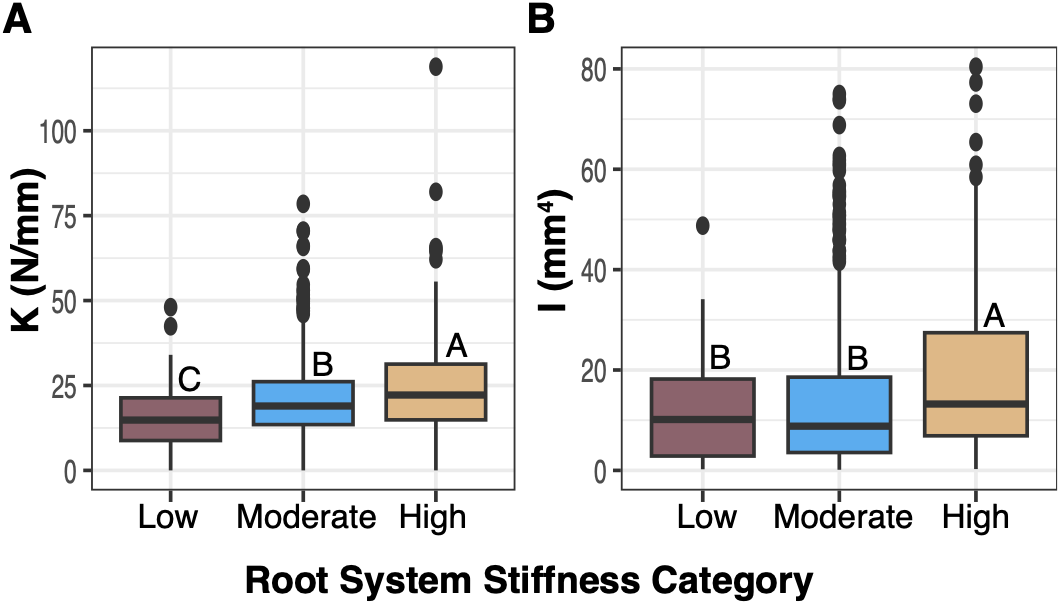
*K* and *I* vary between root system stiffness categories. (A) Inbred genotypes that had a high root system stiffness had a larger structural stiffness (*K*) compared to those with a low root system stiffness. (B) Inbred genotypes that had a high root system stiffness had a larger second moment of area (*I*) compared to those that had a low root system stiffness. Root system stiffness categories that share a letter are not significantly different from one another (Tukey HSD assessed at *P* < 0.05). Black dots indicate outliers.

### Below-ground root architectures are the primary contributor to root system stiffness

Among all three types of root system features – below-ground root architecture, above-ground root architecture, and brace root mechanics – there were traits that varied between the root system stiffness categories. To understand which of these features were most predictive of root system stiffness, random forest modeling approaches were employed (**Table S14**). First, genotypic averages within a year were used to identify which feature types (below-ground root architecture, above-ground root architecture or above-ground mechanics) were most predictive of root system stiffness categories. The seven traits (25%) with the highest variable importance were all below-ground root architecture traits (**Figure 6A**). To determine if this is an artifact of the greater number of below-ground architecture traits (n=15) tested compared to above-ground architecture (n=5) or above-ground mechanics (n=6), the mean decrease in Gini values were normalized by root system feature type. This approach highlighted that below-ground architecture traits had greater normalized variable importance (0.70) compared to above-ground architecture traits (0.34) or above-ground mechanics traits (0.37). Thus, when all feature types were considered together, below-ground root architecture contained the greatest predictive information for root system stiffness category.

**Figure 6.**
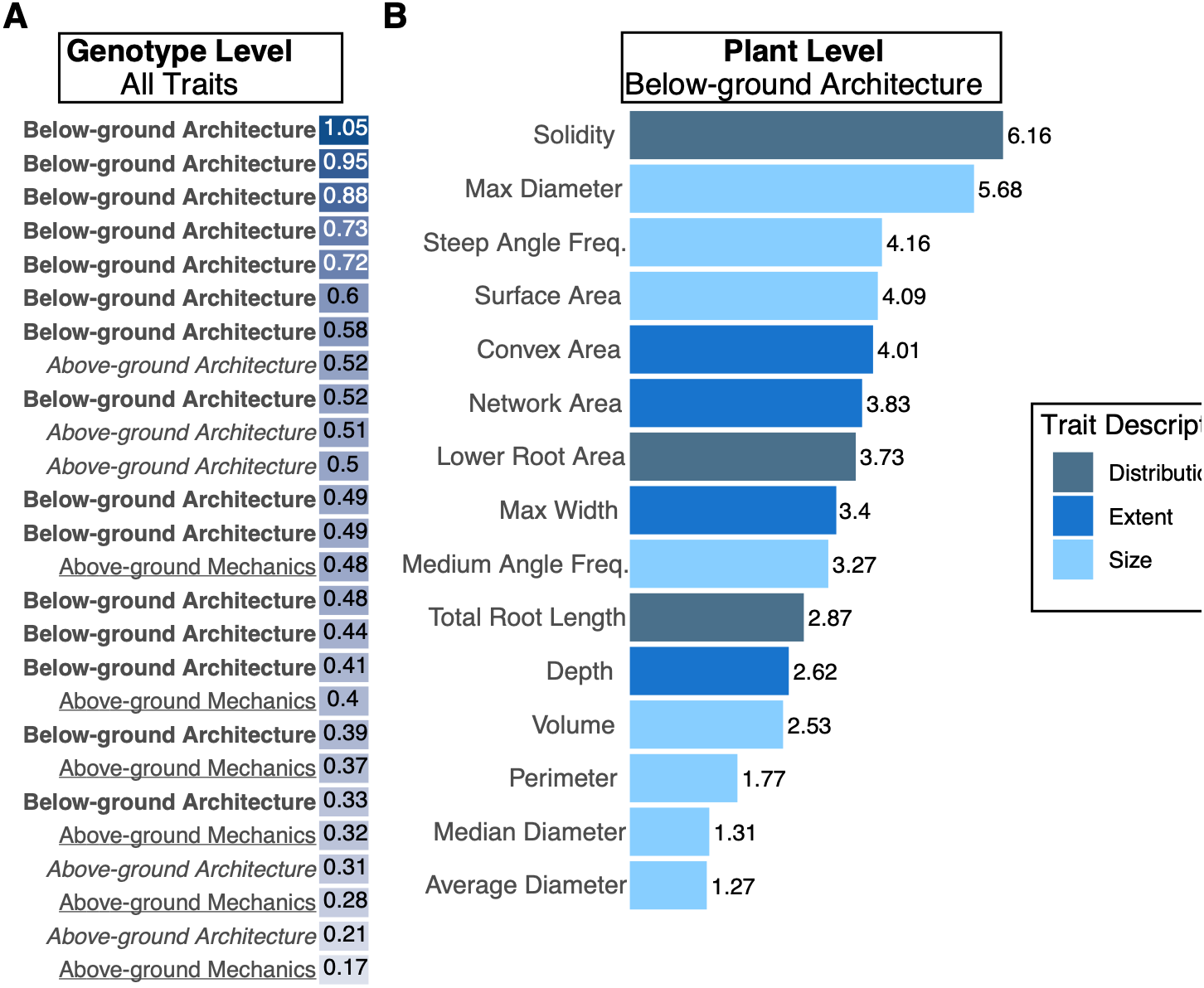
The size and distribution of the below-ground root system best predicts root system stiffness categories. (A) A random forest model was used to predict the root system stiffness category from genotypic averages of below-ground traits, above-ground traits, and above-ground mechanics. Predictor importance was determined by the mean decrease in Gini value. A darker shade of blue indicates that the predictor had a higher mean decrease in Gini value and a lighter shade of blue indicates that the predictor had a lower mean decrease in Gini value. The top 25% of predictors were all below-ground root architecture traits (mean decrease in Gini value 0.58-1.05). (B) A random forest model was used to predict the root system stiffness category from individual plant data for below-ground traits. Predictor importance was determined by the mean decrease in Gini value (shown to the right of the bar). Colored bars describe the below-ground trait description.

The multi-year data required the use of genotypic averages, which limited the power of a predictive model. Therefore, a second random forest model was developed to further examine the relationship between the below-ground root architecture and root system stiffness categories. This model predicted the root system stiffness category with 70% accuracy. A mean decrease in Gini value was used to assess the relative importance of predictors. Traits with the highest variable importance included those related to root system size (maximum crown diameter, steep angle frequency, surface area) and spatial distribution (solidity)(**Figure 6B**). Many of these traits are correlated, thus their individual importance should not be interpreted as evidence of independent effects on root system stiffness. Rather, the model identified a suite of below-ground root architecture traits describing the size and distribution of the root system that collectively predicted root system stiffness.

### Root hairs alter root system stiffness through modulating root system size

The inbred studies focused on quantifying the physical aspects of the root system that contribute to root system stiffness. However, equally important is the root-soil adhesion of the rhizosheath, which is driven by root hairs (Burak et al., 2021). To understand the role of the rhizosheath, the *roothairless3* (*rth3*) mutant (Hochholdinger et al., 2008) was analyzed for changes in root system stiffness. A comparison of the *rth3* mutants with wild-type sibling showed that the root system stiffness was significantly less for *rth3* mutants (**Figure 7A, Table S15**). However, the maximum root system width was also reduced for *rth3* mutants (**Figure 7B, Table S16**). To determine whether a reduction in root system stiffness for *rth3* mutants was solely due to the reduction in root system size, *rth3* data was scaled for comparison to the 43 inbred genotypes. This showed that both the wild-type sibling and the *rth3* mutant plants are within the expected range of the root system stiffness given the root system size (**Figure 7C**). Although the absence of root hairs may contribute to the reduced stiffness of *rth3*, the observed reduction in root system stiffness was consistent with differences in root system size.

**Figure 7.**
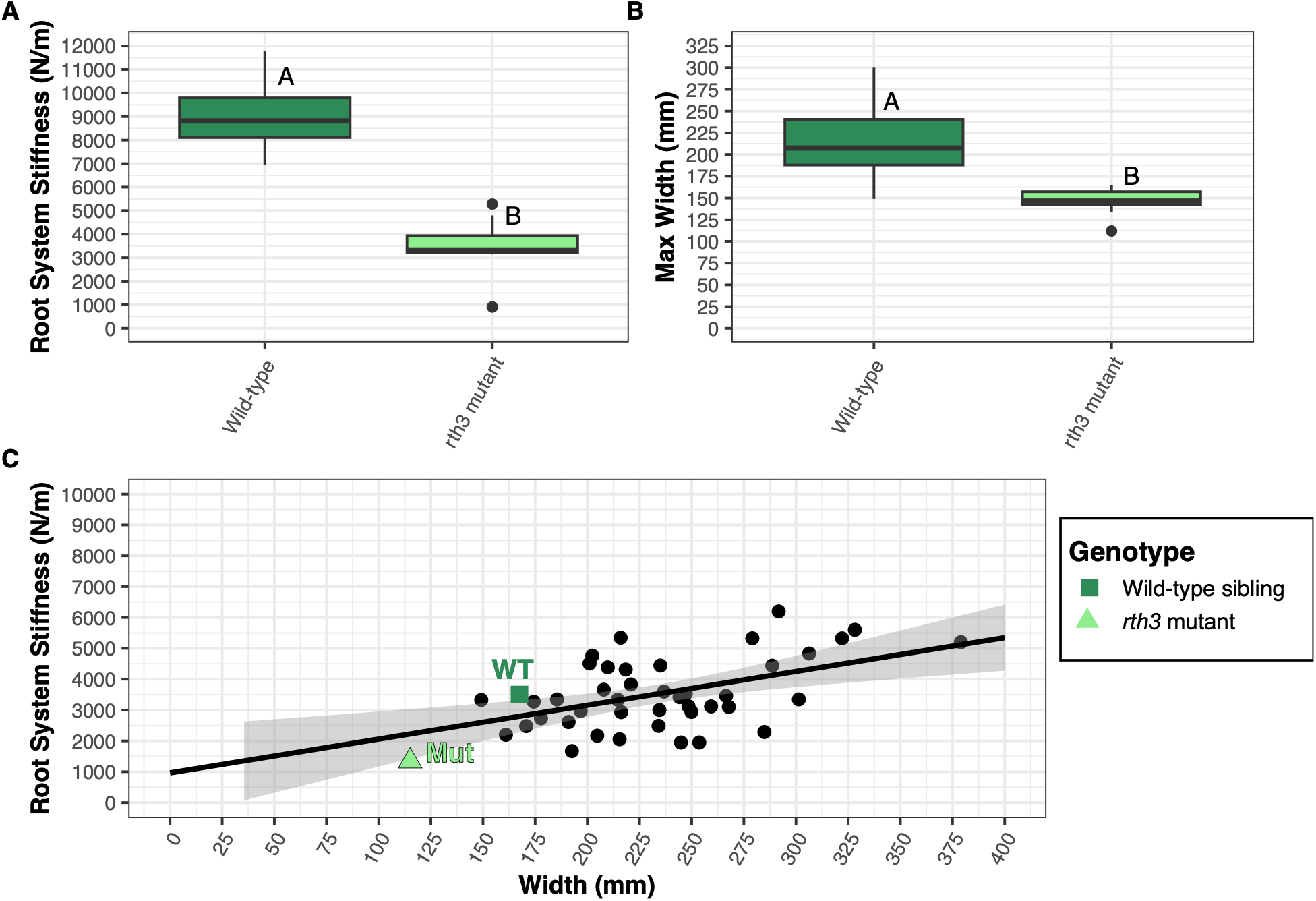
The *rth3* mutant had reduced root system stiffness due to a reduction in root system size. (A) The *roothairless3* (*rth3*) mutant had a lower root system stiffness compared to its wild-type sibling. (B) The *rth3* mutant had a smaller maximum crown width compared to its wild-type sibling. Genotypes that share a letter are not significantly different from one another (Tukey HSD assessed at *P* < 0.05). Black dots indicate outliers. (C) When comparing the *rth3* mutant plant and its wild-type sibling to the 43 inbred genotypes assessed in this study, both the wild-type sibling and *rth3* mutant plant are within the expected range for size (maximum crown width) and function (root system stiffness) compared to the rest of the inbred genotypes.

## DISCUSSION

While root system stiffness provides an indication of root lodging susceptibility, the relationship between root system stiffness and root lodging susceptibility is non-linear. Consistent with previous observations in maize hybrids (Hostetler et al., 2025), inbred genotypes with a higher root system stiffness are associated greater root lodging susceptibility. Yet, this observation does not hold at the individual genotype-level, indicating that root system stiffness is only one component of root lodging susceptibility. Indeed, root lodging occurs when the external bending moment, imposed by the stalk, overcomes the resistance of the root system to rotation. The external bending moment is, in part, influenced by plant height (Stubbs et al., 2023). In our data, LH252 experienced severe root lodging despite being categorized as having a low root system stiffness. When considering plant height, LH252 was among the tallest genotypes (Hostetler et al., 2022b). On the other side, inbred genotypes with a high root system stiffness and low root lodging susceptibility are among the shortest genotypes. Neither root system stiffness nor height can act as stand-alone indicators of root lodging susceptibility. This is supported by a recent publication that evaluated the root system stiffness of short-stature maize and showed that short stature plants with a high root system stiffness still experienced root lodging (Evers et al., 2026). Together these data show that plant height and root system stiffness need to be considered together when targeting root lodging resistance.

This study aimed to understand plant traits underlying root system stiffness. In a small diversity panel of 43 maize inbred genotypes, the below-ground root architecture, above-ground root architecture, and above-ground root mechanics were characterized. For all traits, larger values were consistently associated with stiffer root systems. When considering all traits, random forest modeling showed that the below-ground root traits were the top predictors of root system stiffness categories. Specifically, the most important predictors were traits describing the root system distribution and size, including solidity, maximum diameter, surface area, and the frequency of steep root angles. As with most phenotyping, many traits are highly correlated. Thus, it is impossible to target individual traits, and there is a need to understand the root system more holistically. These data support the proposition that root system stiffness provides a compound measure of the size and extent of the root system, as opposed to individual root traits. The association of larger root systems with higher stiffness is intuitive as larger root systems would require more force to move the same distance, reflected as a higher root system stiffness.

While the size and distribution of the root system are the top predictors of root system stiffness, the mechanics of individual roots is also likely important. We characterized the mechanical properties of individual above-ground roots, but the extent to which these properties reflect those of subterranean roots remains unknown. There are multiple subterranean root types (e.g., crown, seminal, and primary), which likely all have different mechanical properties. We’ve previously shown that even within a root-type (i.e., brace roots), there is variation based on the node of origin (Hostetler et al., 2022a). At present, it is difficult to isolate individual, intact subterranean roots for mechanical testing, especially at the scale and root-type resolution required for this study. Future studies using large-scale rhizoboxes (Henrickson et al., 2026) could help disentangle the contributions of individual root types and their mechanical properties to the overall root system stiffness. However, this approach does not fully replicate field conditions, and the SMURF device has not yet been benchmarked for indoor use.

Soil properties, including texture, moisture, and compaction, may influence both root development and the interaction between the roots and soil (Bengough et al., 2011; Bengough et al., 2016). In this study, we evaluated root system stiffness within a common environment and did not assess the impact of various soil properties. Previous work from our lab showed that the magnitude of root system stiffness changed between soil types and environments, but the relative relationship between genotypes remained constant (Evers et al., 2026). This was only assessed for a limited number of lines and locations, and the role of soil in establishing root system stiffness remains to be fully elucidated.

To analyze the role of root-soil adhesion, we quantified the root system stiffness of *rth3* mutants, which have a reduced rhizosheath. Although *rth3* mutants exhibited a reduced root system stiffness compared to their wild-type siblings, they also had smaller root systems. The reduced root system stiffness values were within expectations of the root system size. Thus, we could not separate an effect of the rhizosheath from the effect of root system size in the *rth3* mutants. The *roothairless1* mutant has a much more severe reduction in root hairs (Wen et al., 2005; Bray and Topp, 2018), however this mutant did not survive in our field conditions. The current device design limits its use to large and mature maize plants. Future approaches that can accommodate smaller and younger plants could help resolve the role of the rhizosheath in root system stiffness and enable characterization of the root system stiffness throughout plant development.

There is substantial opportunity to develop new technologies aimed and quantifying root mechanics that would enable a better understanding of the root system stiffness component of root lodging resistance. Currently, no field-based *in situ* approaches are available to monitor or assess the mechanical properties of individual subterranean root mechanics. Although the device used in this study enables measurements of the overall root system stiffness, it is important to note that it is in its infancy and several important questions remain. These include how root system stiffness changes throughout plant development, the genetic factors underlying the variation in root system stiffness, and the influence of environmental conditions on measurements. As root lodging continues to be a challenge for maize as well as other crops, innovative solutions to develop fail-safe breeding targets are needed to secure future yields.

Collectively, our findings demonstrate that below-ground root system size and spatial distribution are important predictors of root system stiffness in maize. The ability to quantify stiffness rapidly and non-destructively provides opportunities to investigate these relationships across larger populations and across time. Findings here highlight that the root system stiffness may be used as a proxy of below-ground root system architecture and even lodging resistance if other plant aspects are considered (e.g., external bending moment).

## SUPPLEMENTARY DATA

**Figure S1.** Root system stiffness was highly correlated and inbred genotype rankings were consistent across years.

**Figure S2.** Below-ground root architectural traits varied among maize inbred genotypes.

**Figure S3.** Principal component analysis showed that high root system stiffness was associated with larger and more extensive below-ground root systems.

**Figure S4.** Several below-ground root architectural traits did not significantly differ among root system stiffness categories.

**Figure S5.** Above-ground root architectural traits varied among maize inbred genotypes.

**Figure S6.** Principal component analysis showed that high root system stiffness was associated with larger above-ground root systems and greater stalk width.

**Figure S7.** The number of brace root nodes entering the soil did not significantly differ among root system stiffness categories.

**Figure S8.** Brace root emergence date varied among inbred genotypes and was influenced by genotype, year, and their interaction.

**Figure S9.** Brace root mechanical traits varied among inbred genotypes and brace root nodes.

**Figure S10.** Brace root mechanical traits varied among brace root nodes.

**Figure S11.** Plant height varied among genotypes with contrasting combinations of root system stiffness and root lodging susceptibility.

**Table S1.** List of the 43 maize inbred genotypes used in this study and their subpopulation information.

**Table S2.** Summary statistics and genotypic differences in root system stiffness among maize inbred genotypes assessed in 2020.

**Table S3.** Summary statistics and genotypic differences in root system stiffness among maize inbred genotypes assessed in 2021.

**Table S4.** Summary statistics and differences in root system stiffness between years.

**Table S5.** Summary statistics and differences in root system stiffness among root lodging susceptibility categories.

**Table S6.** Summary statistics and genotypic differences in below-ground root architectural traits.

**Table S7.** Summary statistics and differences in below-ground root architectural traits among root system stiffness categories.

**Table S8.** Summary statistics and differences in above-ground root architectural traits among root system stiffness categories.

**Table S9.** Summary statistics and genotypic differences in brace root emergence date across years.

**Table S10.** Summary statistics and differences in brace root emergence among root system stiffness categories.

**Table S11.** Summary statistics for genotypic and node-specific differences in brace root mechanical traits.

**Table S12.** Summary statistics and differences in brace root mechanical traits among root system stiffness categories.

**Table S13.** Summary statistics and node-specific differences in brace root mechanical traits.

**Table S14.** Input data used to train and test random forest classification models predicting root system stiffness categories.

**Table S15.** Summary statistics and differences in root system stiffness between *roothairless3* (*rth3*) mutants and wild-type siblings.

**Table S16.** Summary statistics and differences in maximum root crown width between *roothairless3* (*rth3*) mutants and wild-type siblings.

## ACKNOWLEDGEMENTS

We gratefully acknowledge Justin Sapp and Dr. Joey Norikane for image acquisition of excavated root crowns, Sparks lab members who assisted in root digs and cutting roots, Dr. Jim Holland for providing original seed stocks, Irene Ikiriko for helping measure roots for 3-point bending, Joe Cristiano for assistance with root painter, Dr. Larry York for helpful discussions on Rhizovision Explorer, Dr. Ian Dodd for suggesting the rhizosheath experiment, Dr. Jingjing Tong for assisting with the *roothairless3* (*rth3*) mutant experiment, and Dr. Philip Benfey for providing resources for this work to be initiated in his lab.

## CRediT

Ashley N. Hostetler - Data Curation, Formal Analysis, Visualization, Writing - original draft, Writing - review & editing

Emilia Pierce - Formal Analysis, Investigation, Writing - original draft, Writing - review & editing

Jonathan W. Reneau - Conceptualization, Investigation, Methodology, Writing - review & editing

David Griffin - Investigation, Writing - review & editing

Erin E. Sparks - Conceptualization, Formal Analysis, Funding Acquisition, Investigation, Methodology, Project Administration, Supervision, Writing - original draft, Writing - review & editing

## CONFLICT OF INTEREST

EES and JWR are co-founders of Izbe Innovations LLC, an engineering design and prototyping company. The company was not involved in the design, conduct, analysis, or funding of this work.

## FUNDING

This work was partially supported by NSF CMMI award #2040346 to EES, and a United States Department of Agriculture (USDA) National Institute of Food and Agriculture (NIFA) Postdoctoral Fellowship awarded to ANH (#2022-67012-36840).

## DATA AVAILABILITY

All raw data, processed data, and scripts used to analyze data are available on GitHub: https://github.com/EESparksLab/Hostetler_Pierce_et_al_2026

## Abbreviations

SMURF: Sorghum and Maize Under Rotational Force
dap: days after planting
analysis of variance: ANOVA
PCA: Principal Component Analysis
PC: Principal Component
RSS: Root System Stiffness

## Notes

https://github.com/EESparksLab/Hostetler_Pierce_et_al_2026

### Summary of Updates

There were updates made to the discussion.

