## Supplemental Figure for "Maize root system stiffness is determined by the size and distribution of the below-ground root system"

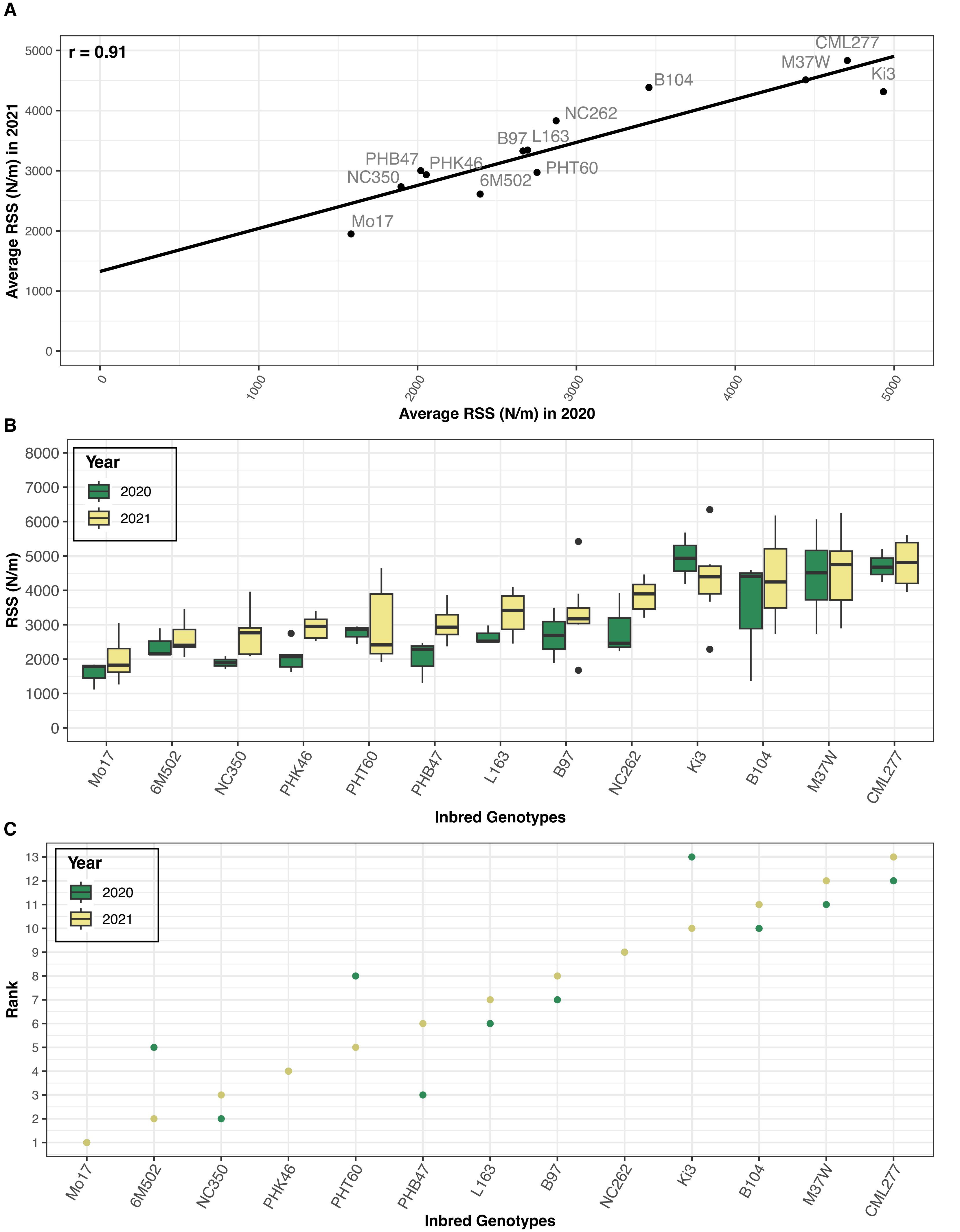


Figure S1. The root system stiffness was consistent across years. (A) The root system stiffness was highly correlated between years (r=0.91). (B) There was an inbred genotype and year effect, but not interaction effect. In 2020, the root system stiffness was on average less than it was in 2021 regardless of genotype. (C) Inbred genotypes were ranked from lowest (1) to highest (13) root system stiffness within each year. Comparing rankings between years showed that inbred genotypes ranked similarly across years.

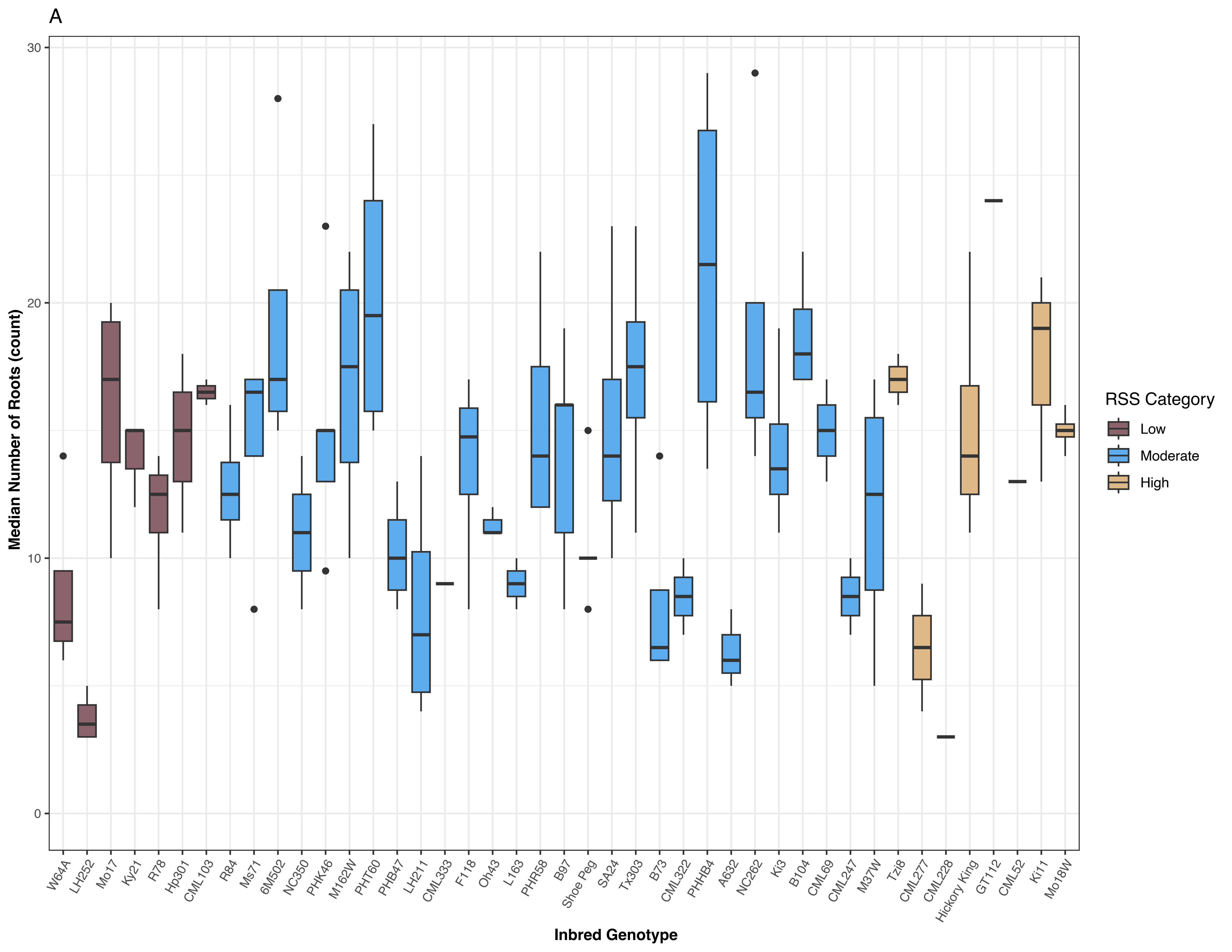
A
30
20
10
0
Median Number of Roots (count)
W64A
LH252
Mo17
Ky21
R78
Hp301
CML103
R84
Ms71
6M502
NC350
PHK46
M162W
PHT60
PHB47
LH211
CML333
F118
Oh43
L163
PHR58
B97
Shoe Peg
SA24
Tx303
B73
CML322
PHHB4
A632
NC262
Ki3
B104
CML69
CML247
M37W
Tzi8
CML277
CML228
Hickory King
GT112
CML52
Ki11
Mo18W
Inbred Genotype
RSS Category
Low
Moderate
High

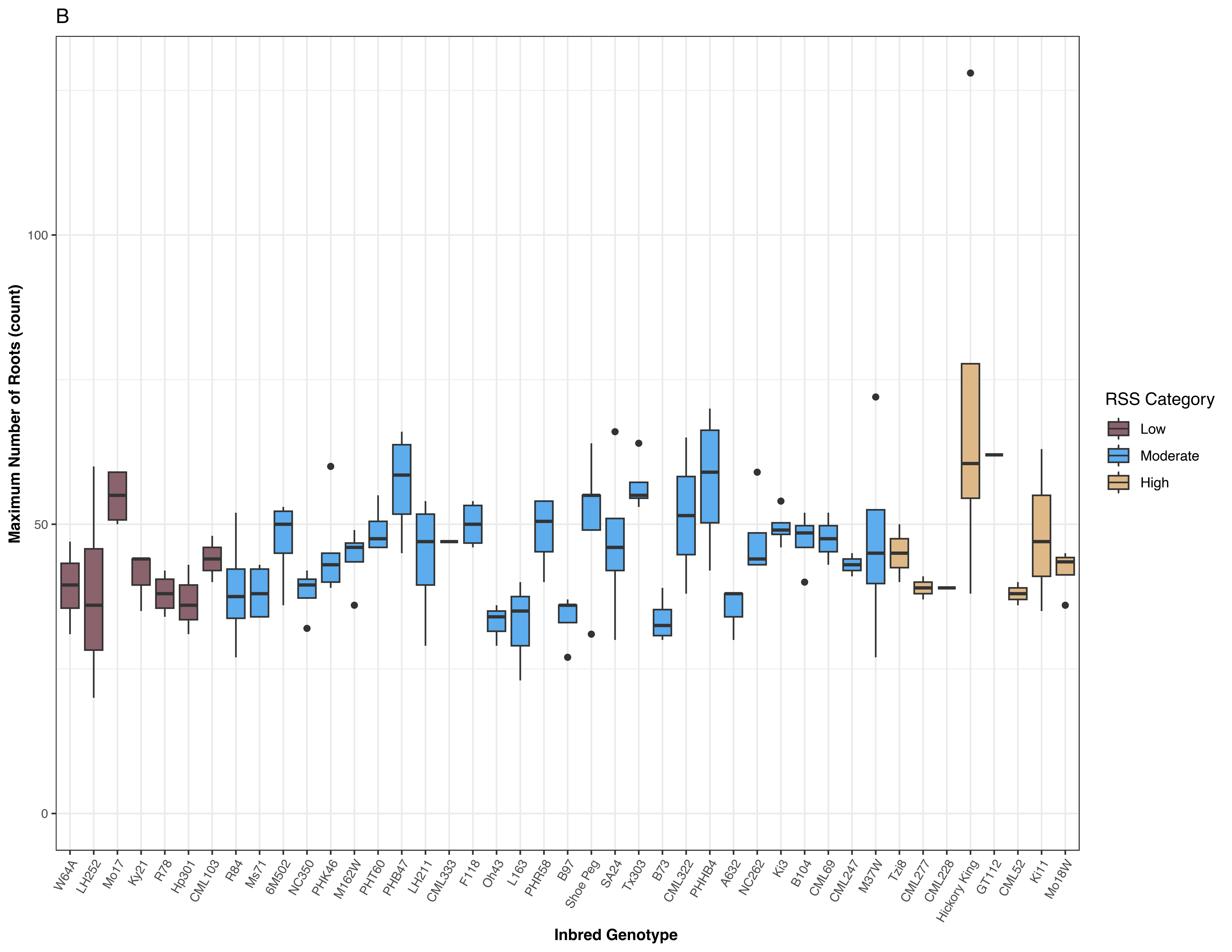

B
Maximum Number of Roots (count)
0
50
100
Inbred Genotype
W64A
LH252
Mo17
Ky21
R78
Hp301
CML103
R84
Ms71
6M502
NC350
PHK46
M162W
PHT60
PHB47
LH211
CML333
F118
Oh43
L163
PHR58
B97
Shoe Peg
SA24
Tx303
B73
CML322
PHHB4
A632
NC262
Ki3
B104
CML69
CML247
M37W
Tzi8
CML277
CML228
Hickory King
GT112
CML52
Ki11
Mo18W
RSS Category
Low
Moderate
High

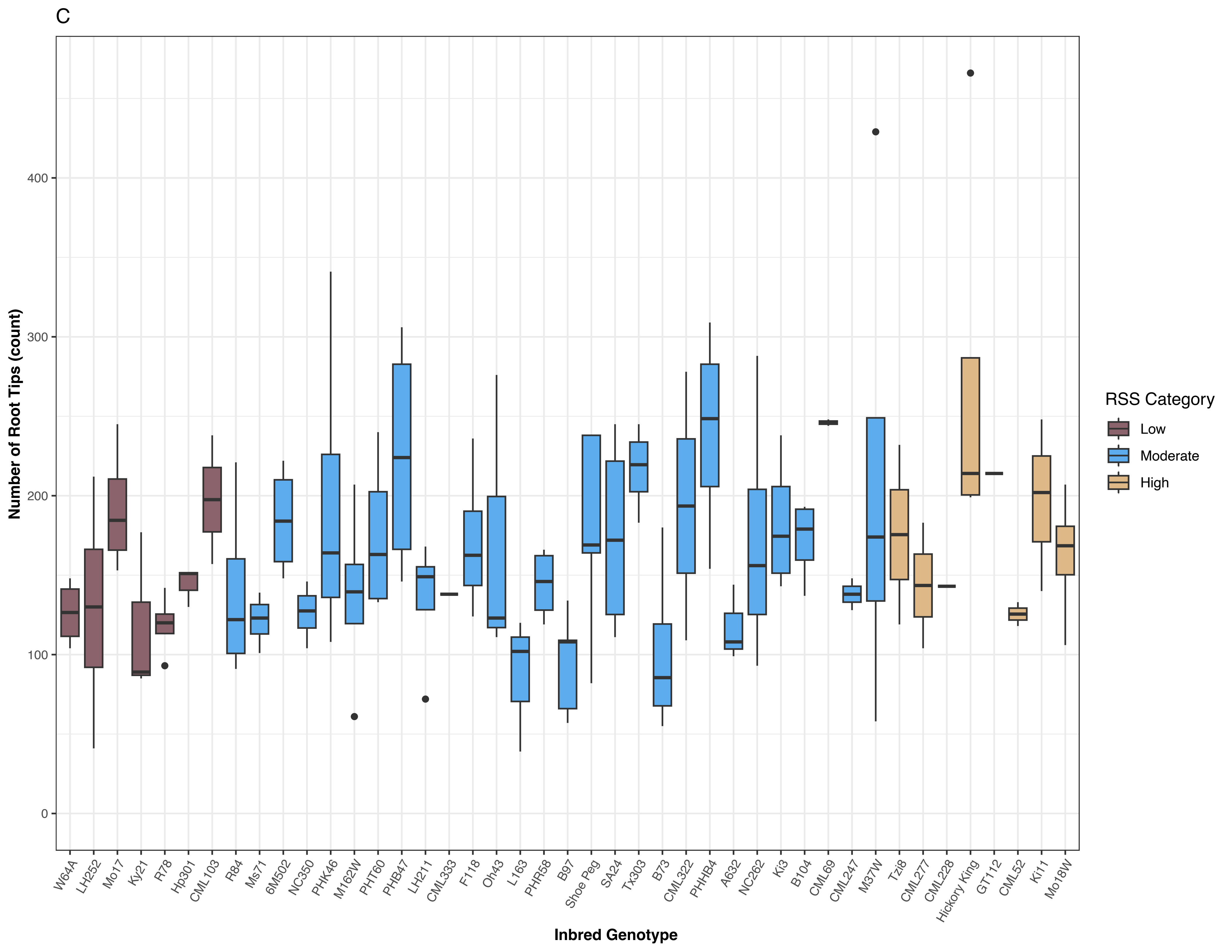
C
Number of Root Tips (count)
0
100
200
300
400
Inbred Genotype
W64A
LH252
Mo17
Ky21
R78
Hp301
CML103
R84
Ms71
6M502
NC350
PHK46
M162W
PHT60
PHB47
LH211
CML333
F118
Oh43
L163
PHR58
B97
Shoe Peg
SA24
Tx303
B73
CML322
PHHB4
A632
NC262
Ki3
B104
CML69
CML247
M37W
Tzi8
CML277
CML228
Hickory King
GT112
CML52
Ki11
Mo18W
RSS Category
Low
Moderate
High

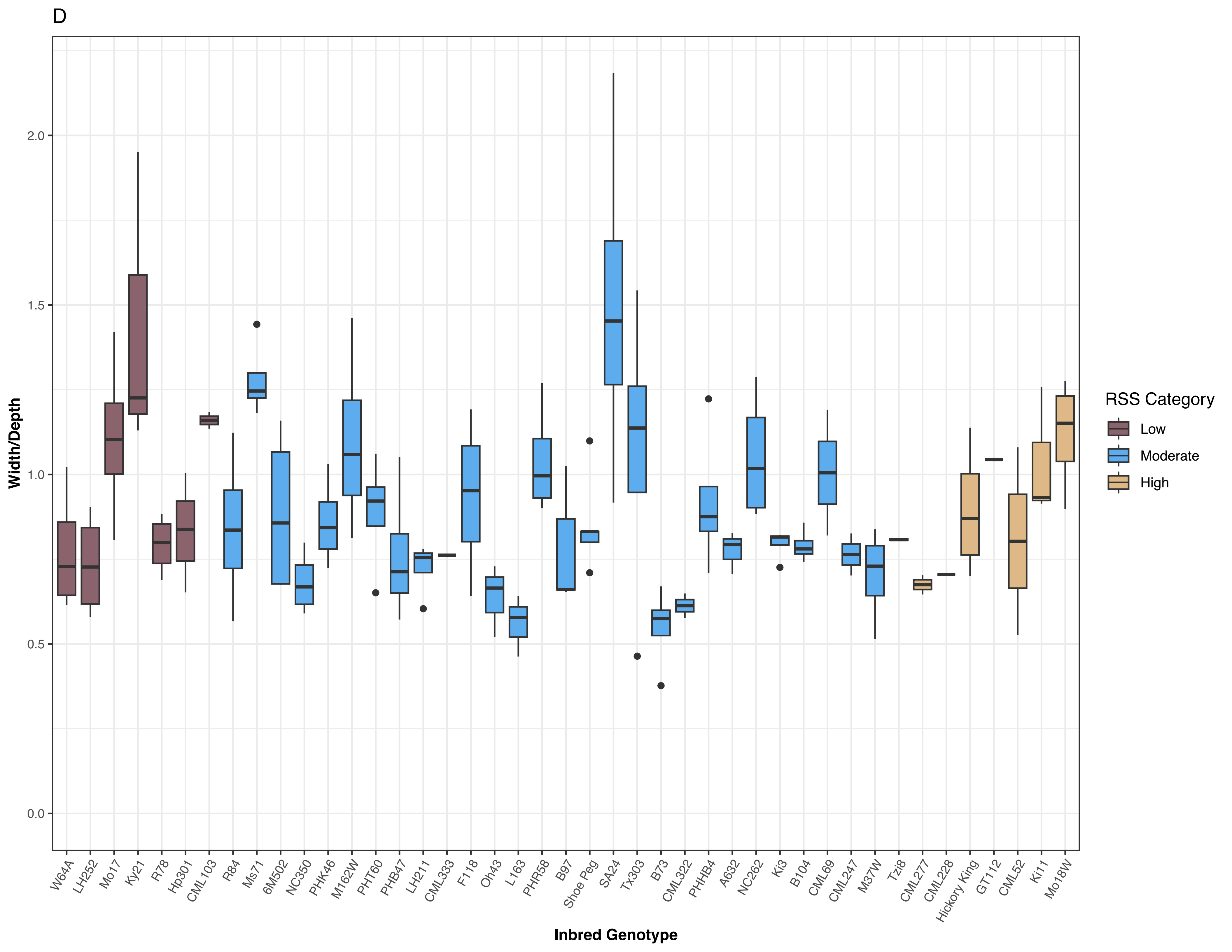

D
Width/Depth
2.0
1.5
1.0
0.5
0.0
Inbred Genotype
W64A
LH252
Mo17
Ky21
R78
Hp301
CML103
R84
Ms71
6M502
NC350
PHK46
M162W
PHT60
PHB47
LH211
CML333
F118
Oh43
L163
PHR58
B97
Shoe Peg
SA24
Tx303
B73
CML322
PHHB4
A632
NC262
Ki3
B104
CML69
CML247
M37W
Tzi8
CML277
CML228
Hickory King
GT112
CML52
Ki11
Mo18W
RSS Category
Low
Moderate
High

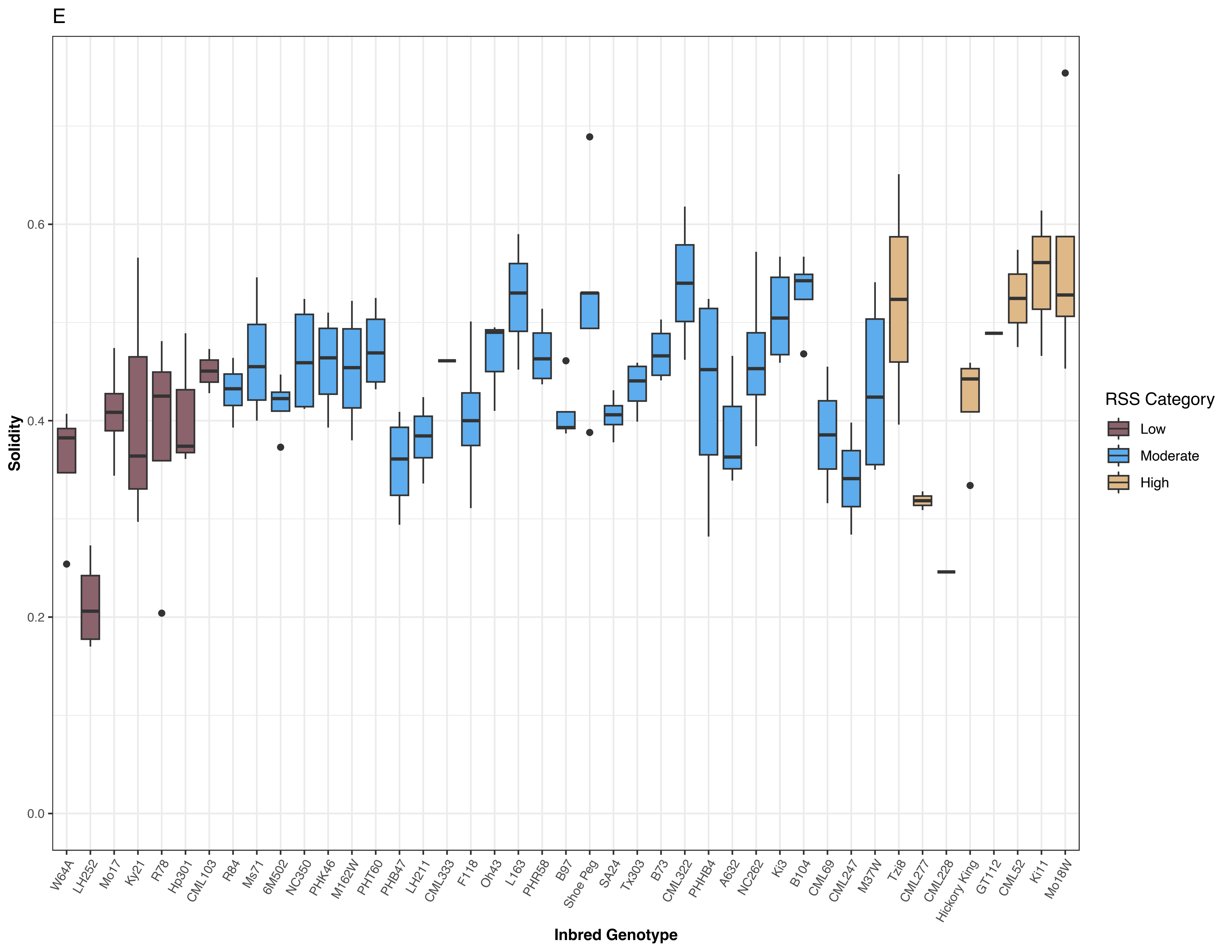
E
Solidity
0.0
0.2
0.4
0.6
Inbred Genotype
W64A
LH252
Mo17
Ky21
R78
Hp301
CML103
R84
Ms71
6M502
NC350
PHK46
M162W
PHT60
PHB47
LH211
CML333
F118
Oh43
L163
PHR58
B97
Shoe Peg
SA24
Tx303
B73
CML322
PHHB4
A632
NC262
Ki3
B104
CML69
CML247
M37W
Tzi8
CML277
CML228
Hickory King
GT112
CML52
Ki11
Mo18W
RSS Category
Low
Moderate
High

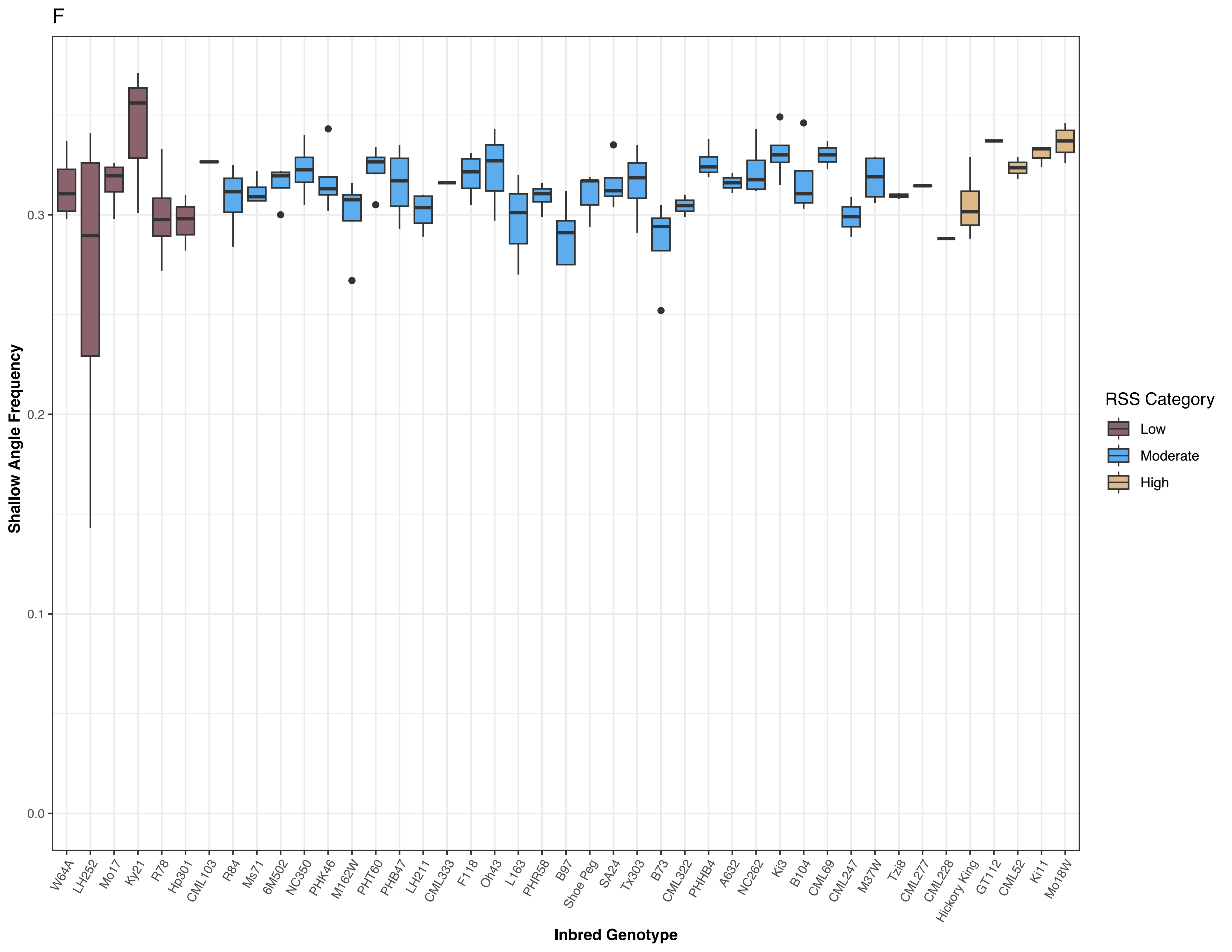
F
Shallow Angle Frequency
0.0
0.1
0.2
0.3
Inbred Genotype
W64A
LH252
Mo17
Ky21
R78
Hp301
CML103
R84
Ms71
6M502
NC350
PHK46
M162W
PHT60
PHB47
LH211
CML333
F118
Oh43
L163
PHR58
B97
Shoe Peg
SA24
Tx303
B73
CML322
PHHB4
A632
NC262
Ki3
B104
CML69
CML247
M37W
Tzi8
CML277
CML228
Hickory King
GT112
CML52
Ki11
Mo18W
RSS Category
Low
Moderate
High

G

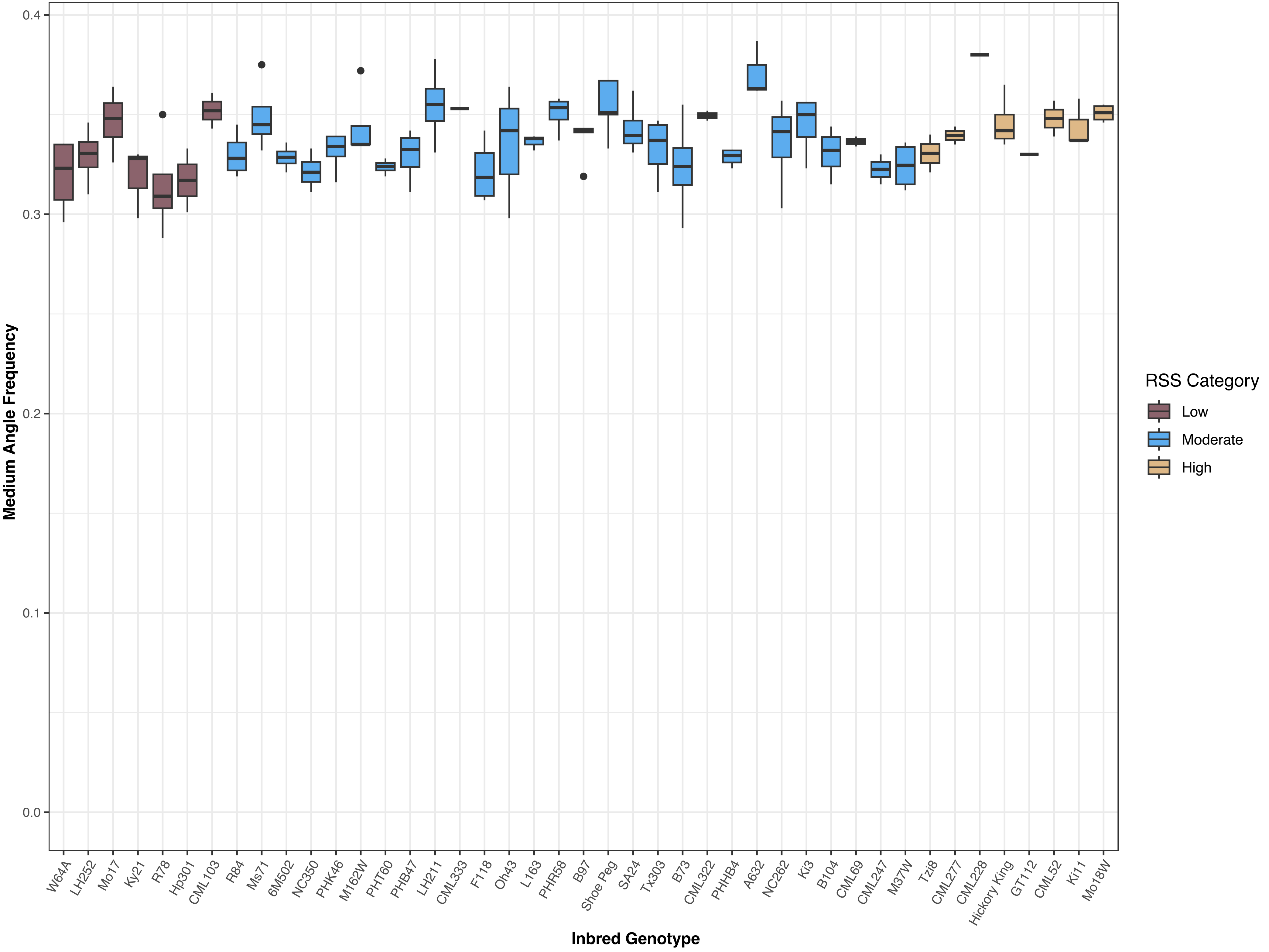

Medium Angle Frequency
0.4
0.3
0.2
0.1
0.0
Inbred Genotype
W64A
LH252
Mo17
Ky21
R78
Hp301
CML103
R84
Ms71
6M502
NC350
PHK46
M162W
PHT60
PHB47
LH211
CML333
F118
Oh43
L163
PHR58
B97
Shoe Peg
SA24
Tx303
B73
CML322
PHHB4
A632
NC262
Ki3
B104
CML69
CML247
M37W
Tzi8
CML277
CML228
Hickory King
GT112
CML52
Ki11
Mo18W
RSS Category
Low
Moderate
High

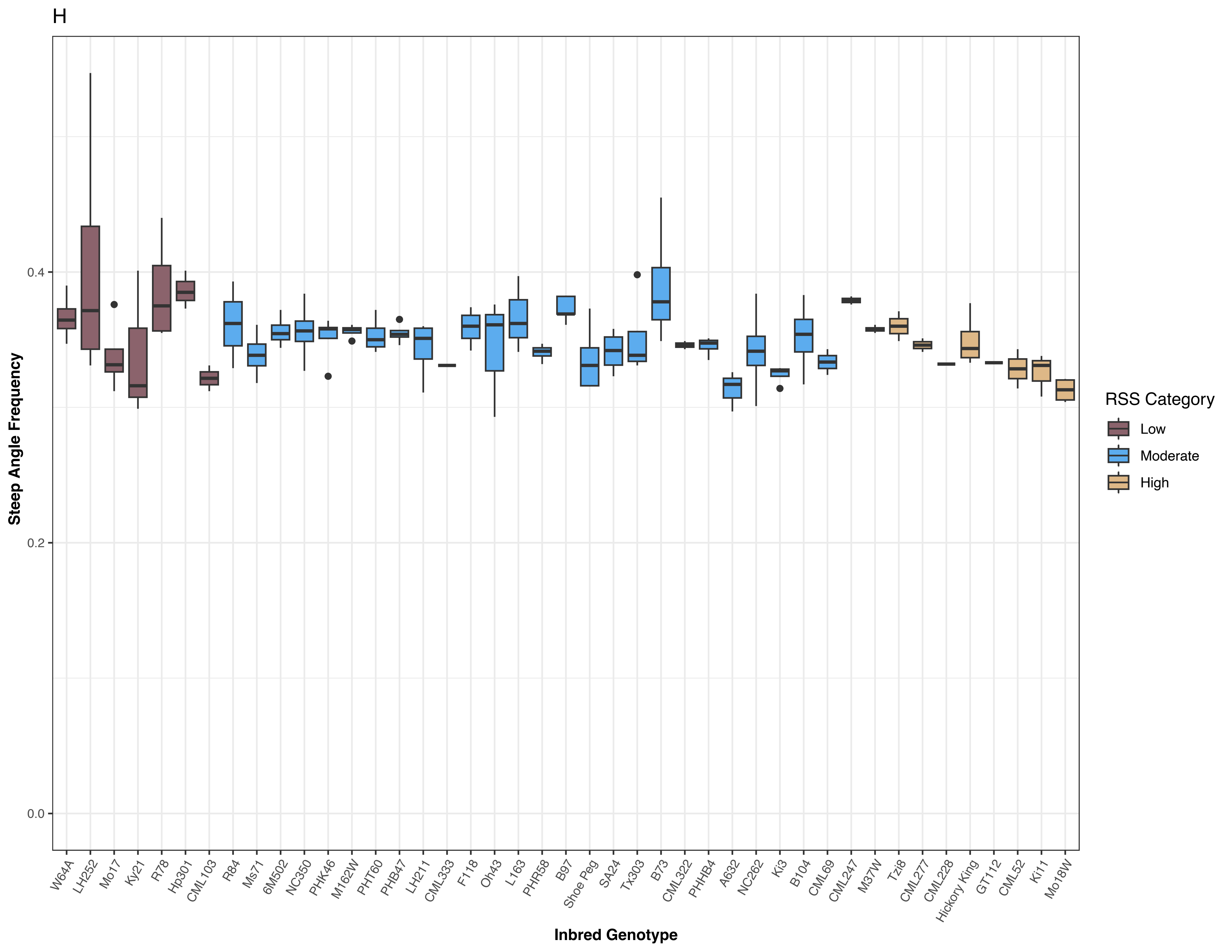

H
Steep Angle Frequency
0.0
0.2
0.4
Inbred Genotype
W64A
LH252
Mo17
Ky21
R78
Hp301
CML103
R84
Ms71
6M502
NC350
PHK46
M162W
PHT60
PHB47
LH211
CML333
F118
Oh43
L163
PHR58
B97
Shoe Peg
SA24
Tx303
B73
CML322
PHHB4
A632
NC262
Ki3
B104
CML69
CML247
M37W
Tzi8
CML277
CML228
Hickory King
GT112
CML52
Ki11
Mo18W
RSS Category
Low
Moderate
High

I

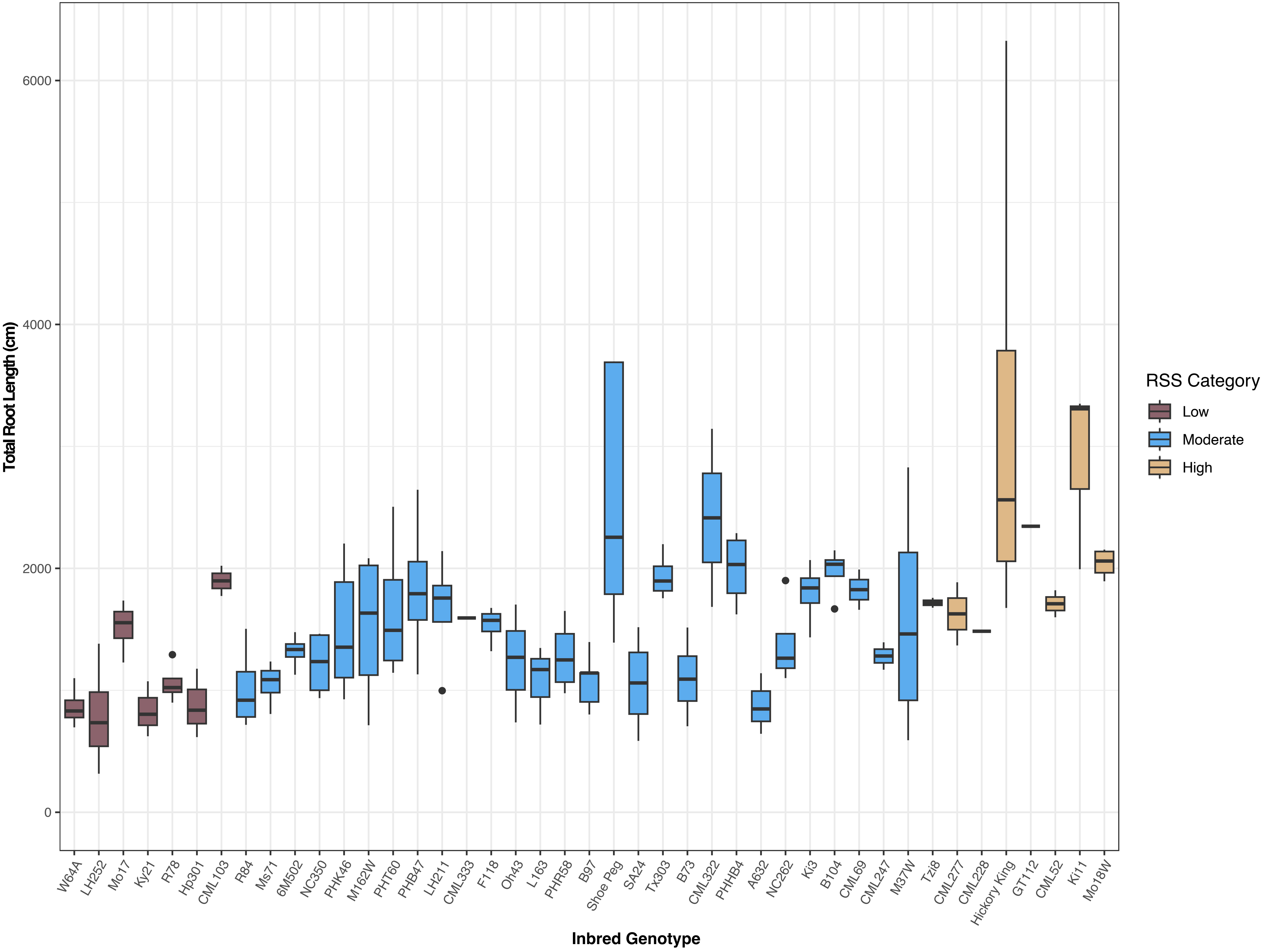

Total Root Length (cm)
6000
4000
2000
0
Inbred Genotype
W64A
LH252
Mo17
Ky21
R78
Hp301
CML103
R84
Ms71
6M502
NC350
PHK46
M162W
PHT60
PHB47
LH211
CML333
F118
Oh43
L163
PHR58
B97
Shoe Peg
SA24
Tx303
B73
CML322
PHHB4
A632
NC262
Ki3
B104
CML69
CML247
M37W
Tzi8
CML277
CML228
Hickory King
GT112
CML52
Ki11
Mo18W
RSS Category
Low
Moderate
High

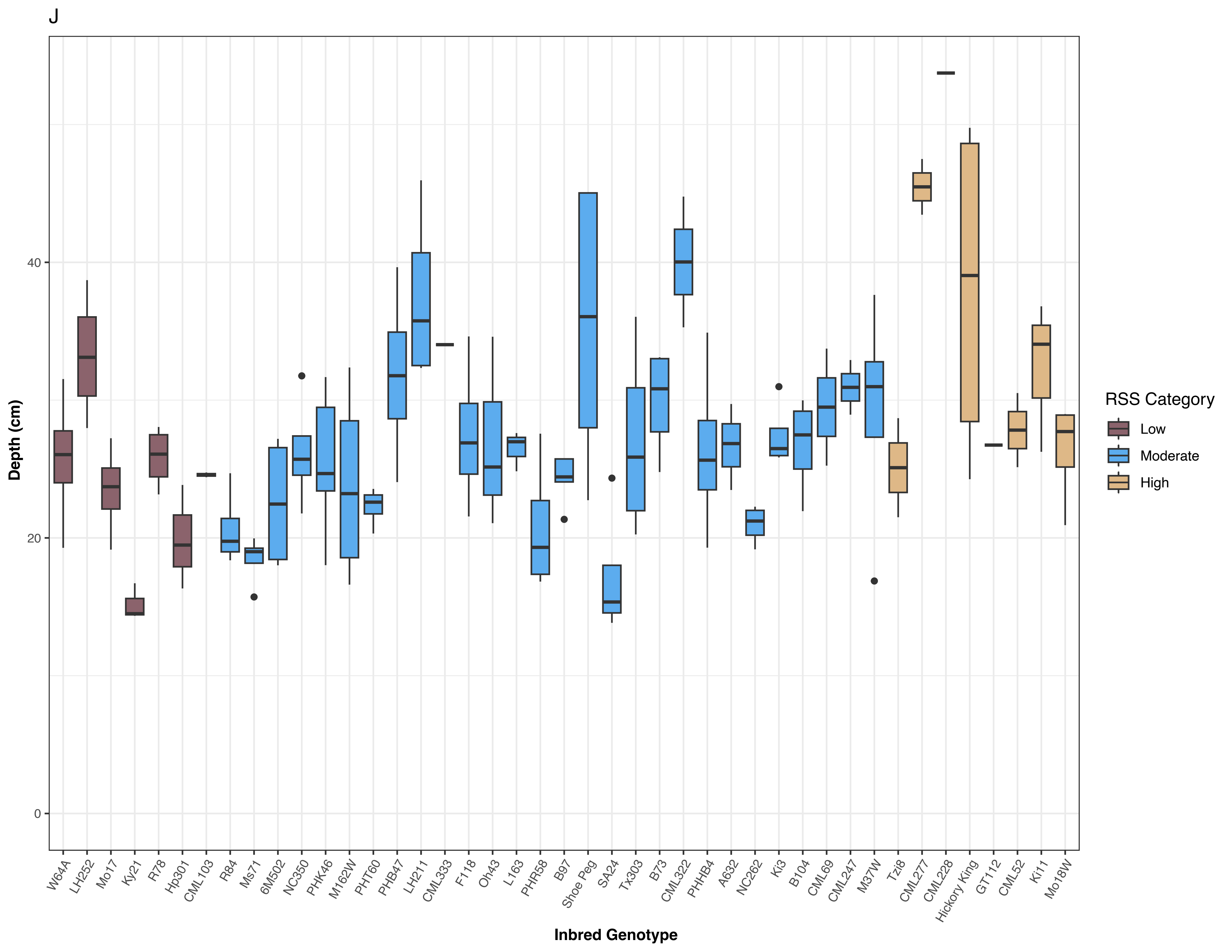
J
Depth (cm)
40
20
0
W64A
LH252
Mo17
Ky21
R78
Hp301
CML103
R84
Ms71
6M502
NC350
PHK46
M162W
PHT60
PHB47
LH211
CML333
F118
Oh43
L163
PHR58
B97
Shoe Peg
SA24
Tx303
B73
CML322
PHHB4
A632
NC262
Ki3
B104
CML69
CML247
M37W
Tzi8
CML277
CML228
Hickory King
GT112
CML52
Ki11
Mo18W
Inbred Genotype
RSS Category
Low
Moderate
High

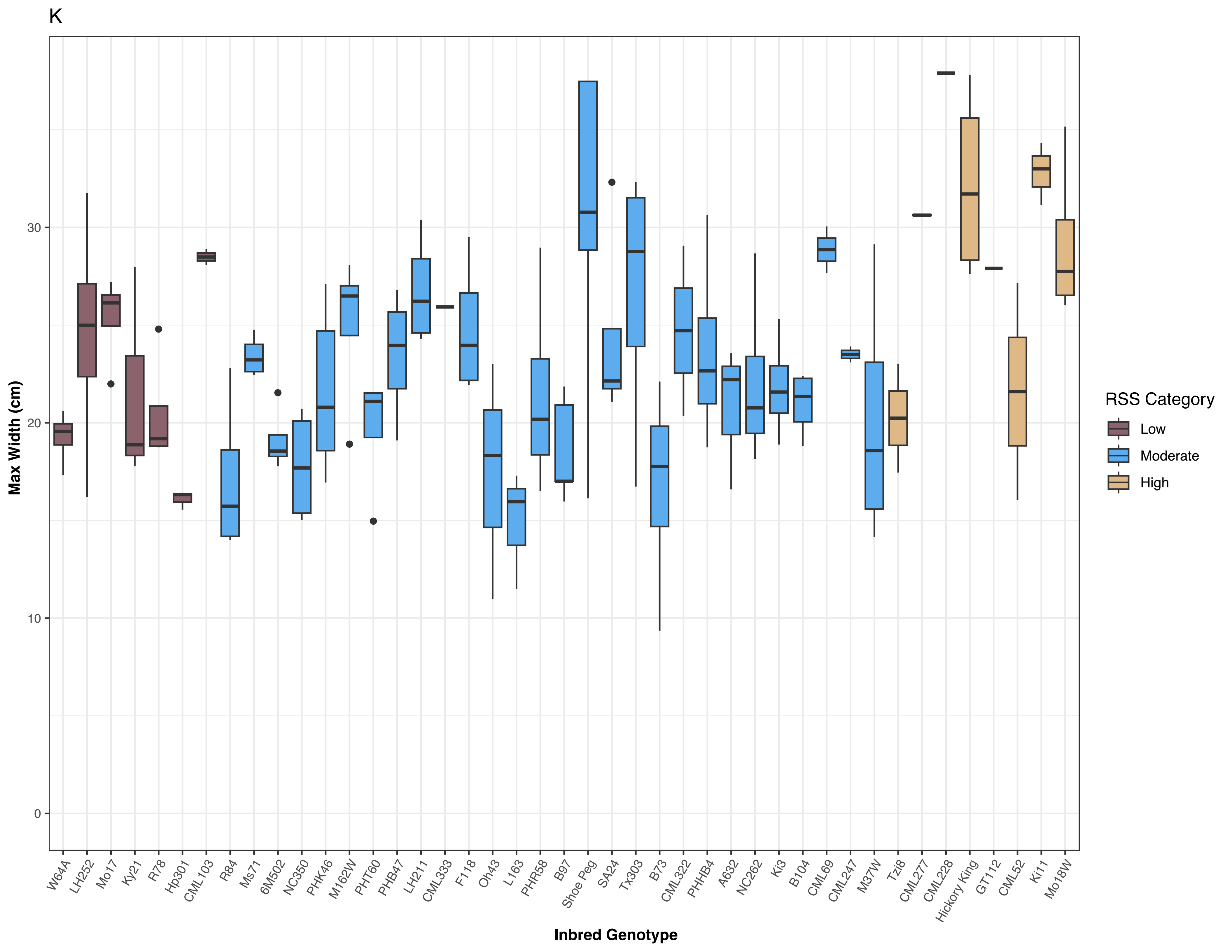
K
Max Width (cm)
0
10
20
30
W64A
LH252
Mo17
Ky21
R78
Hp301
CML103
R84
Ms71
6M502
NC350
PHK46
M162W
PHT60
PHB47
LH211
CML333
F118
Oh43
L163
PHR58
B97
Shoe Peg
SA24
Tx303
B73
CML322
PHHB4
A632
NC262
Ki3
B104
CML69
CML247
M37W
Tzi8
CML277
CML228
Hickory King
GT112
CML52
Ki11
Mo18W
Inbred Genotype
RSS Category
Low
Moderate
High

L

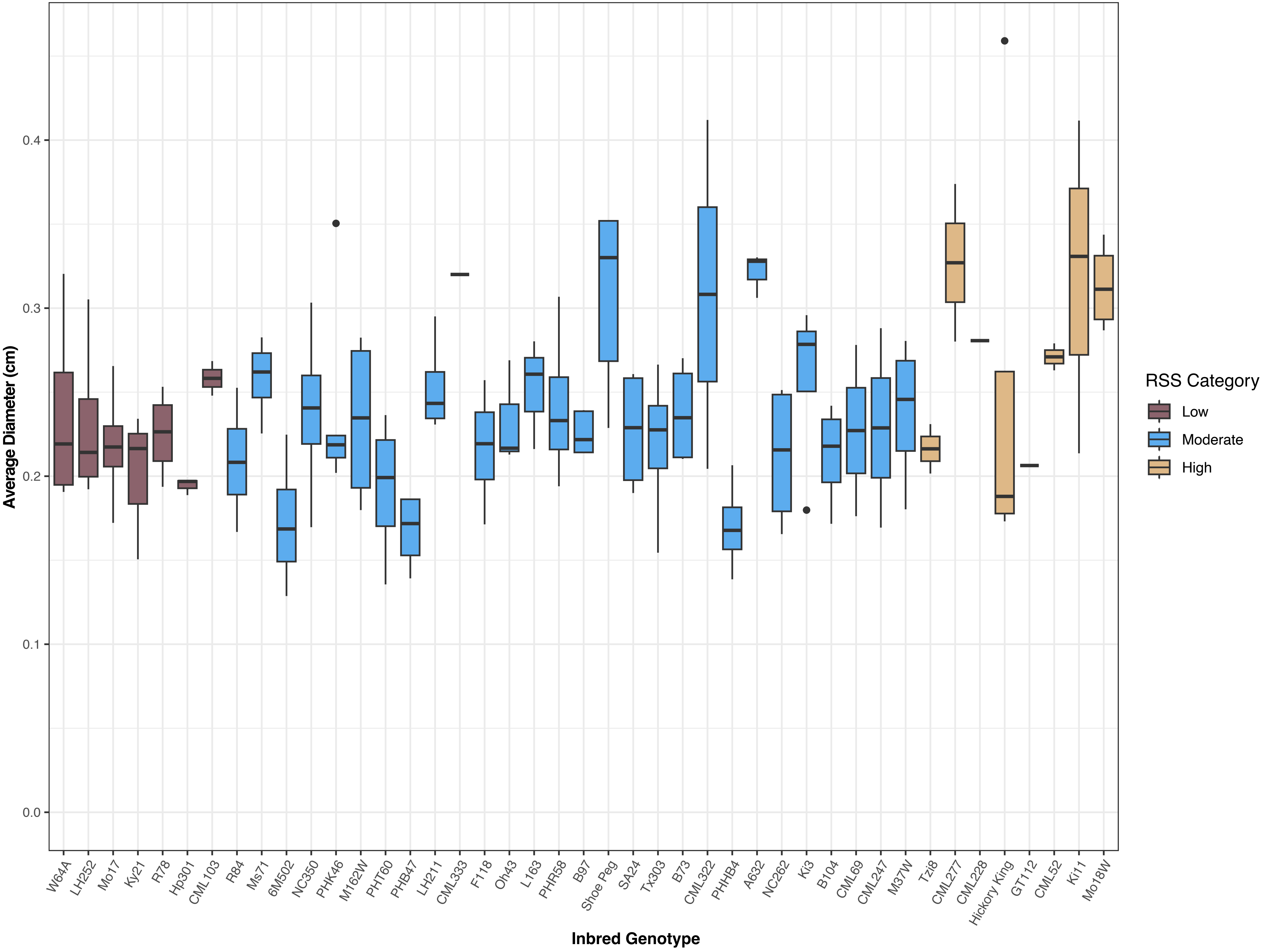

Average Diameter (cm)
0.0
0.1
0.2
0.3
0.4
W64A
LH252
Mo17
Ky21
R78
Hp301
CML103
R84
Ms71
6M502
NC350
PHK46
M162W
PHT60
PHB47
LH211
CML333
F118
Oh43
L163
PHR58
B97
Shoe Peg
SA24
Tx303
B73
CML322
PHHB4
A632
NC262
Ki3
B104
CML69
CML247
M37W
Tzi8
CML277
CML228
Hickory King
GT112
CML52
Ki11
Mo18W
Inbred Genotype
RSS Category
Low
Moderate
High

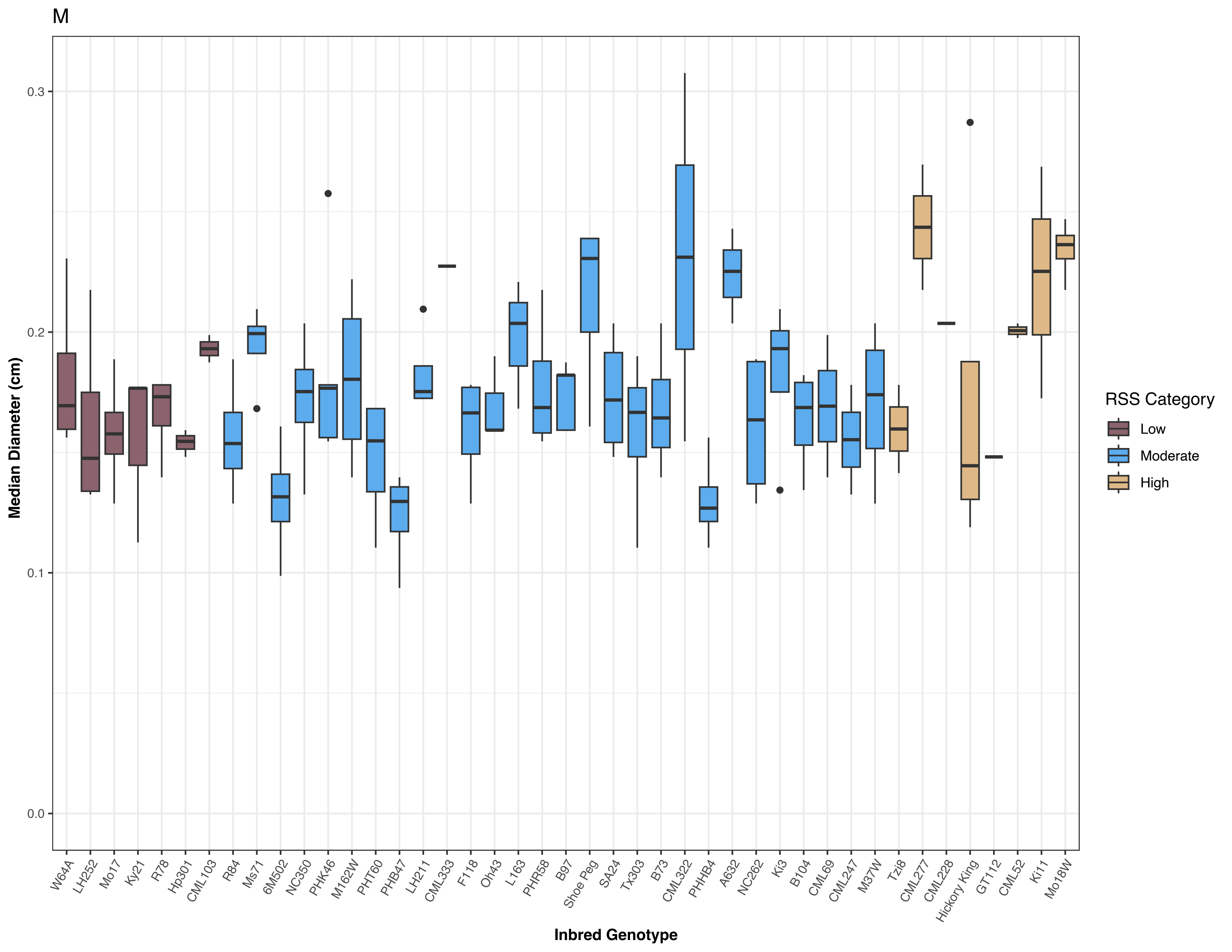

M
Median Diameter (cm)
0.0
0.1
0.2
0.3
Inbred Genotype
W64A
LH252
Mo17
Ky21
R78
Hp301
CML103
R84
Ms71
6M502
NC350
PHK46
M162W
PHT60
PHB47
LH211
CML333
F118
Oh43
L163
PHR58
B97
Shoe Peg
SA24
Tx303
B73
CML322
PHHB4
A632
NC262
Ki3
B104
CML69
CML247
M37W
Tzi8
CML277
CML228
Hickory King
GT112
CML52
Ki11
Mo18W
RSS Category
Low
Moderate
High

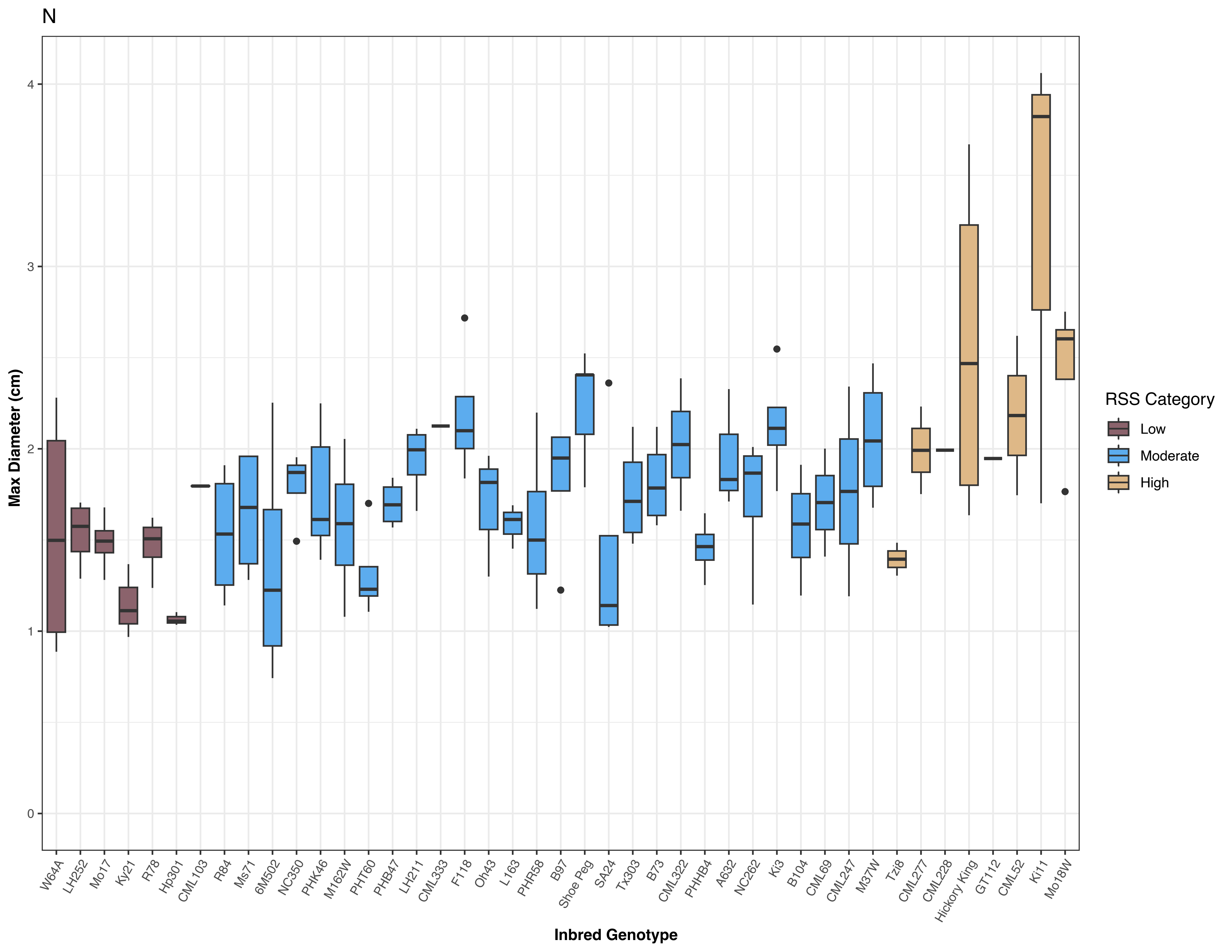
N
Max Diameter (cm)
0
1
2
3
4
W64A
LH252
Mo17
Ky21
R78
Hp301
CML103
R84
Ms71
6M502
NC350
PHK46
M162W
PHT60
PHB47
LH211
CML333
F118
Oh43
L163
PHR58
B97
Shoe Peg
SA24
Tx303
B73
CML322
PHHB4
A632
NC262
Ki3
B104
CML69
CML247
M37W
Tzi8
CML277
CML228
Hickory King
GT112
CML52
Ki11
Mo18W
Inbred Genotype
RSS Category
Low
Moderate
High

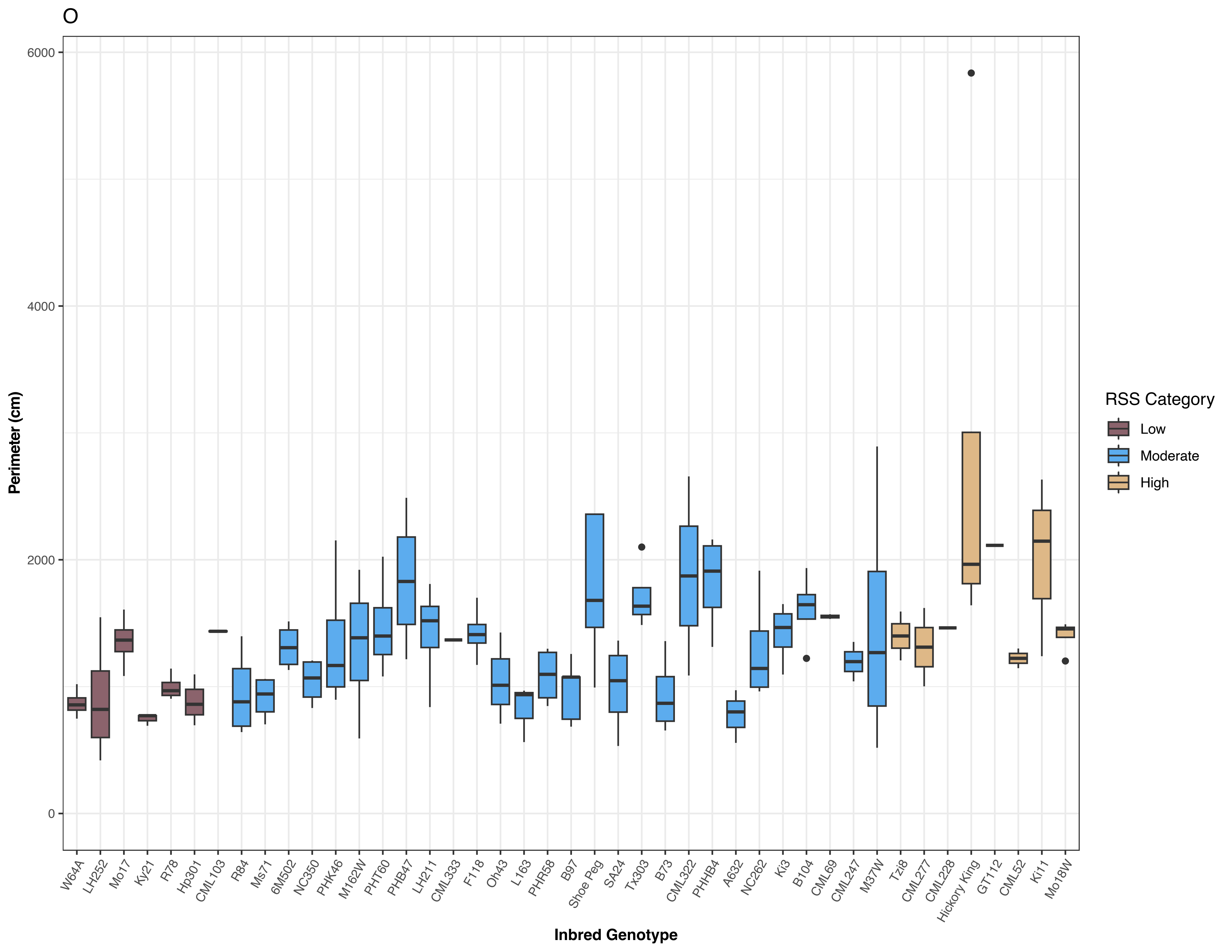
O
6000
4000
2000
0
Perimeter (cm)
W64A
LH252
Mo17
Ky21
R78
Hp301
CML103
R84
Ms71
6M502
NC350
PHK46
M162W
PHT60
PHB47
LH211
CML333
F118
Oh43
L163
PHR58
B97
Shoe Peg
SA24
Tx303
B73
CML322
PHHB4
A632
NC262
Ki3
B104
CML69
CML247
M37W
Tzi8
CML277
CML228
Hickory King
GT112
CML52
Ki11
Mo18W
Inbred Genotype
RSS Category
Low
Moderate
High

P

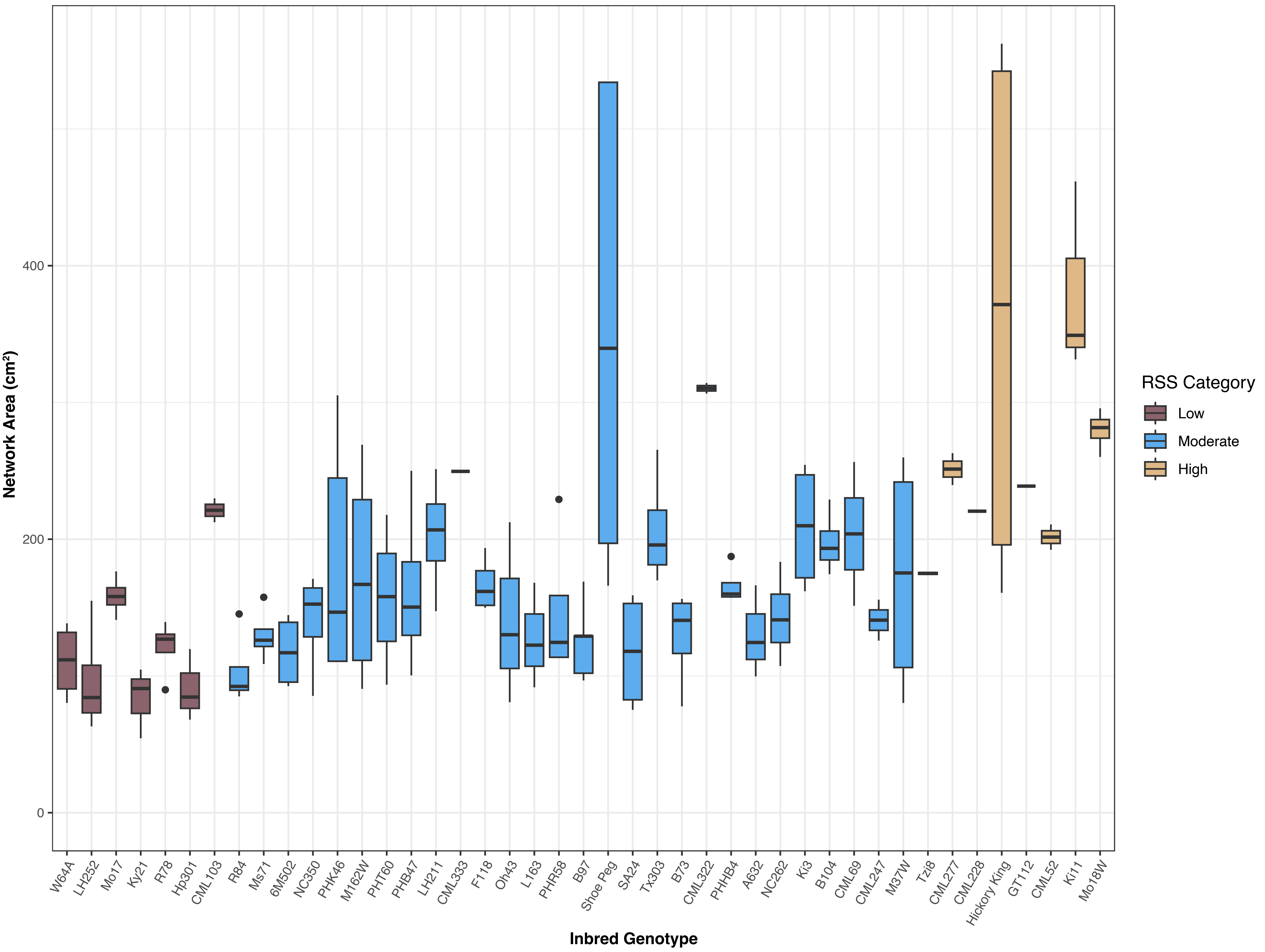

Network Area (cm²)
400
200
0
W64A
LH252
Mo17
Ky21
R78
Hp301
CML103
R84
Ms71
6M502
NC350
PHK46
M162W
PHT60
PHB47
LH211
CML333
F118
Oh43
L163
PHR58
B97
Shoe Peg
SA24
Tx303
B73
CML322
PHHB4
A632
NC262
Ki3
B104
CML69
CML247
M37W
Tzi8
CML277
CML228
Hickory King
GT112
CML52
Ki11
Mo18W
Inbred Genotype
RSS Category
Low
Moderate
High

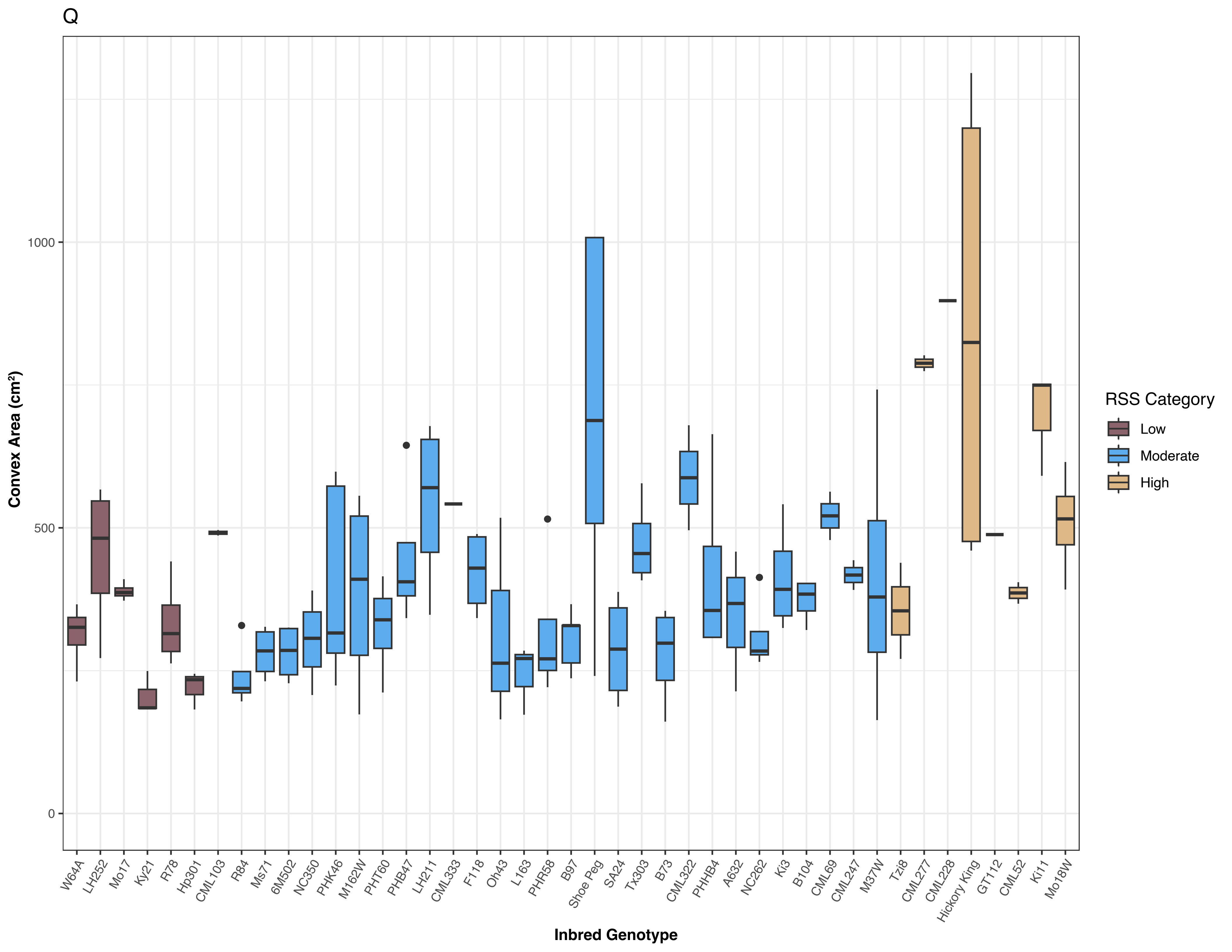
Q
Convex Area (cm²)
1000
500
0
W64A
LH252
Mo17
Ky21
R78
Hp301
CML103
R84
Ms71
6M502
NC350
PHK46
M162W
PHT60
PHB47
LH211
CML333
F118
Oh43
L163
PHR58
B97
Shoe Peg
SA24
Tx303
B73
CML322
PHHB4
A632
NC262
Ki3
B104
CML69
CML247
M37W
Tzi8
CML277
CML228
Hickory King
GT112
CML52
Ki11
Mo18W
Inbred Genotype
RSS Category
Low
Moderate
High

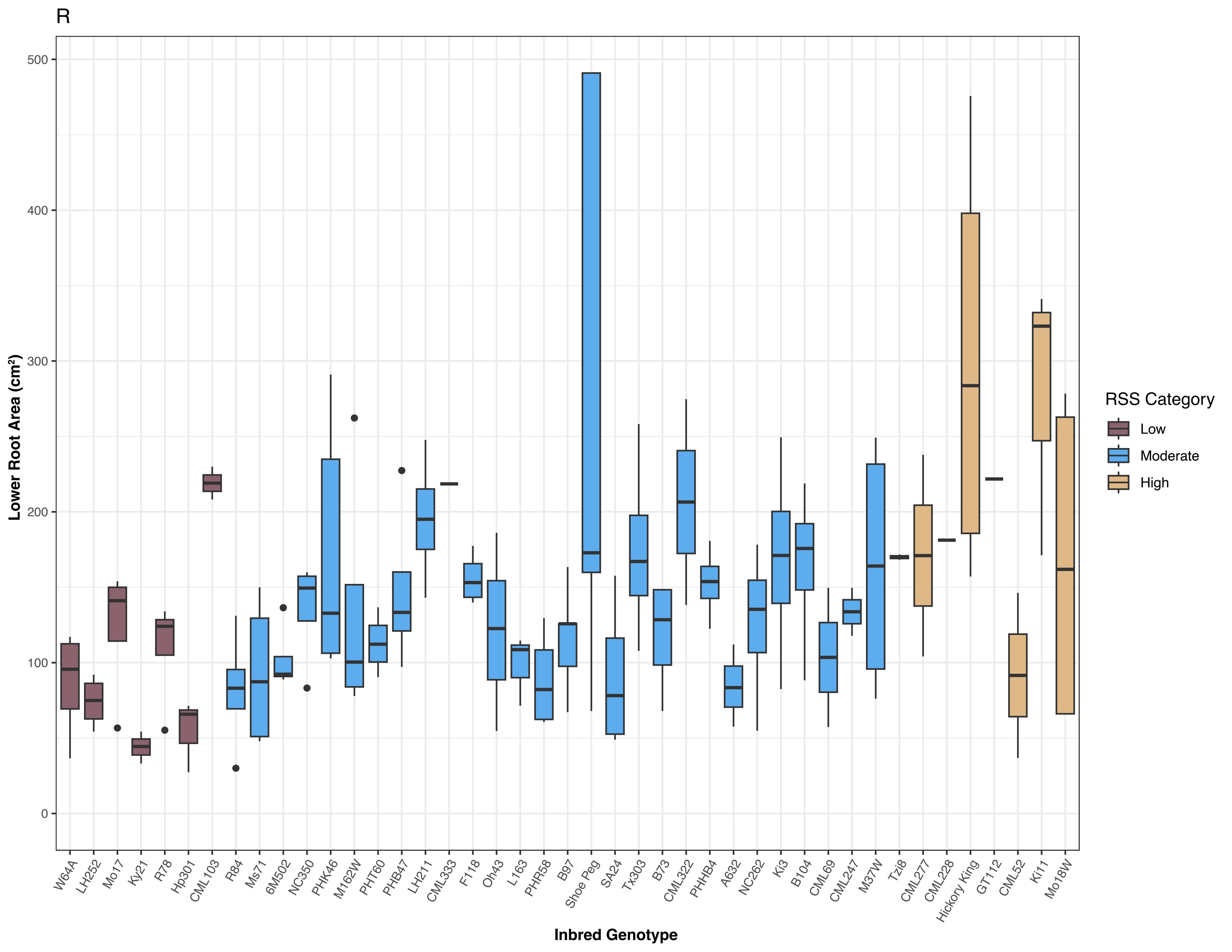
R
Lower Root Area (cm²)
0
100
200
300
400
500
Inbred Genotype
W64A
LH252
Mo17
Ky21
R78
Hp301
CML103
R84
Ms71
6M502
NC350
PHK46
M162W
PHT60
PHB47
LH211
CML333
F118
Oh43
L163
PHR58
B97
Shoe Peg
SA24
Tx303
B73
CML322
PHHB4
A632
NC262
Ki3
B104
CML69
CML247
M37W
Tzi8
CML277
CML228
Hickory King
GT112
CML52
Ki11
Mo18W
RSS Category
Low
Moderate
High

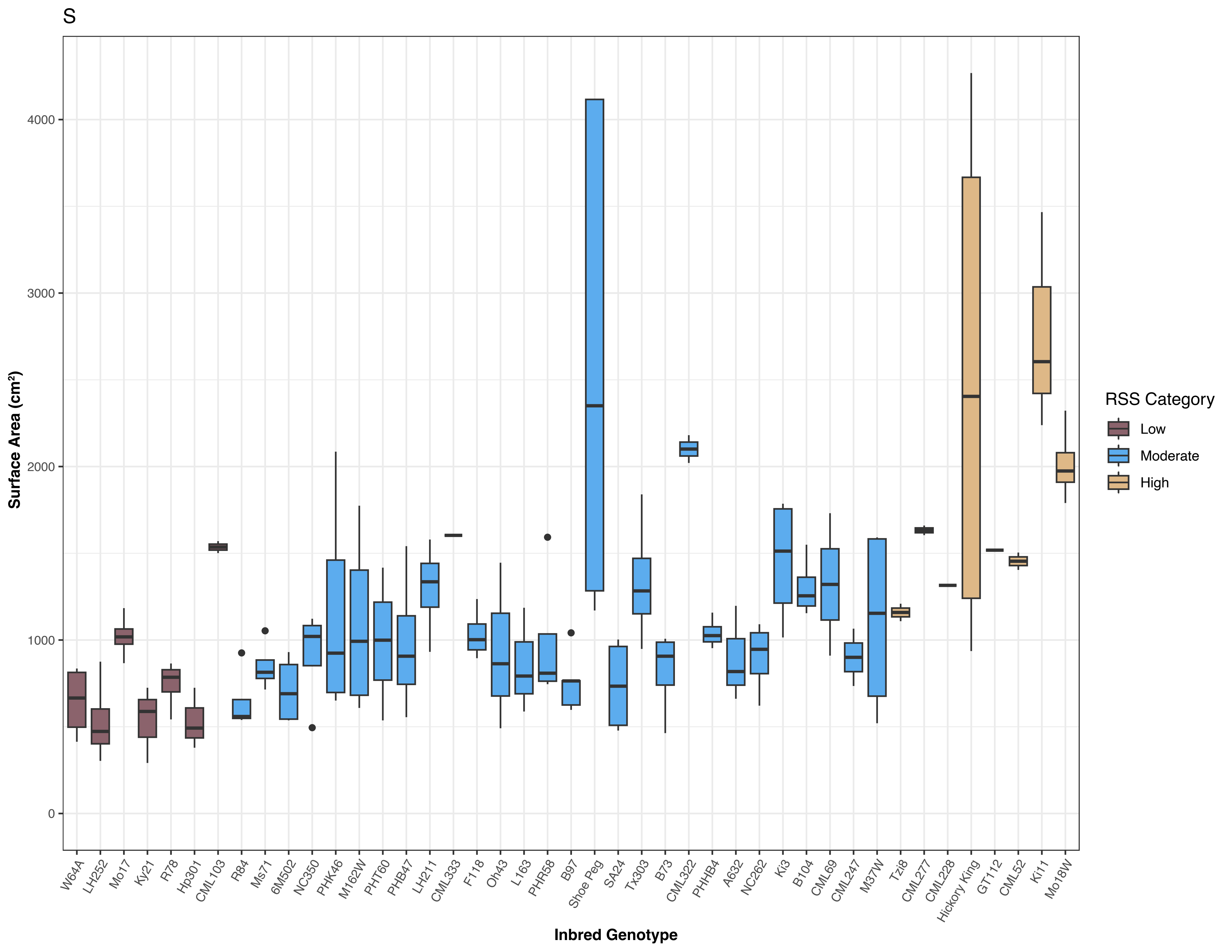

S
Surface Area (cm²)
0
1000
2000
3000
4000
Inbred Genotype
W64A
LH252
Mo17
Ky21
R78
Hp301
CML103
R84
Ms71
6M502
NC350
PHK46
M162W
PHT60
PHB47
LH211
CML333
F118
Oh43
L163
PHR58
B97
Shoe Peg
SA24
Tx303
B73
CML322
PHHB4
A632
NC262
Ki3
B104
CML69
CML247
M37W
Tzi8
CML277
CML228
Hickory King
GT112
CML52
Ki11
Mo18W
RSS Category
Low
Moderate
High

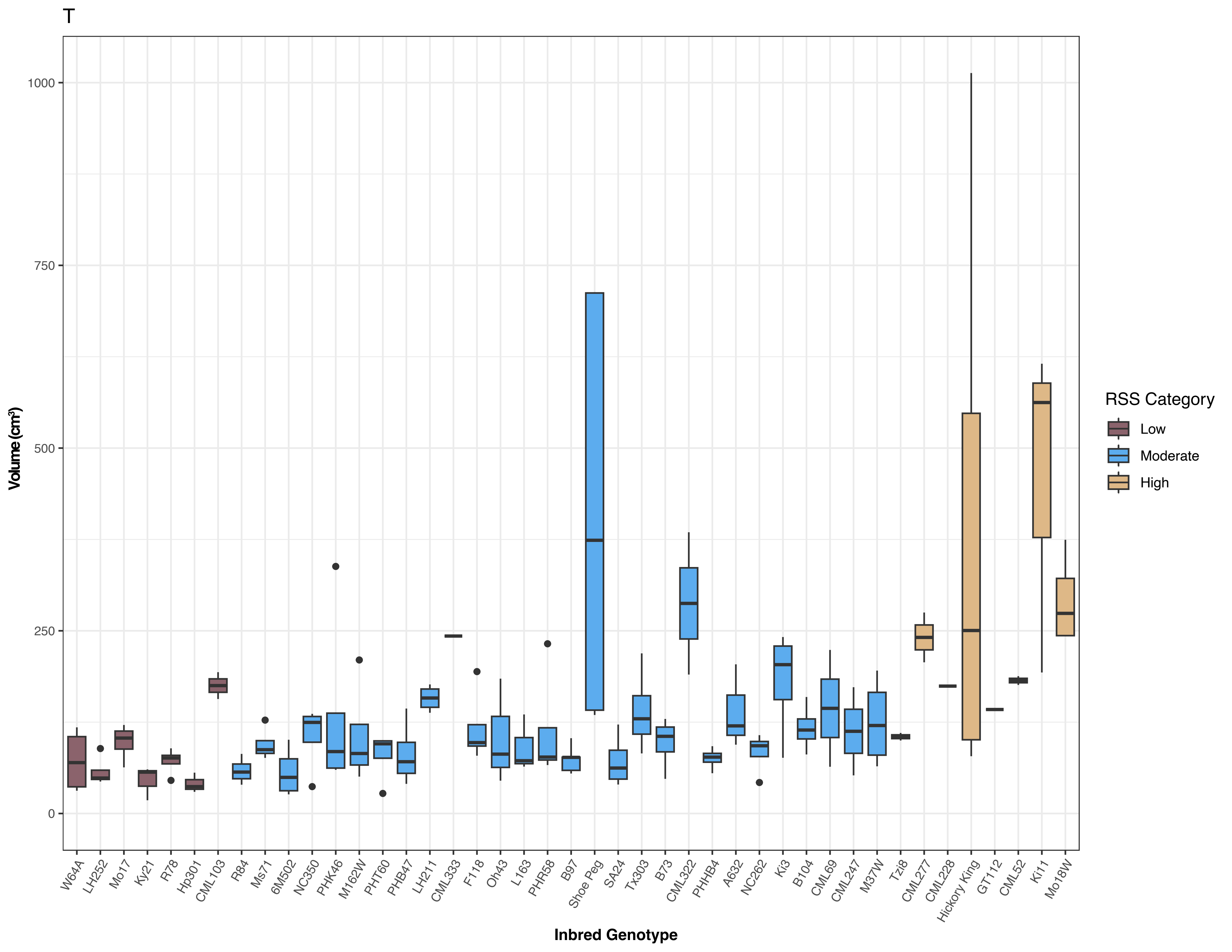
T
Volume (cm³)
1000
750
500
250
0
Inbred Genotype
W64A
LH252
Mo17
Ky21
R78
Hp301
CML103
R84
Ms71
6M502
NC350
PHK46
M162W
PHT60
PHB47
LH211
CML333
F118
Oh43
L163
PHR58
B97
Shoe Peg
SA24
Tx303
B73
CML322
PHHB4
A632
NC262
Ki3
B104
CML69
CML247
M37W
Tzi8
CML277
CML228
Hickory King
GT112
CML52
Ki11
Mo18W
RSS Category
Low
Moderate
High

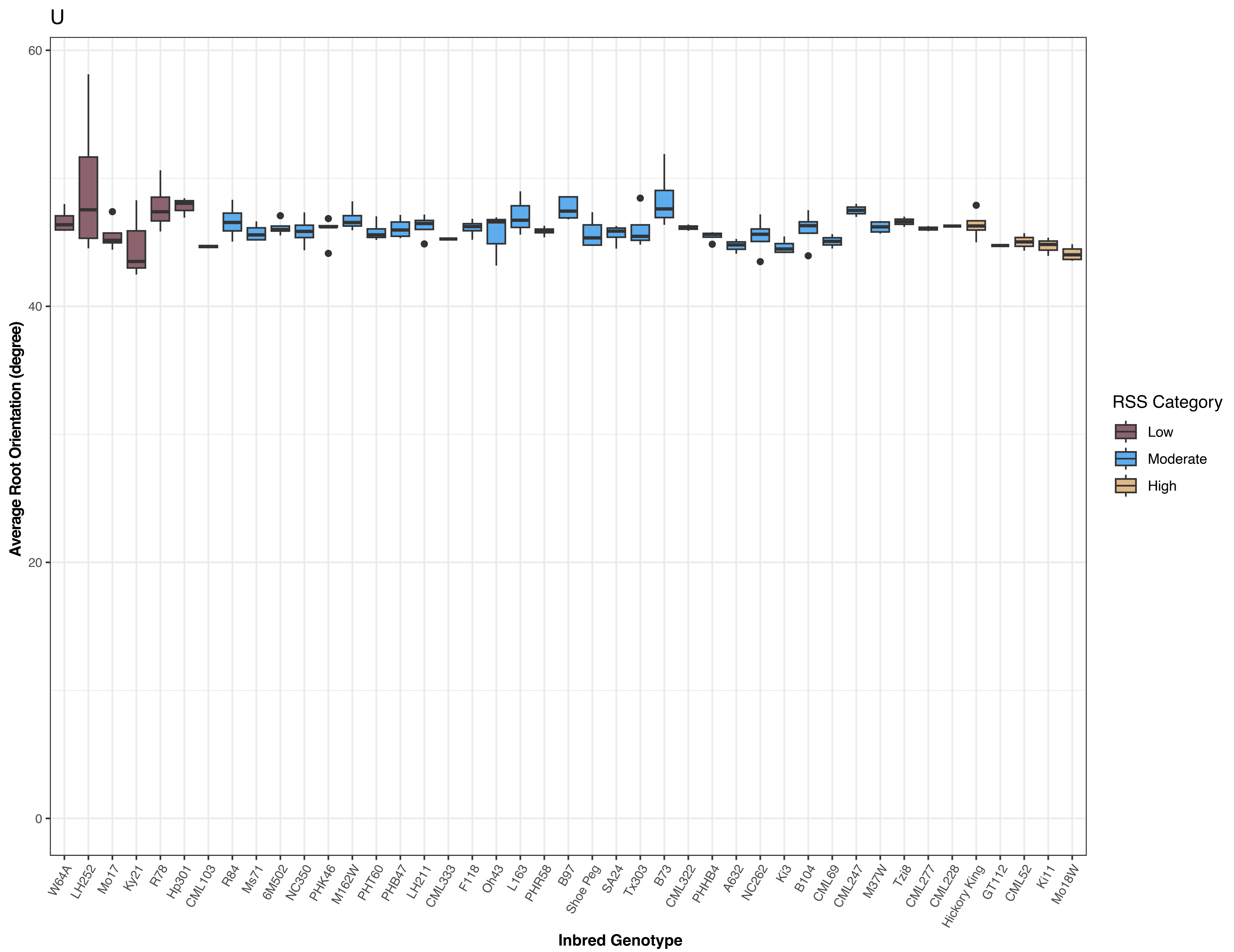


Figure S2. Inbred genotypes varied for all below-ground root architectural traits. (A) Median number of roots, (B) maximum number of roots, (C) number of root tips, (D) width/depth, (E) solidity, (F) shallow angle frequency, (G) medium angle frequency, (H) steep angle frequency, (I) total root length, (J) depth, (K) maximum width, (L) average diameter, (M) median diameter, (N) maximum diameter, (O) perimeter, (P) network area, (Q) convex area, (R) lower root area, (S) surface area, (T) volume, and (U) average root orientation. Colors indicate root system stiffness categories.

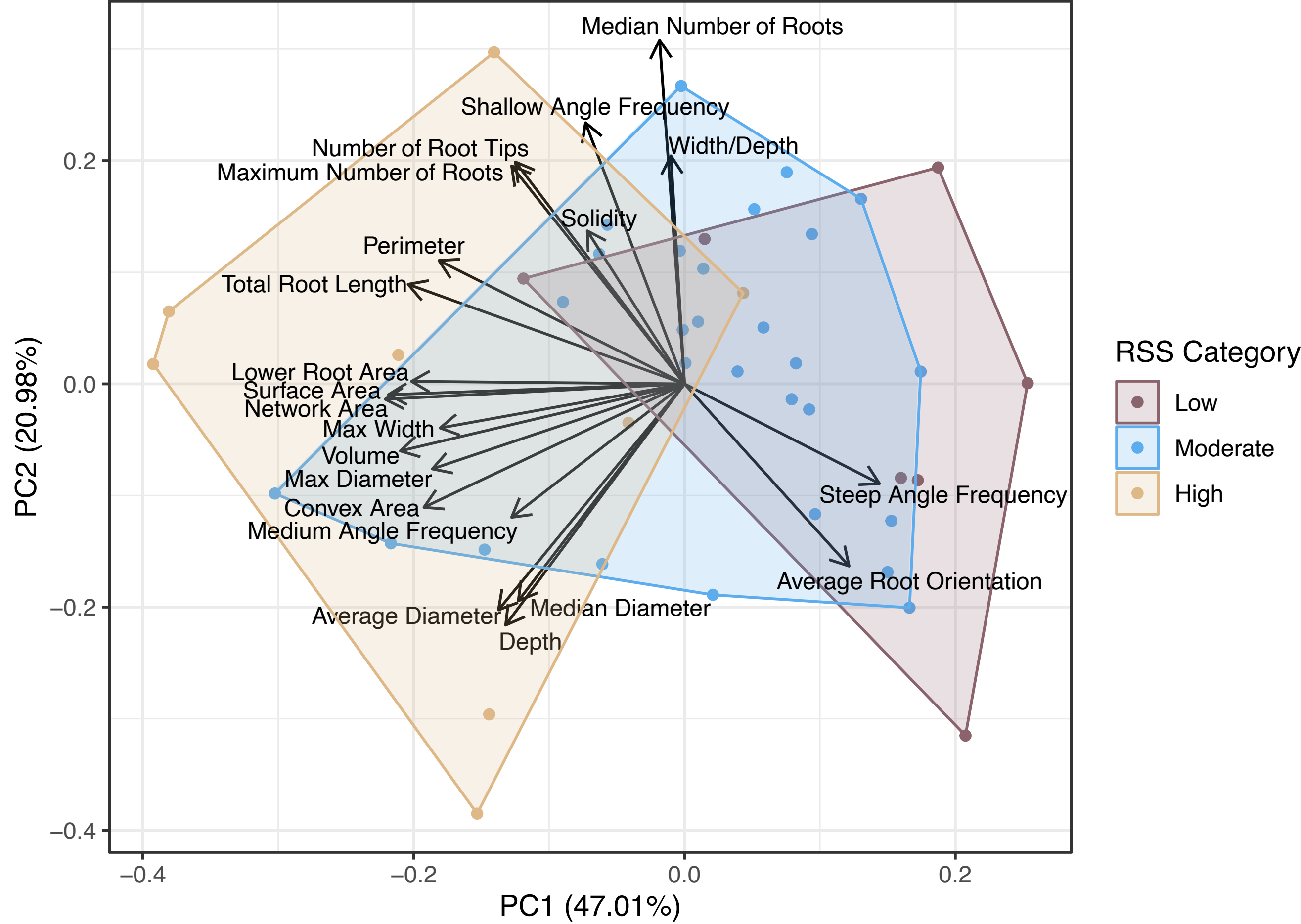


Figure S3. A PCA with trait loadings showed that inbred genotypes categorized as having a high root system stiffness had larger and more extensive below-ground root systems.

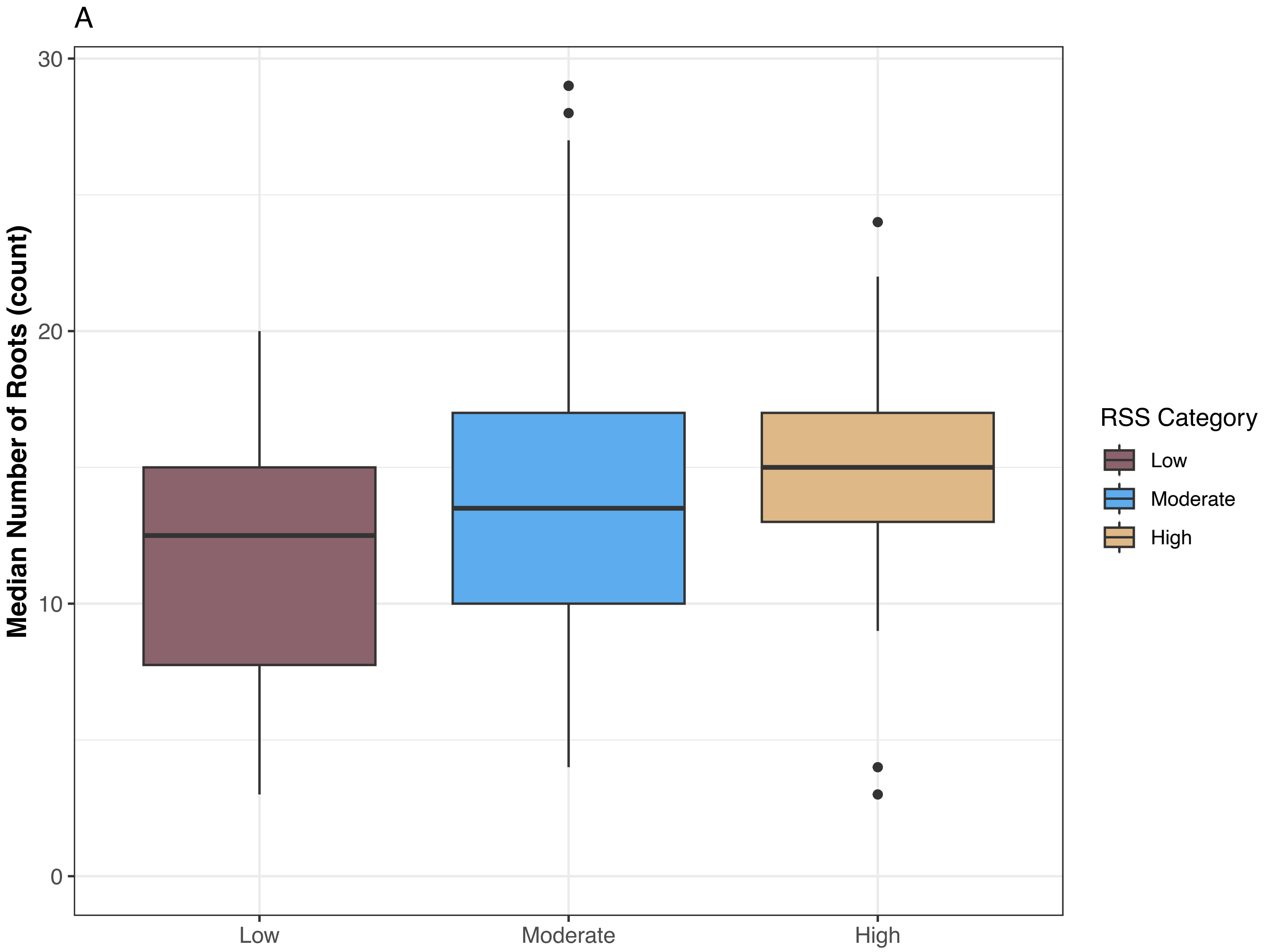
A
Median Number of Roots (count)
30
20
10
0
Low
Moderate
High
RSS Category
Low
Moderate
High

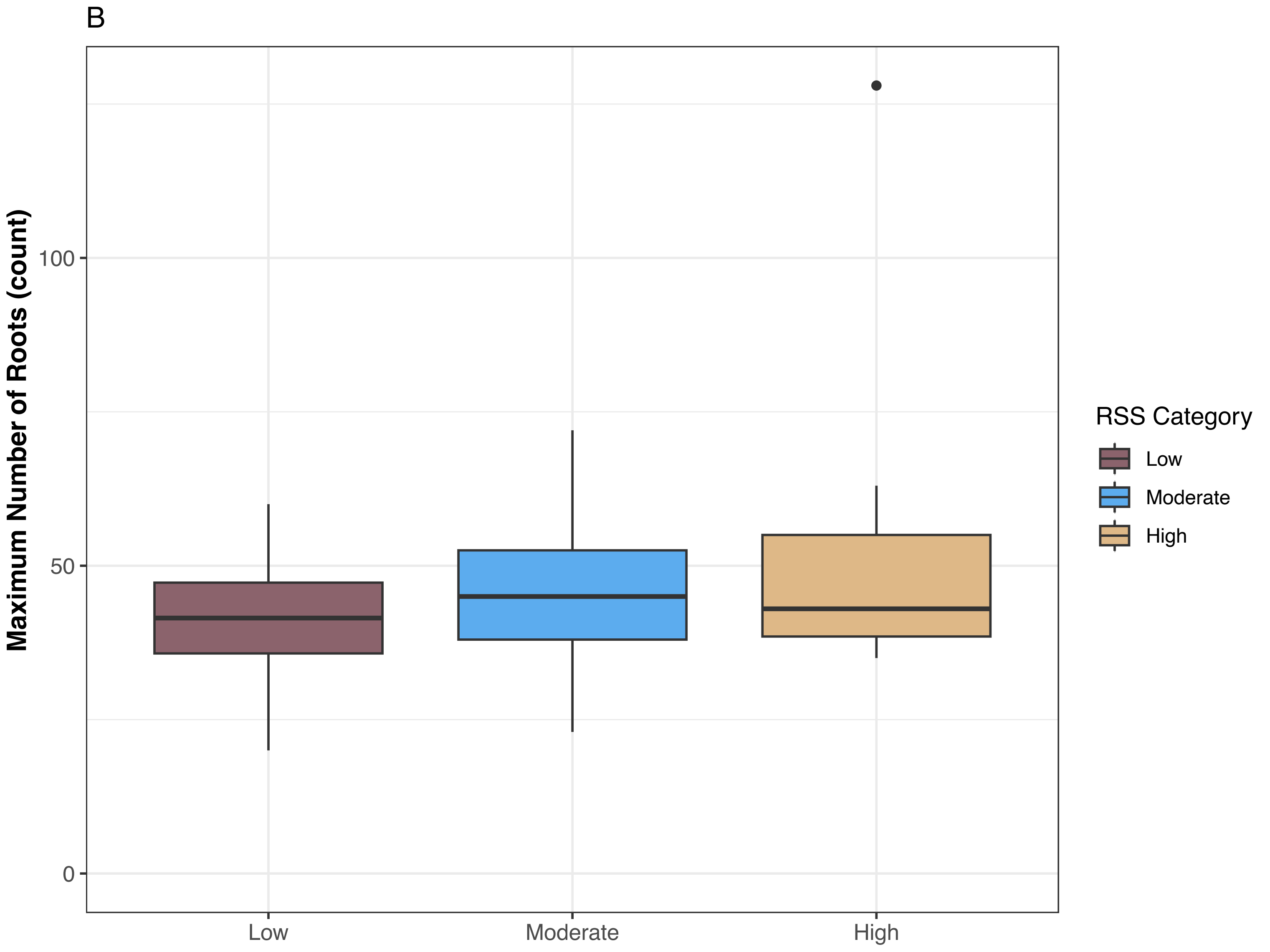
B
Maximum Number of Roots (count)
100
50
0
Low
Moderate
High
RSS Category
Low
Moderate
High

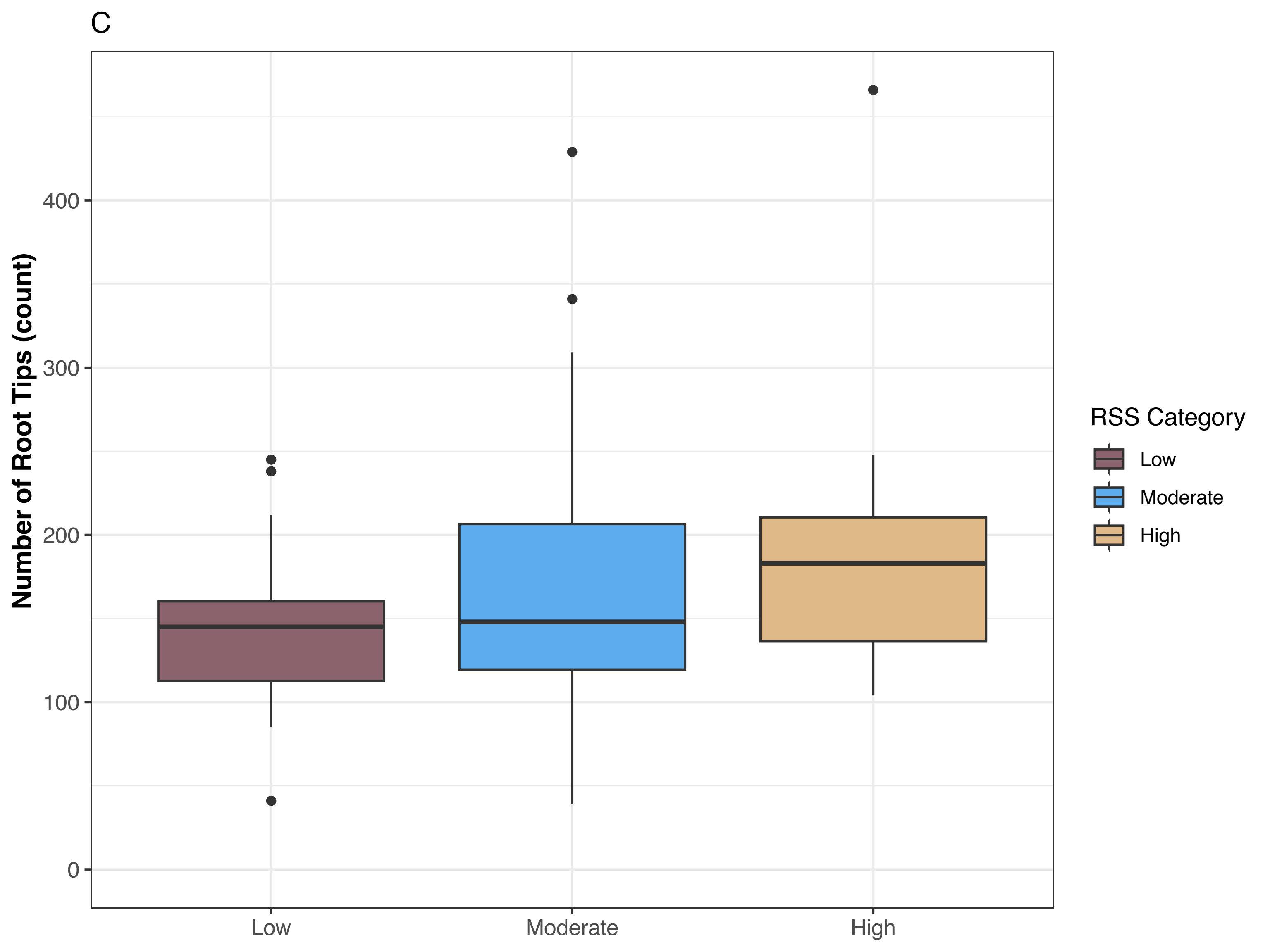
C
Number of Root Tips (count)
0
100
200
300
400
Low
Moderate
High
RSS Category
Low
Moderate
High

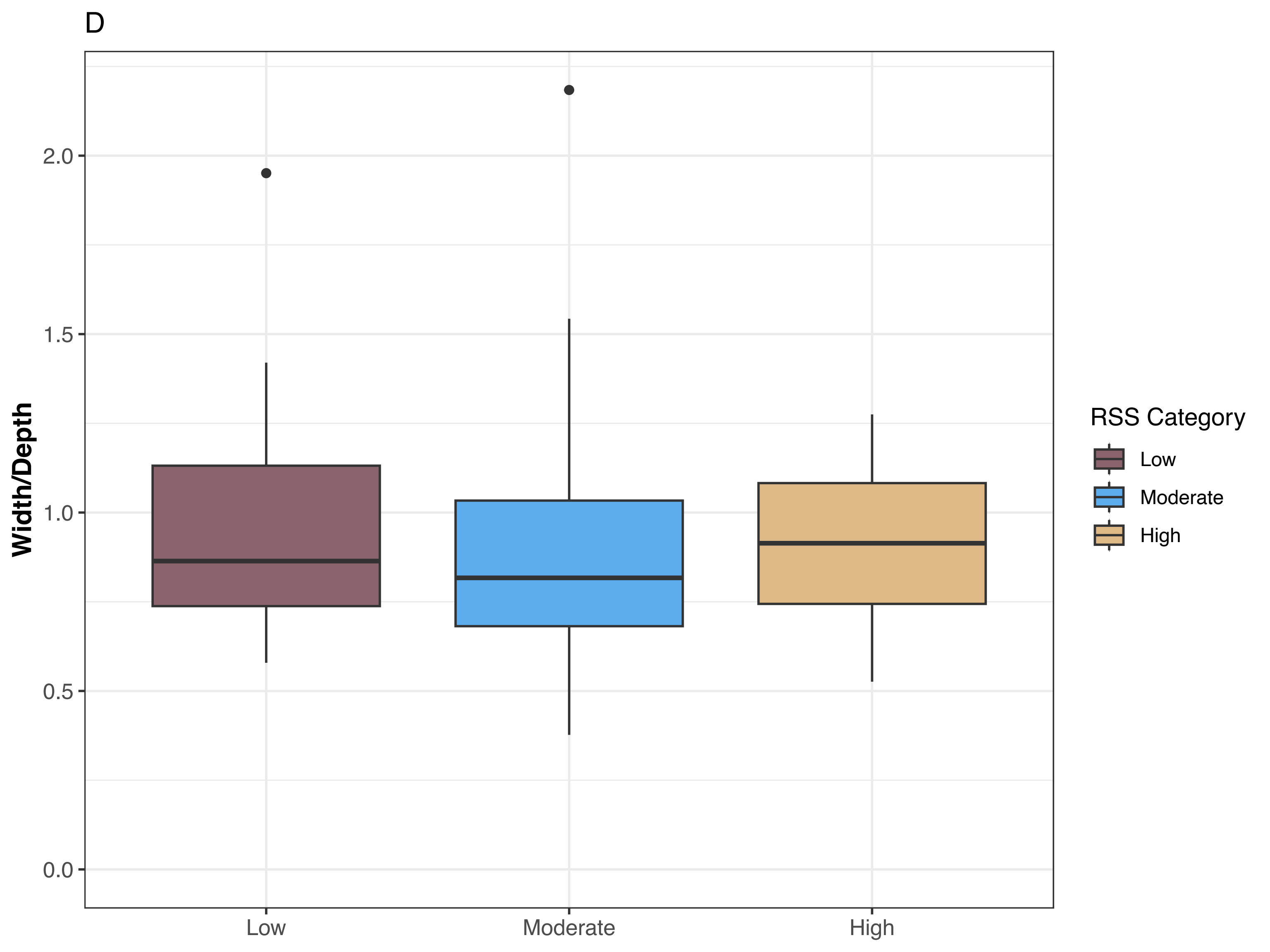
D
Width/Depth
2.0
1.5
1.0
0.5
0.0
Low
Moderate
High
RSS Category
Low
Moderate
High

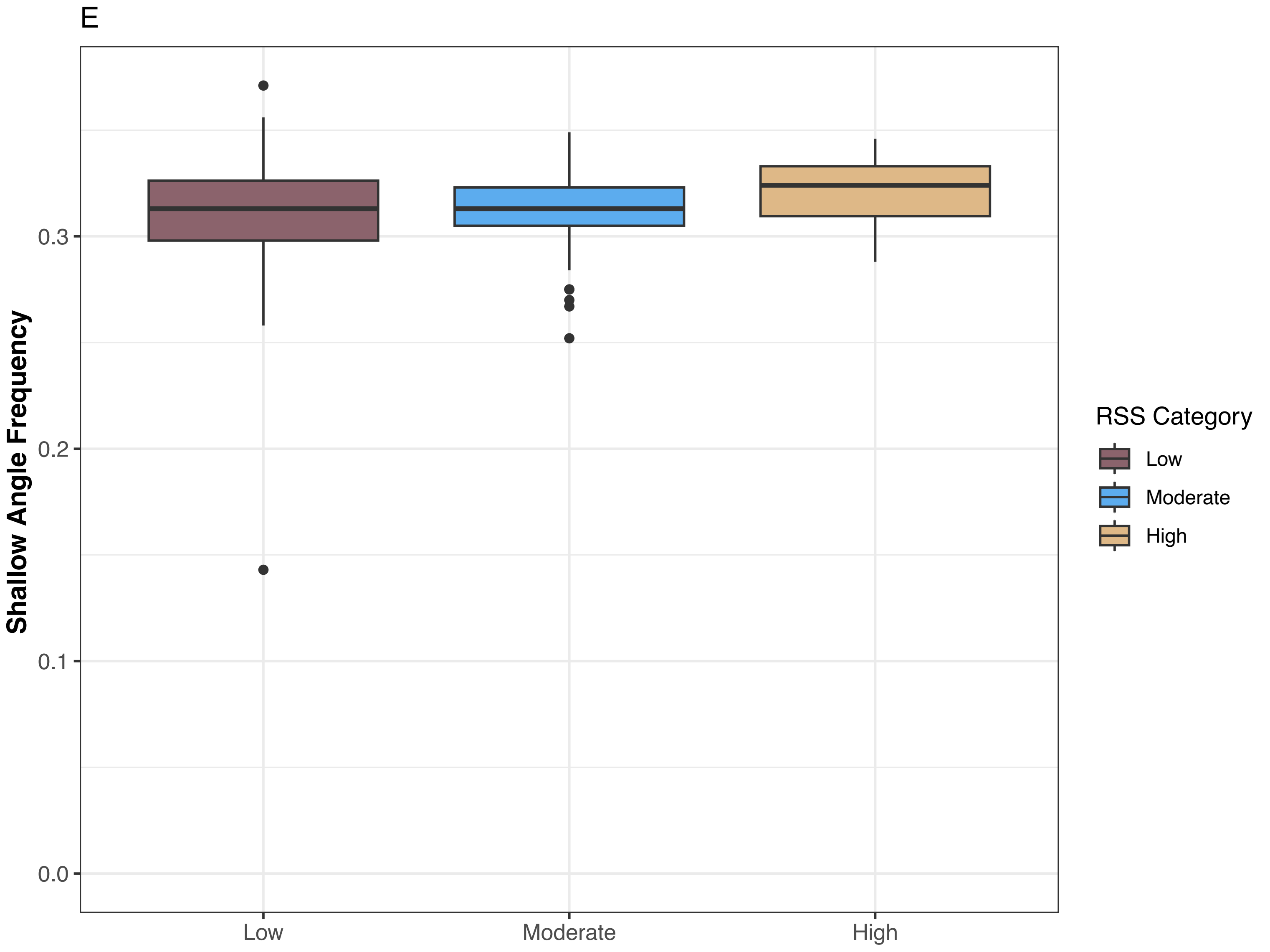
E
Shallow Angle Frequency
0.0
0.1
0.2
0.3
Low
Moderate
High
RSS Category
Low
Moderate
High

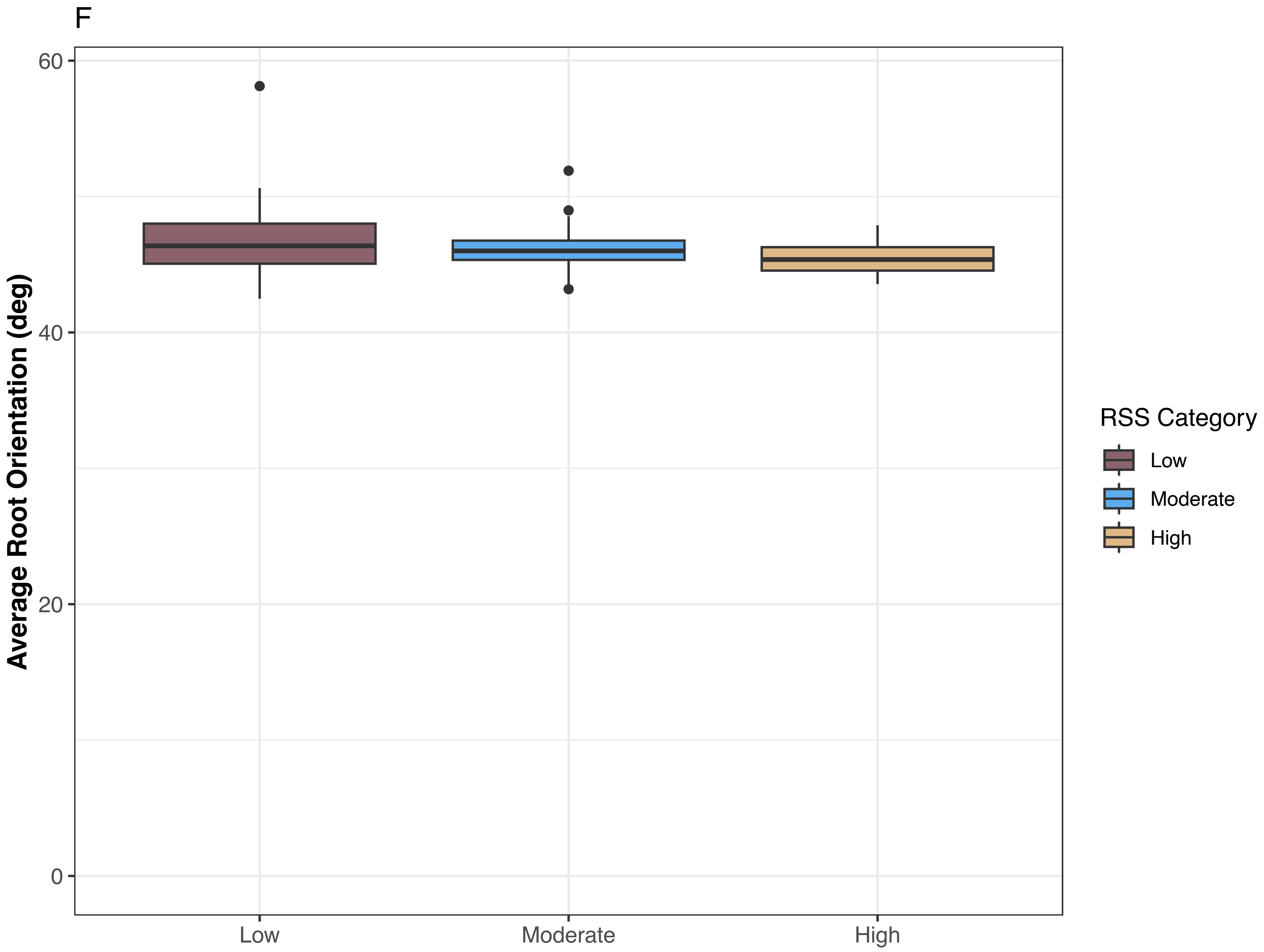


Figure S4. Root system stiffness categories were not significant for some below-ground root architectural traits. (A) Median number of roots (B) maximum number of roots, (C) number of root tips, (D) width/depth, (E) shallow angle frequency, and (F) average root orientation. While traits varied among root system stiffness categories, the difference did not pass the threshold for significance. Colors indicate root system stiffness categories.

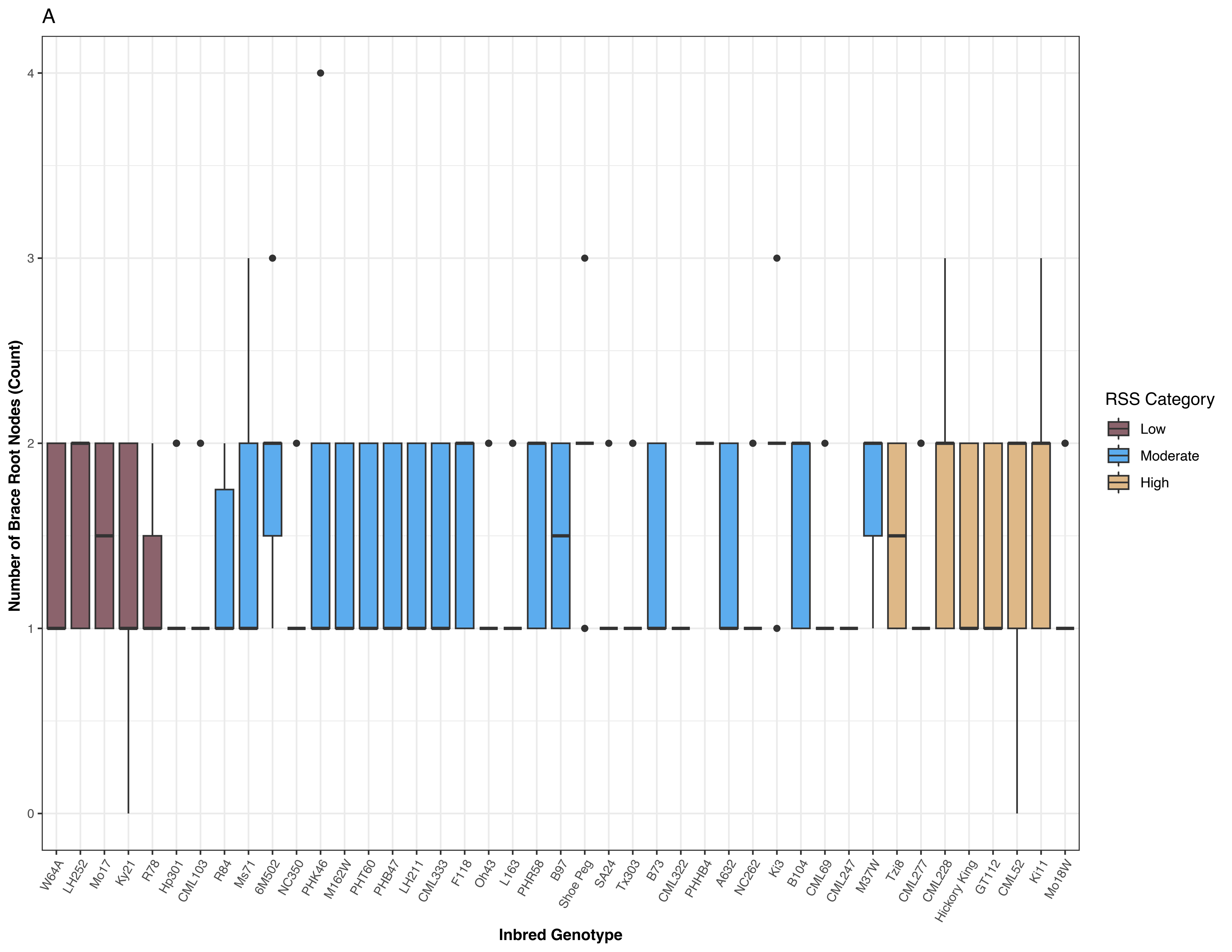
A
Number of Brace Root Nodes (Count)
4
3
2
1
0
W64A
LH252
Mo17
Ky21
R78
Hp301
CML103
R84
Ms71
6M502
NC350
PHK46
M162W
PHT60
PHB47
LH211
CML333
F118
Oh43
L163
PHR58
B97
Shoe Peg
SA24
Tx303
B73
CML322
PHHB4
A632
NC262
Ki3
B104
CML69
CML247
M37W
Tzi8
CML277
CML228
Hickory King
GT112
CML52
Ki11
Mo18W
Inbred Genotype
RSS Category
Low
Moderate
High

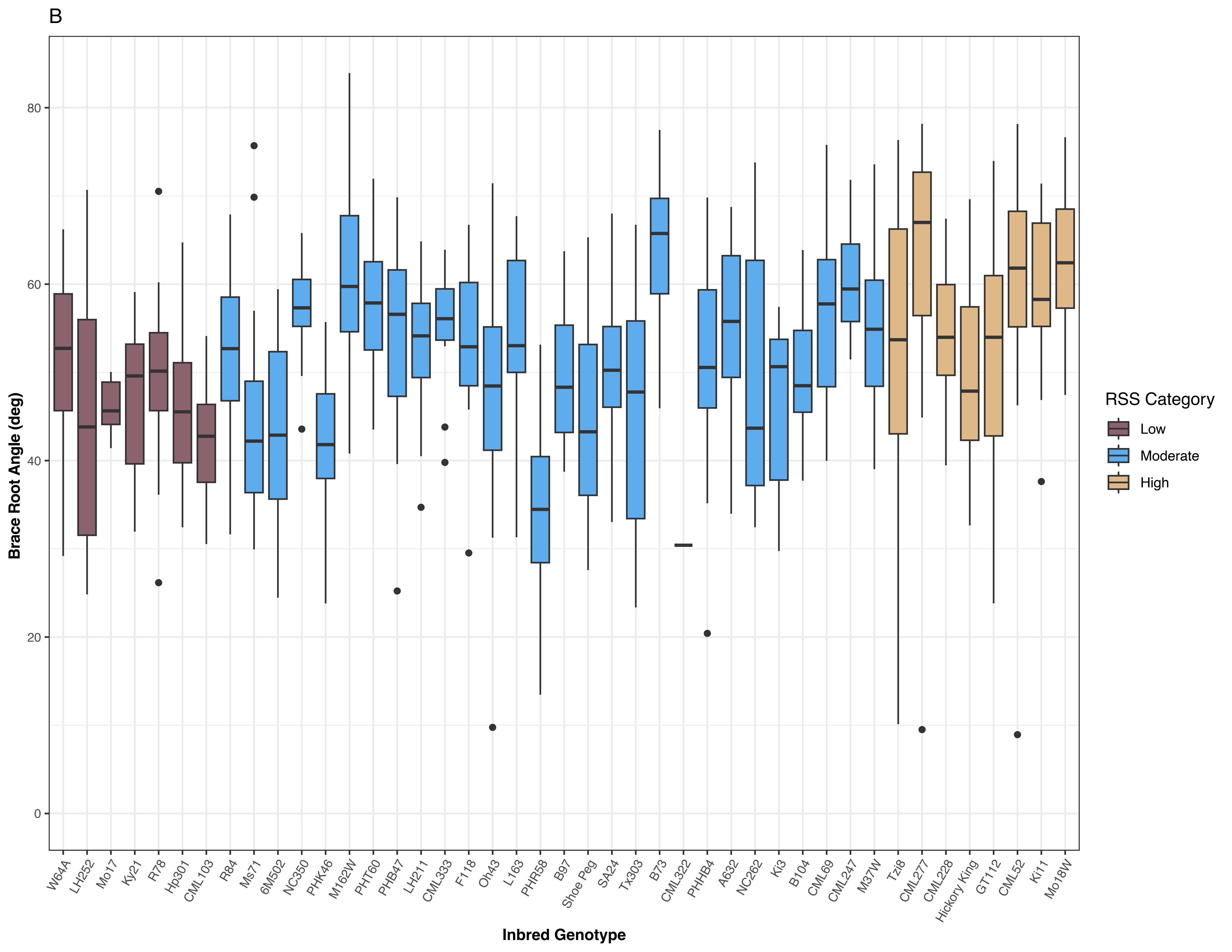

B
Brace Root Angle (deg)
0
20
40
60
80
W64A
LH252
Mo17
Ky21
R78
Hp301
CML103
R84
Ms71
6M502
NC350
PHK46
M162W
PHT60
PHB47
LH211
CML333
F118
Oh43
L163
PHR58
B97
Shoe Peg
SA24
Tx303
B73
CML322
PHHB4
A632
NC262
Ki3
B104
CML69
CML247
M37W
Tzi8
CML277
CML228
Hickory King
GT112
CML52
Ki11
Mo18W
Inbred Genotype
RSS Category
Low
Moderate
High

C

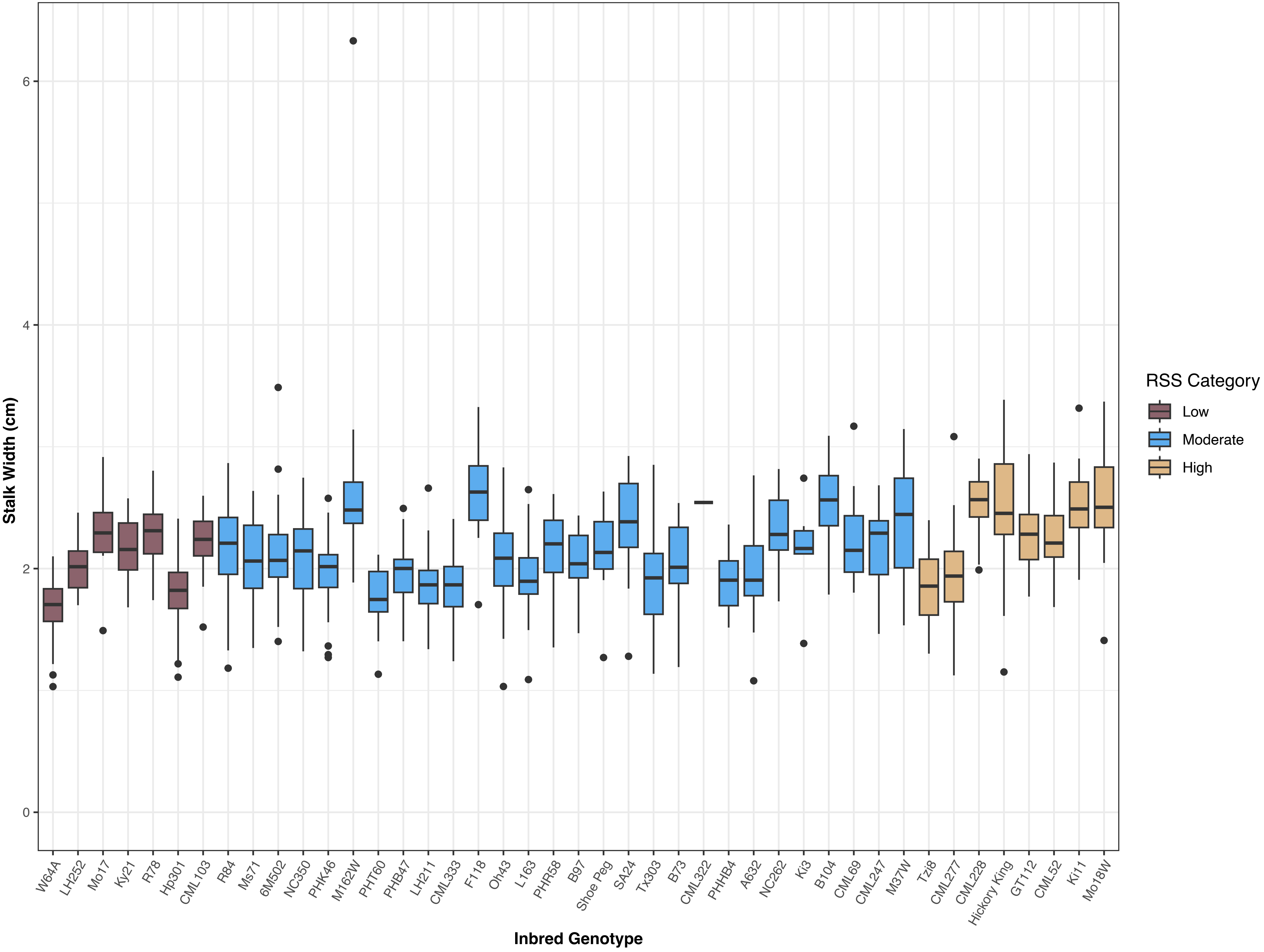

Stalk Width (cm)
0
2
4
6
W64A
LH252
Mo17
Ky21
R78
Hp301
CML103
R84
Ms71
6M502
NC350
PHK46
M162W
PHT60
PHB47
LH211
CML333
F118
Oh43
L163
PHR58
B97
Shoe Peg
SA24
Tx303
B73
CML322
PHHB4
A632
NC262
Ki3
B104
CML69
CML247
M37W
Tzi8
CML277
CML228
Hickory King
GT112
CML52
Ki11
Mo18W
Inbred Genotype
RSS Category
Low
Moderate
High

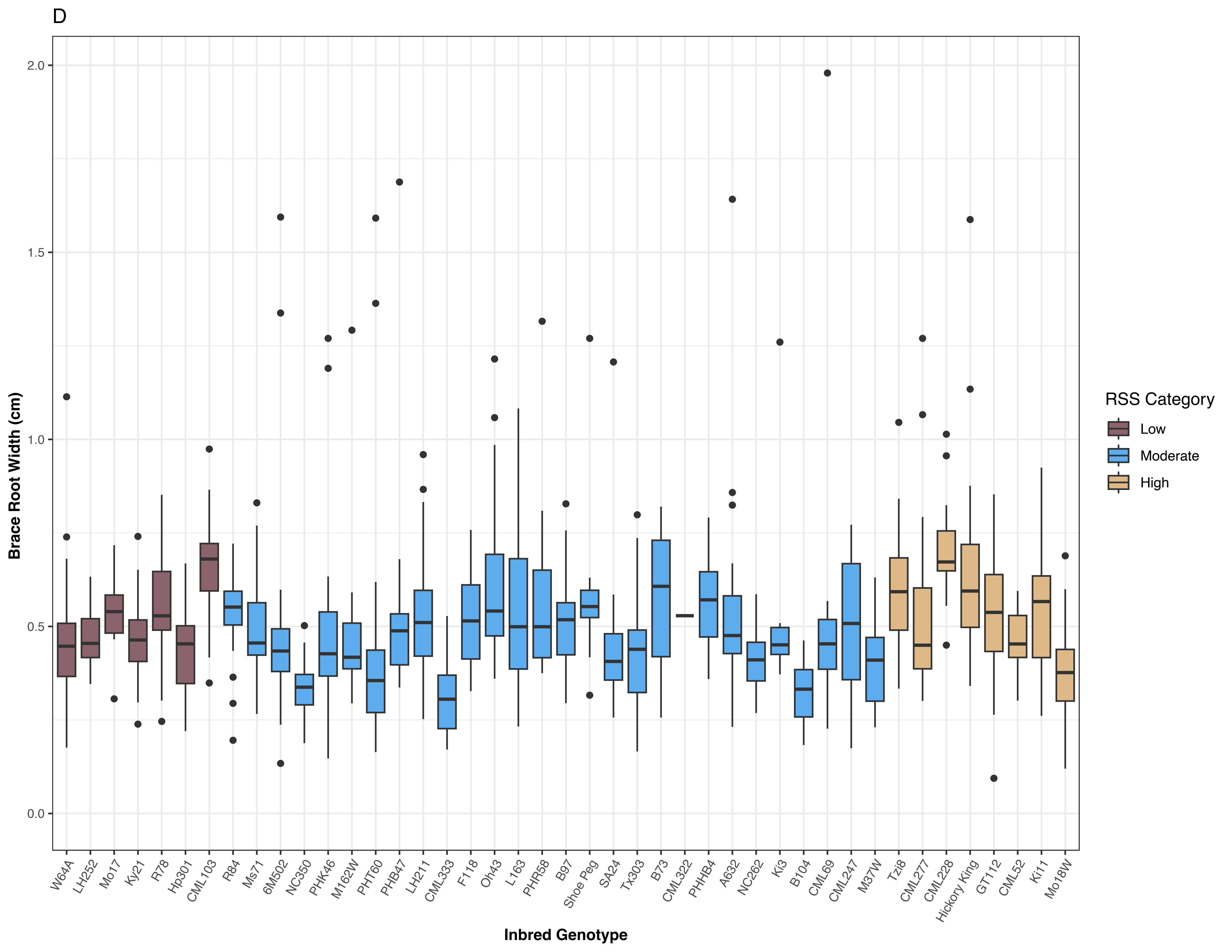
D
Brace Root Width (cm)
2.0
1.5
1.0
0.5
0.0
W64A
LH252
Mo17
Ky21
R78
Hp301
CML103
R84
Ms71
6M502
NC350
PHK46
M162W
PHT60
PHB47
LH211
CML333
F118
Oh43
L163
PHR58
B97
Shoe Peg
SA24
Tx303
B73
CML322
PHHB4
A632
NC262
Ki3
B104
CML69
CML247
M37W
Tzi8
CML277
CML228
Hickory King
GT112
CML52
Ki11
Mo18W
Inbred Genotype
RSS Category
Low
Moderate
High

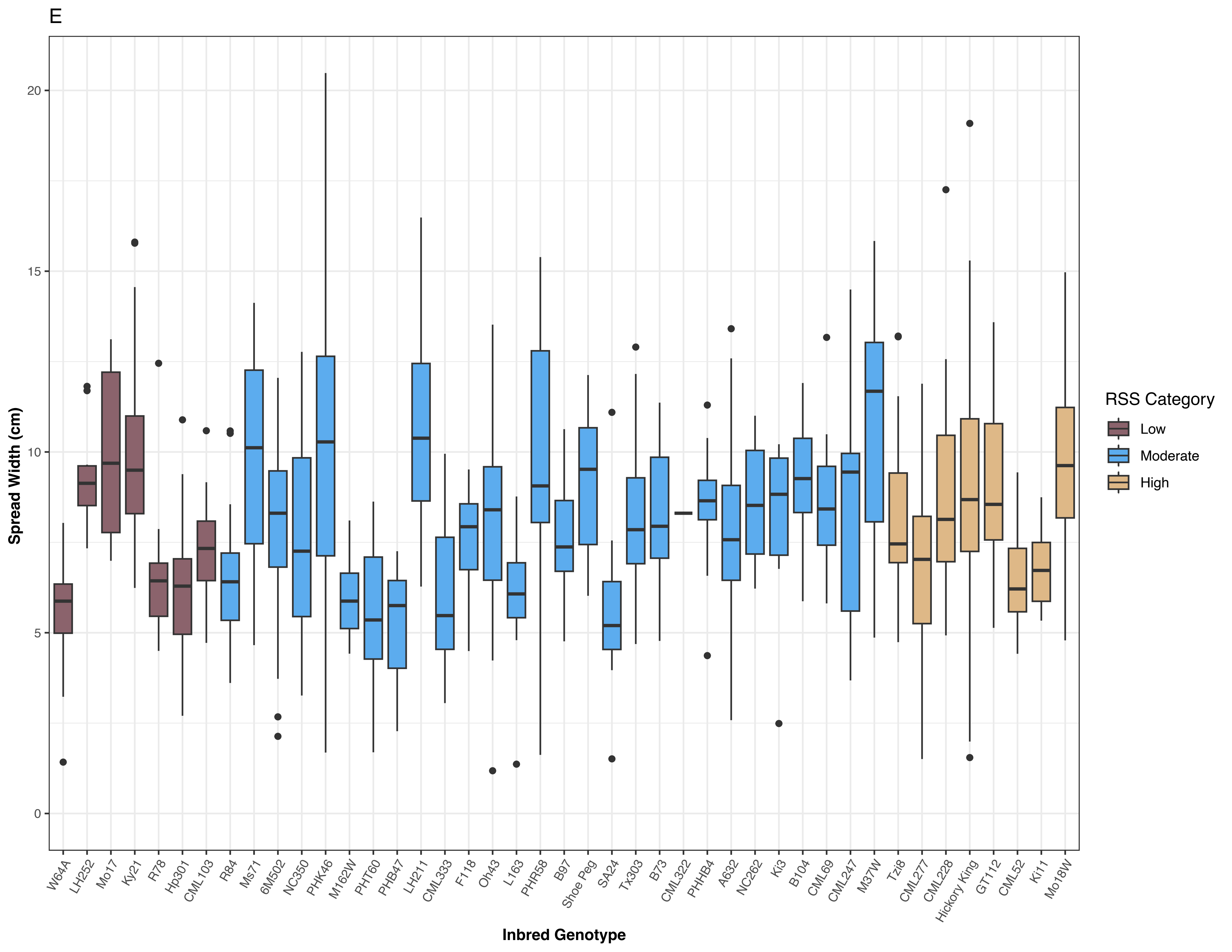

E
Spread Width (cm)
0
5
10
15
20
W64A
LH252
Mo17
Ky21
R78
Hp301
CML103
R84
Ms71
6M502
NC350
PHK46
M162W
PHT60
PHB47
LH211
CML333
F118
Oh43
L163
PHR58
B97
Shoe Peg
SA24
Tx303
B73
CML322
PHHB4
A632
NC262
Ki3
B104
CML69
CML247
M37W
Tzi8
CML277
CML228
Hickory King
GT112
CML52
Ki11
Mo18W
Inbred Genotype
RSS Category
Low
Moderate
High

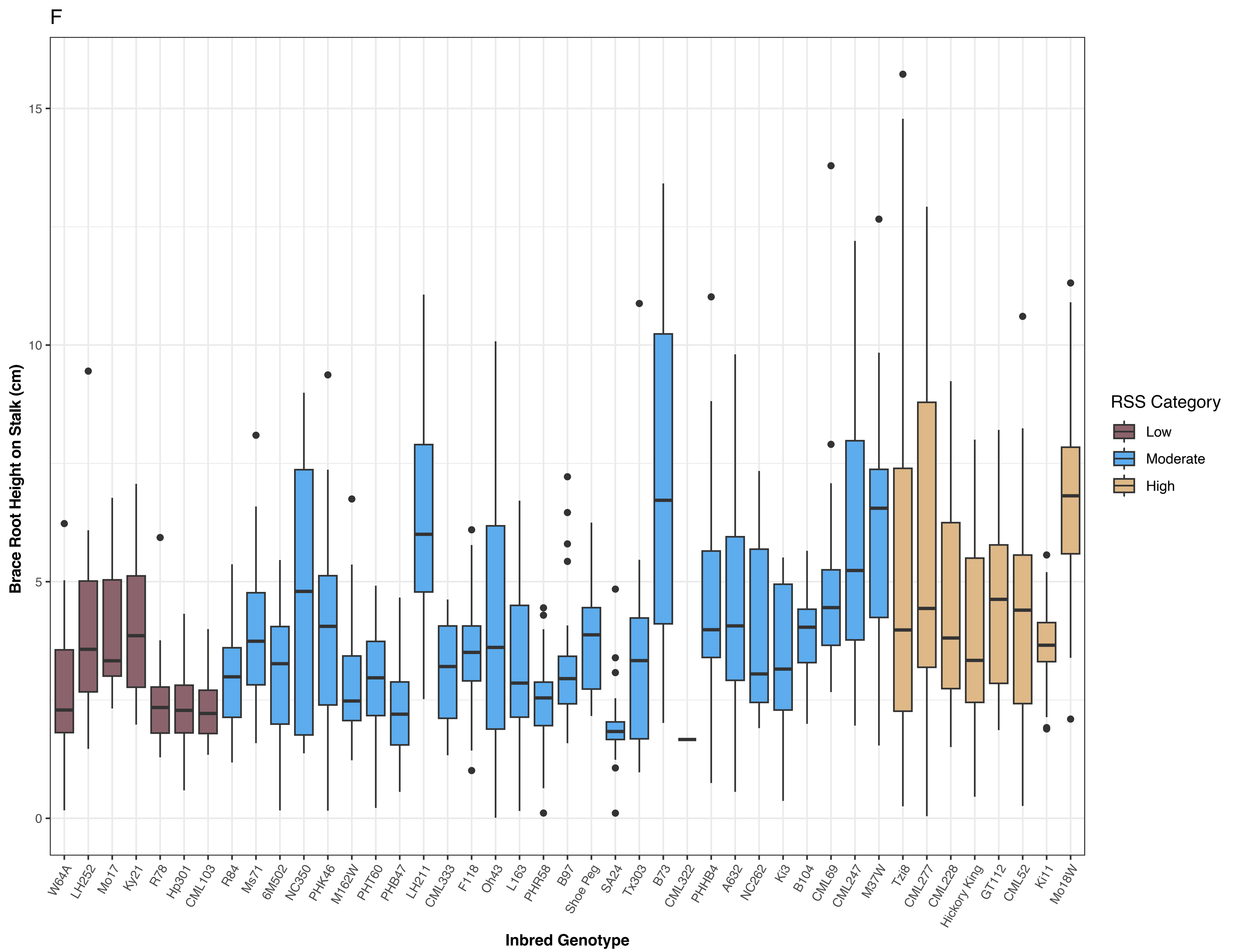


Figure S5. Inbred genotypes varied for all above-ground root architectural traits. (A) Number of brace root nodes, (B) brace root angle, (C) stalk width, (D) brace root width, (E) brace root spread width, and (F) brace root height on stalk. Colors indicate root system stiffness categories.

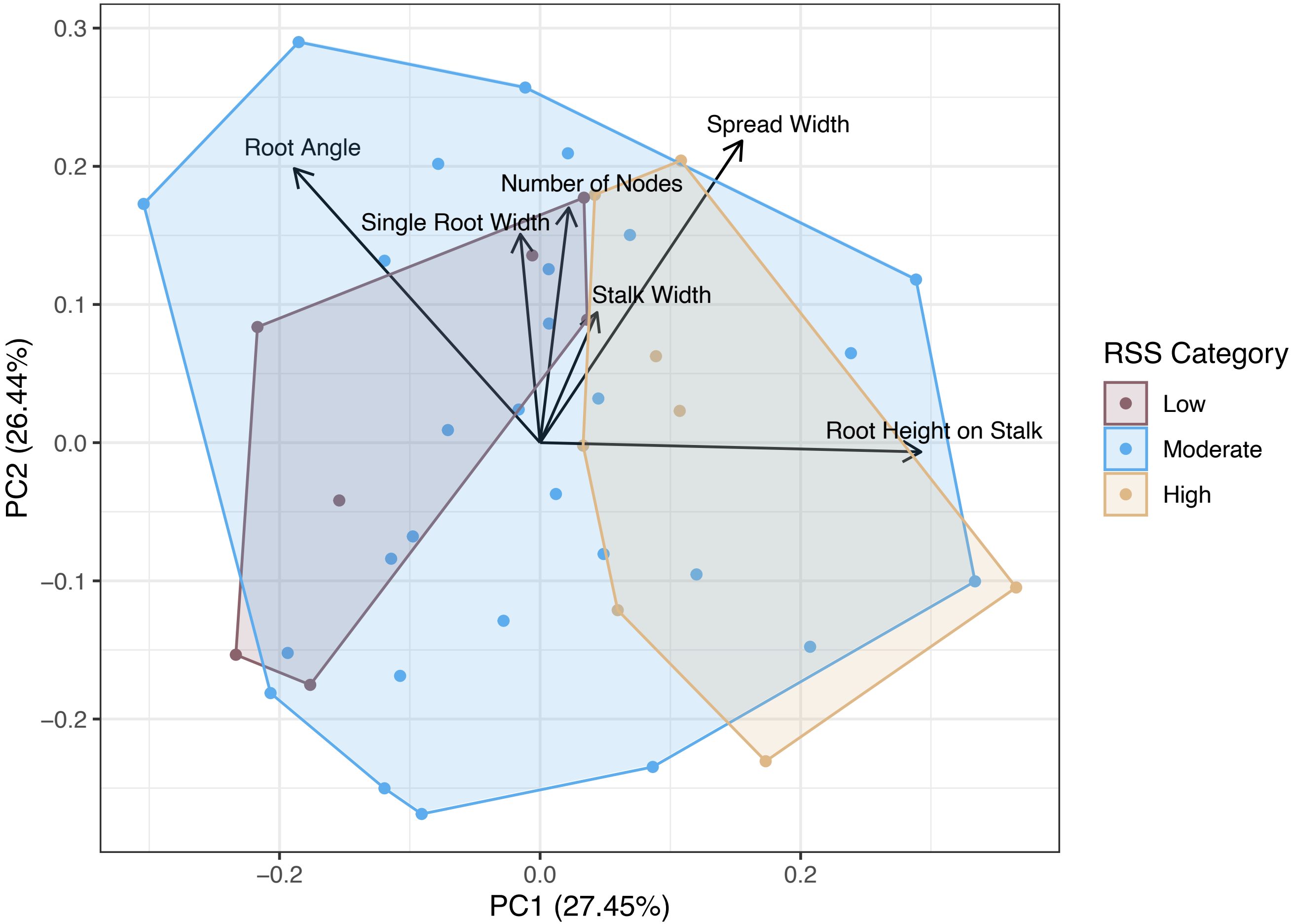


Figure S6. A PCA with trait loadings showed that inbred genotypes that were categorized as having a high root system stiffness had a greater stalk width and larger above-ground root systems (i.e., brace roots attached higher on the stalk at a steeper angle and wider spread width).

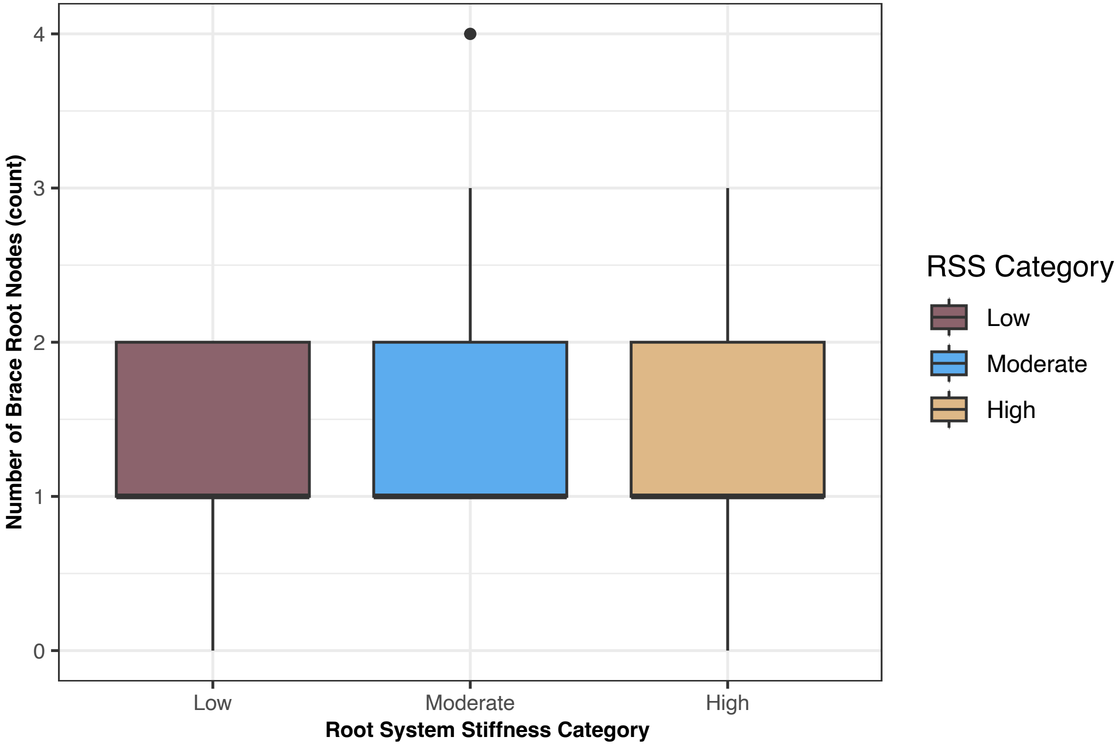


Figure S7. The number of brace root nodes in the soil was not significantly different among root system stiffness categories.

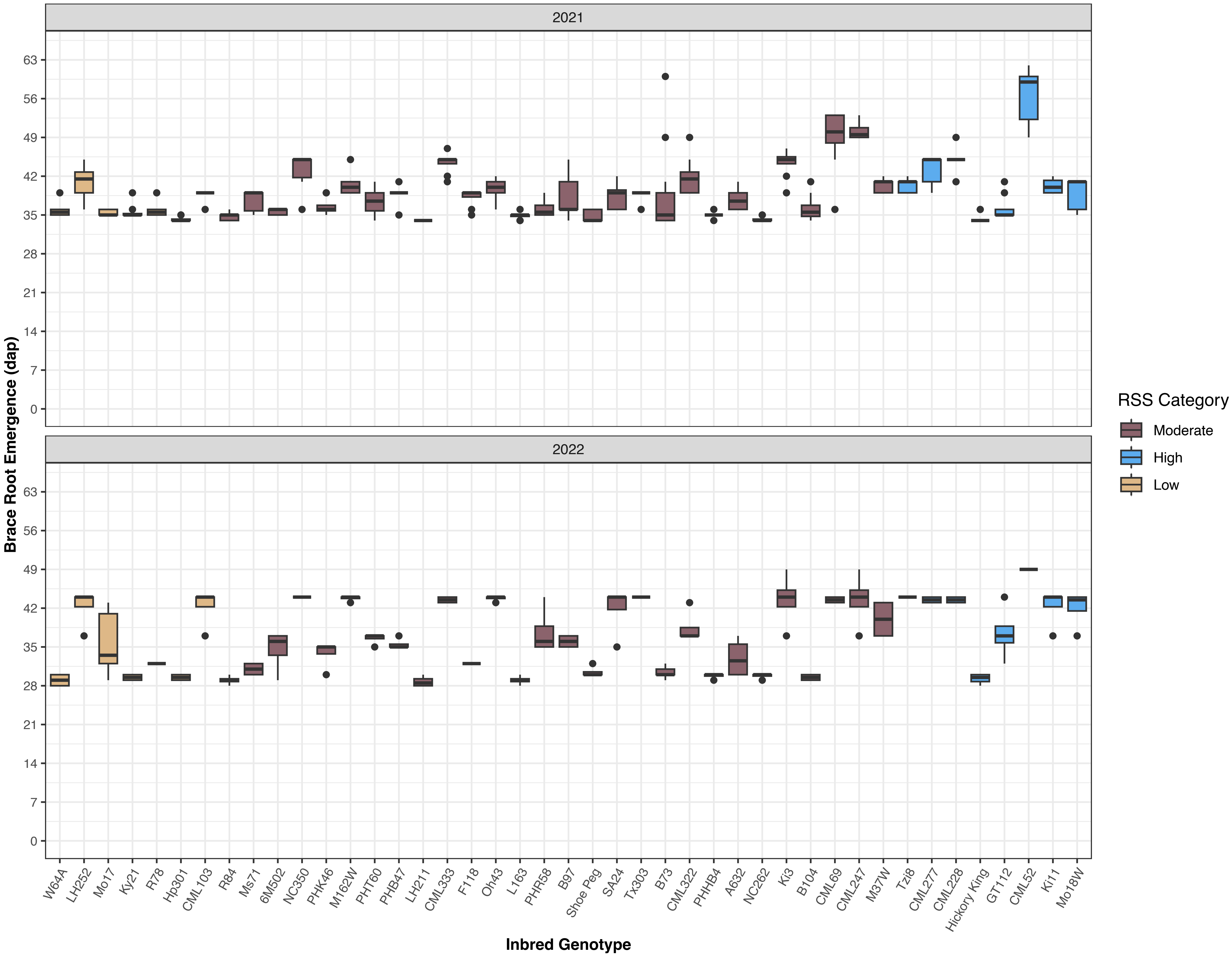


Figure S8. Inbred genotypes varied for brace root emergence date. Brace roots emergence date was dependent on genotype, year, and their interaction. Colors indicate root system stiffness categories.

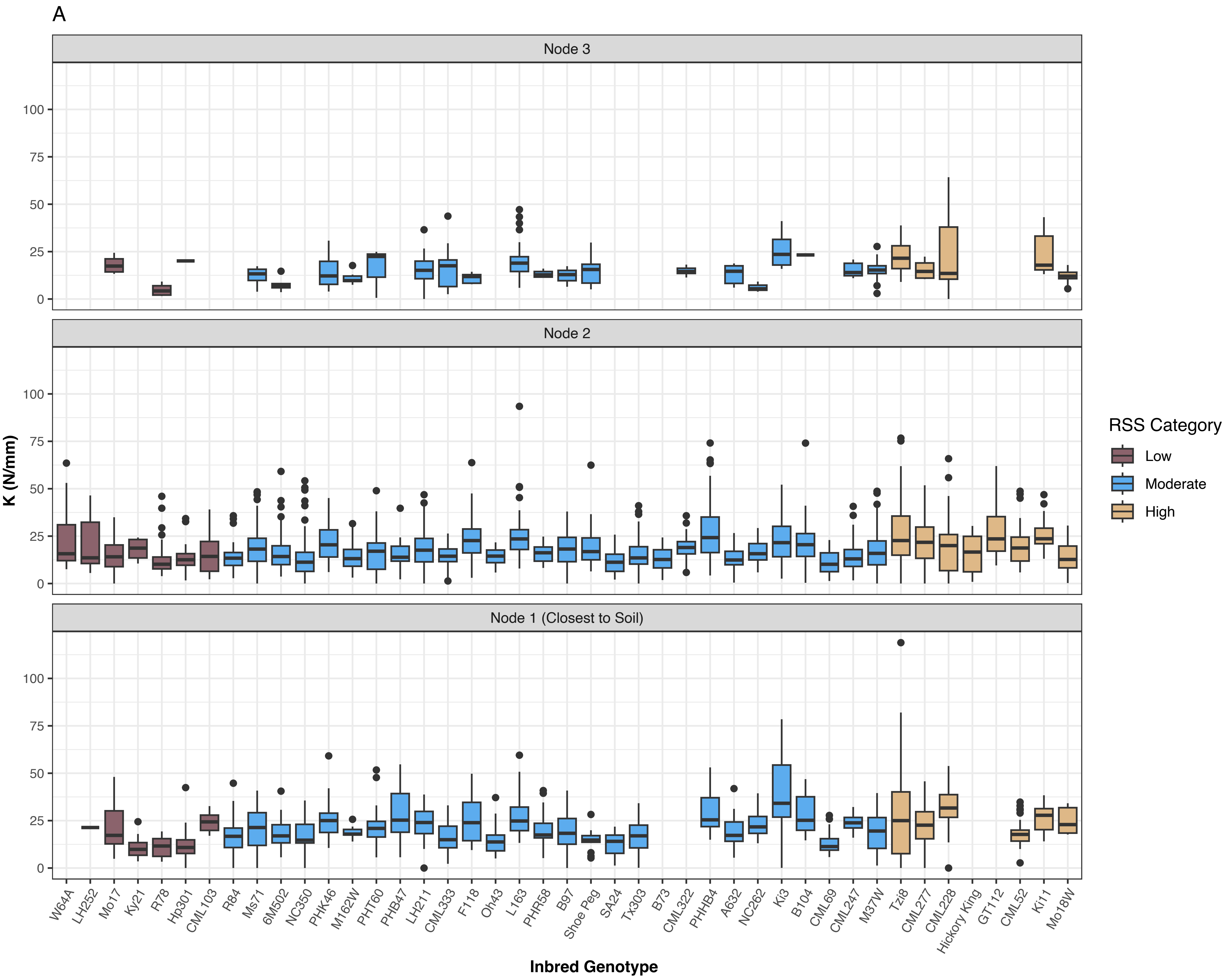

A
Node 3
Node 2
Node 1 (Closest to Soil)
K (N/mm)
0
25
50
75
100
Inbred Genotype
W64A
LH252
Mo17
Ky21
R78
Hp301
CML103
R84
Ms71
6M502
NC350
PHK46
M162W
PHT60
PHB47
LH211
CML333
F118
Oh43
L163
PHR58
B97
Shoe Peg
SA24
Tx303
B73
CML322
PHHB4
A632
NC262
Ki3
B104
CML69
CML247
M37W
Tzi8
CML277
CML228
Hickory King
GT112
CML52
Ki11
Mo18W
RSS Category
Low
Moderate
High

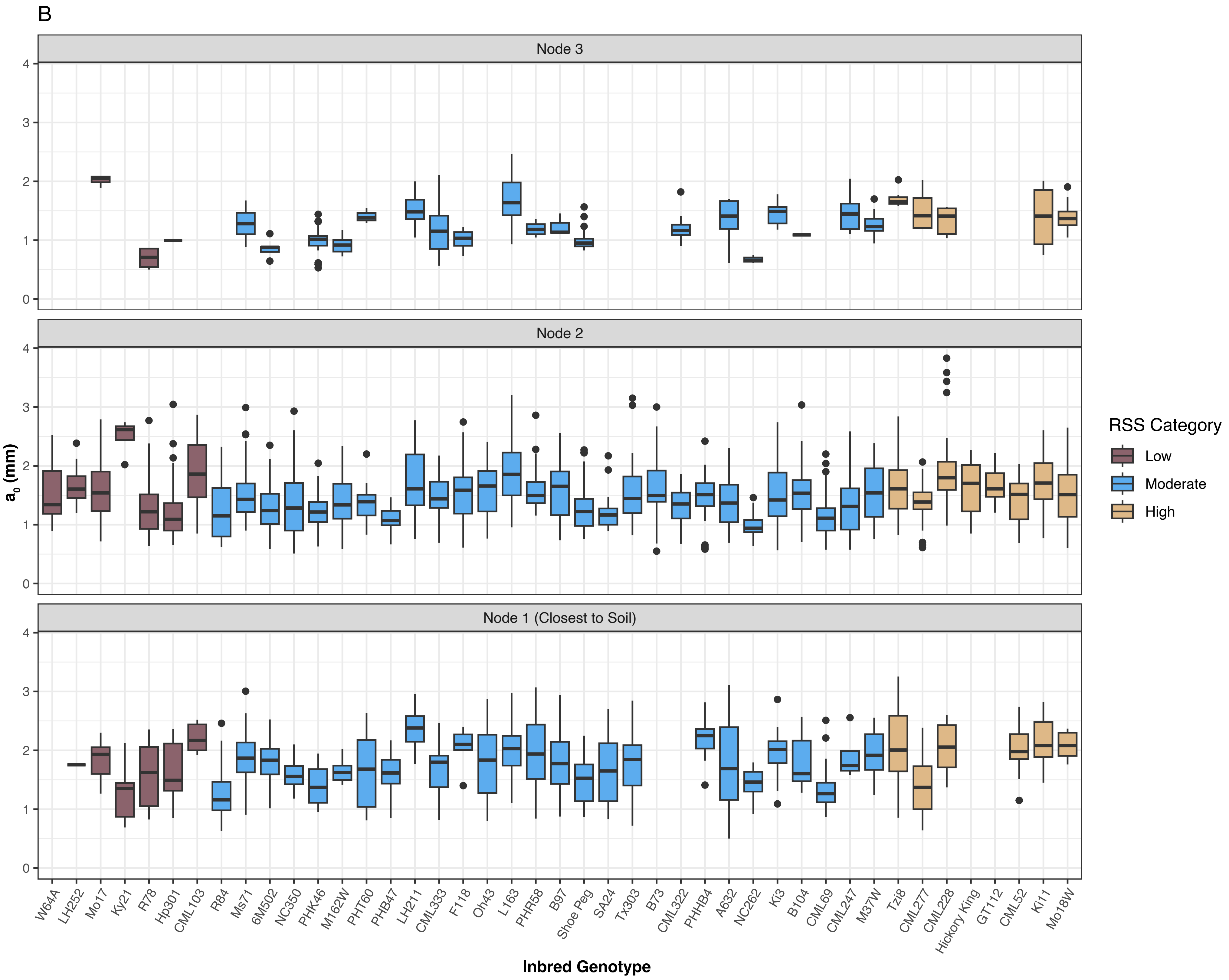

B
Node 3
Node 2
Node 1 (Closest to Soil)
$a_0$ (mm)
0
1
2
3
4
Inbred Genotype
W64A
LH252
Mo17
Ky21
R78
Hp301
CML103
R84
Ms71
6M502
NC350
PHK46
M162W
PHT60
PHB47
LH211
CML333
F118
Oh43
L163
PHR58
B97
Shoe Peg
SA24
Tx303
B73
CML322
PHHB4
A632
NC262
Ki3
B104
CML69
CML247
M37W
Tzi8
CML277
CML228
Hickory King
GT112
CML52
Ki11
Mo18W
RSS Category
Low
Moderate
High

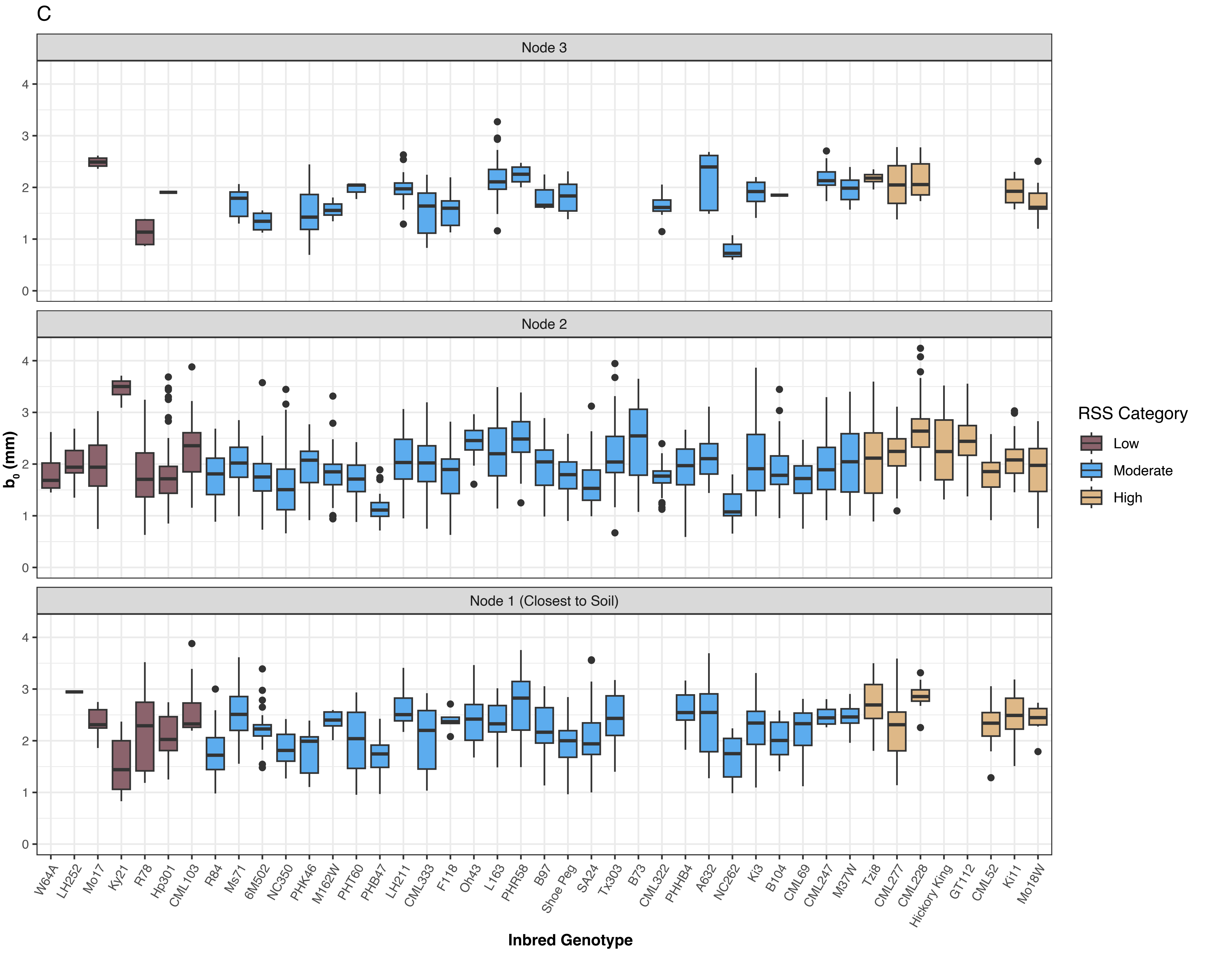

C
Node 3
Node 2
Node 1 (Closest to Soil)
b0 (mm)
Inbred Genotype
W64A
LH252
Mo17
Ky21
R78
Hp301
CML103
R84
Ms71
6M502
NC350
PHK46
M162W
PHT60
PHB47
LH211
CML333
F118
Oh43
L163
PHR58
B97
Shoe Peg
SA24
Tx303
B73
CML322
PHHB4
A632
NC262
Ki3
B104
CML69
CML247
M37W
Tzi8
CML277
CML228
Hickory King
GT112
CML52
Ki11
Mo18W
RSS Category
Low
Moderate
High

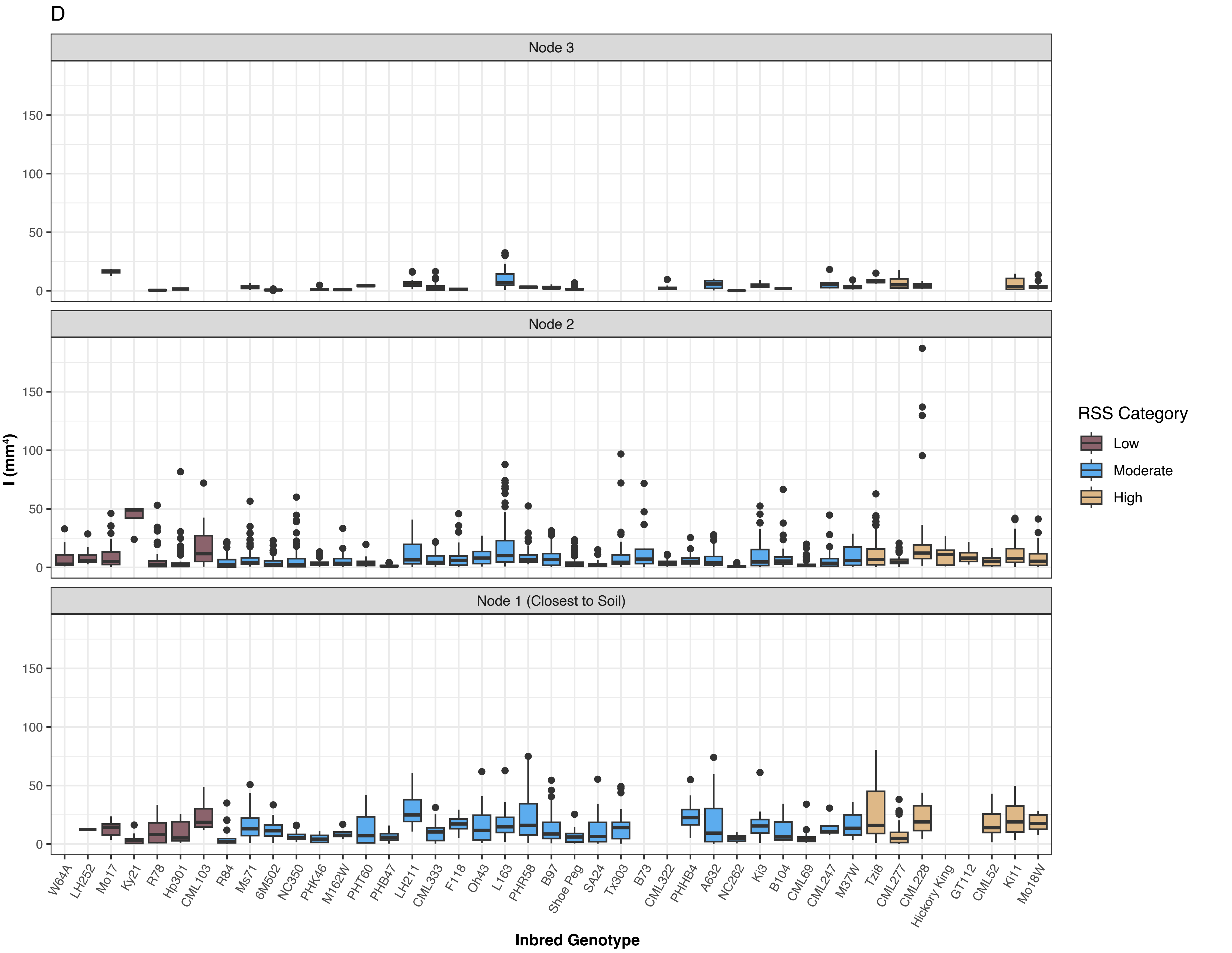
D
Node 3
Node 2
Node 1 (Closest to Soil)
I (mm⁴)
150
100
50
0
W64A
LH252
Mo17
Ky21
R78
Hp301
CML103
R84
Ms71
6M502
NC350
PHK46
M162W
PHT60
PHB47
LH211
CML333
F118
Oh43
L163
PHR58
B97
Shoe Peg
SA24
Tx303
B73
CML322
PHHB4
A632
NC262
Ki3
B104
CML69
CML247
M37W
Tzi8
CML277
CML228
Hickory King
GT112
CML52
Ki11
Mo18W
Inbred Genotype
RSS Category
Low
Moderate
High

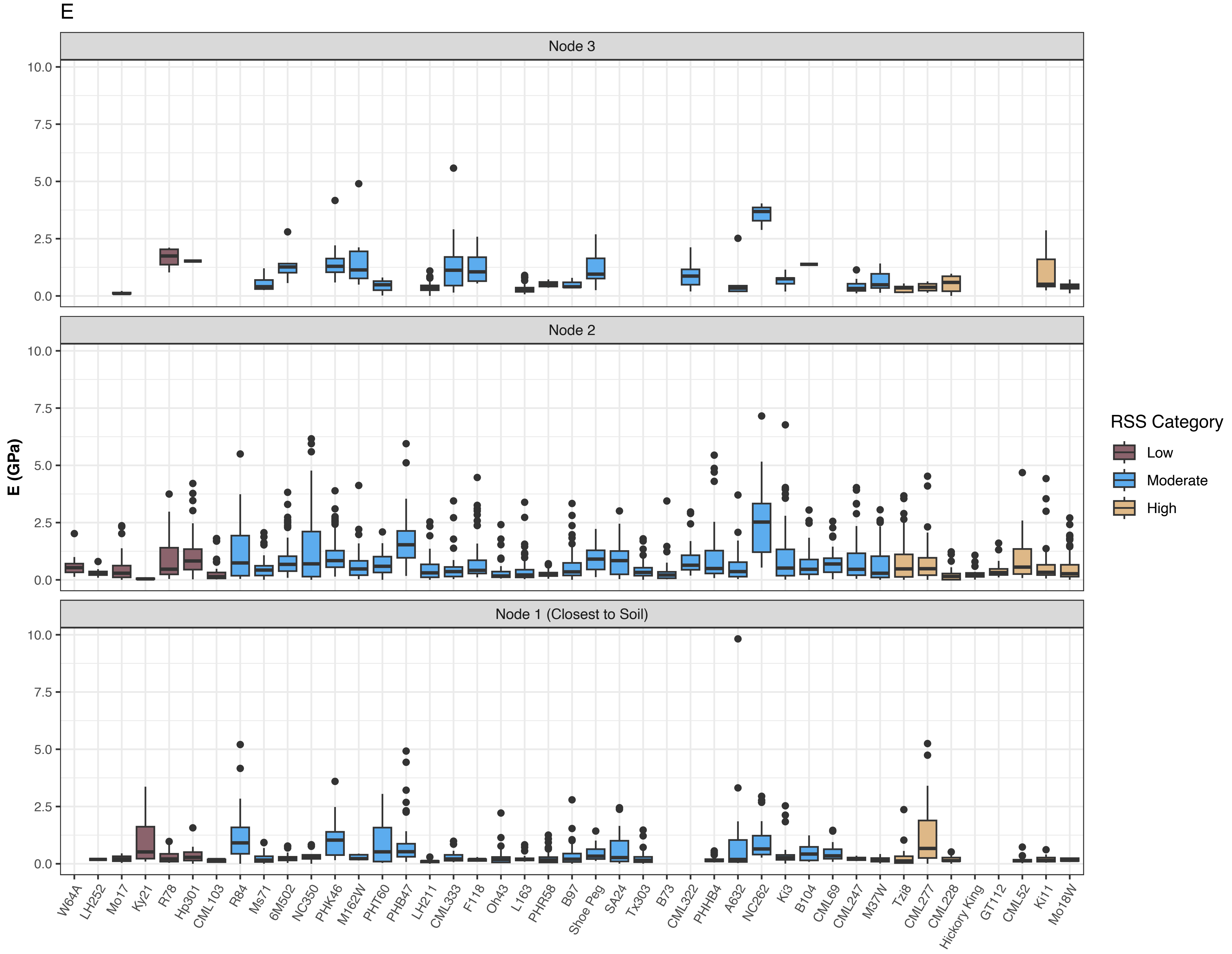


Figure S9. Brace root mechanics varied by inbred genotype and node. (A) structural stiffness (K), (B) the minor radius axis parallel to the plane of displacement (a0), (C) the major radius axis perpendicular to the plane of displacement (b0), (D) the second moment of area (I), and (E) the bending modulus (E). Colors indicate root system stiffness categories.

A

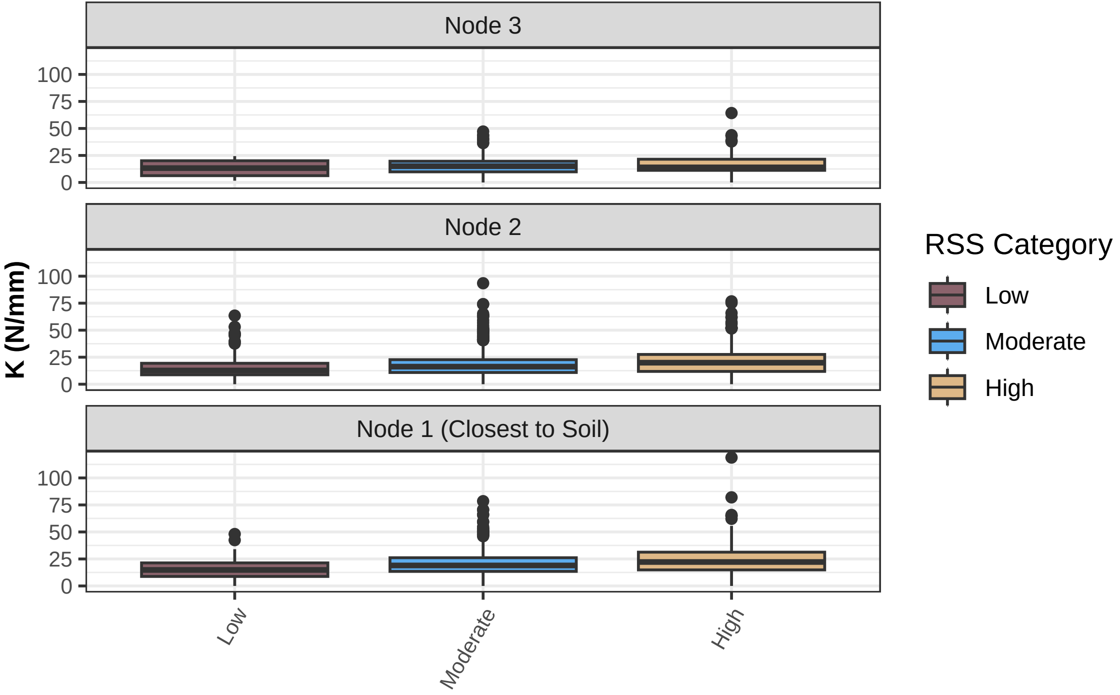
Node 3
Node 2
Node 1 (Closest to Soil)
K (N/mm)
0
25
50
75
100
Low
Moderate
High
RSS Category
Low
Moderate
High

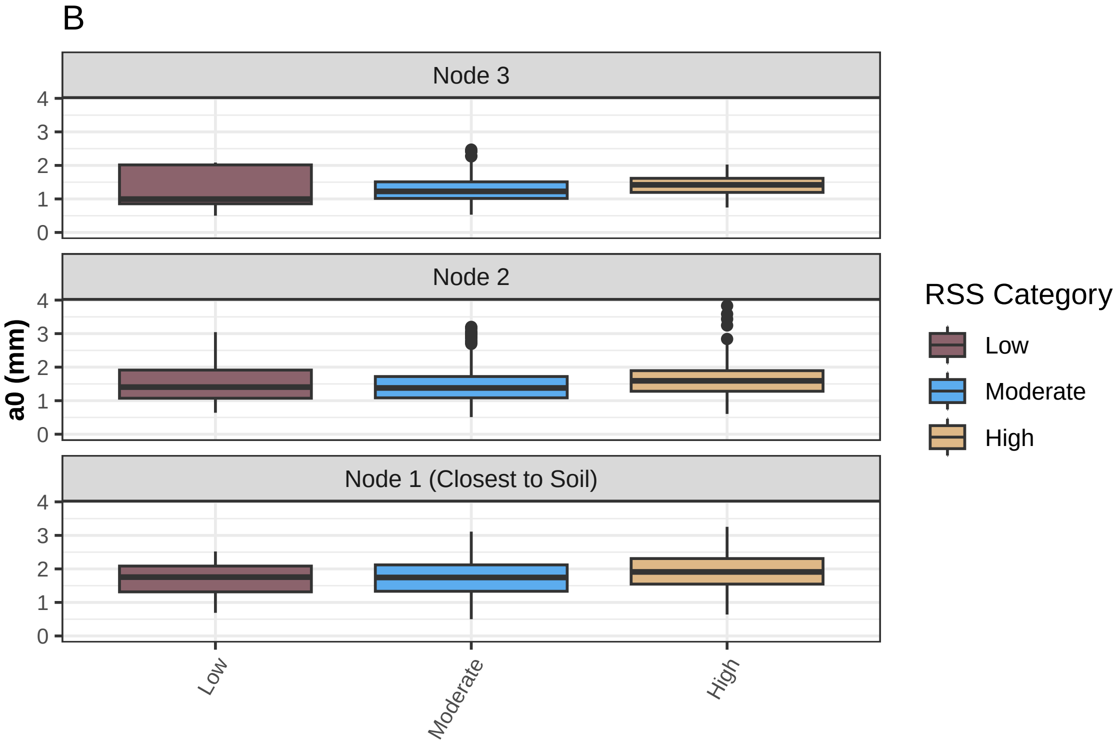
B
Node 3
Node 2
Node 1 (Closest to Soil)
a0 (mm)
0
1
2
3
4
Low
Moderate
High
RSS Category
Low
Moderate
High

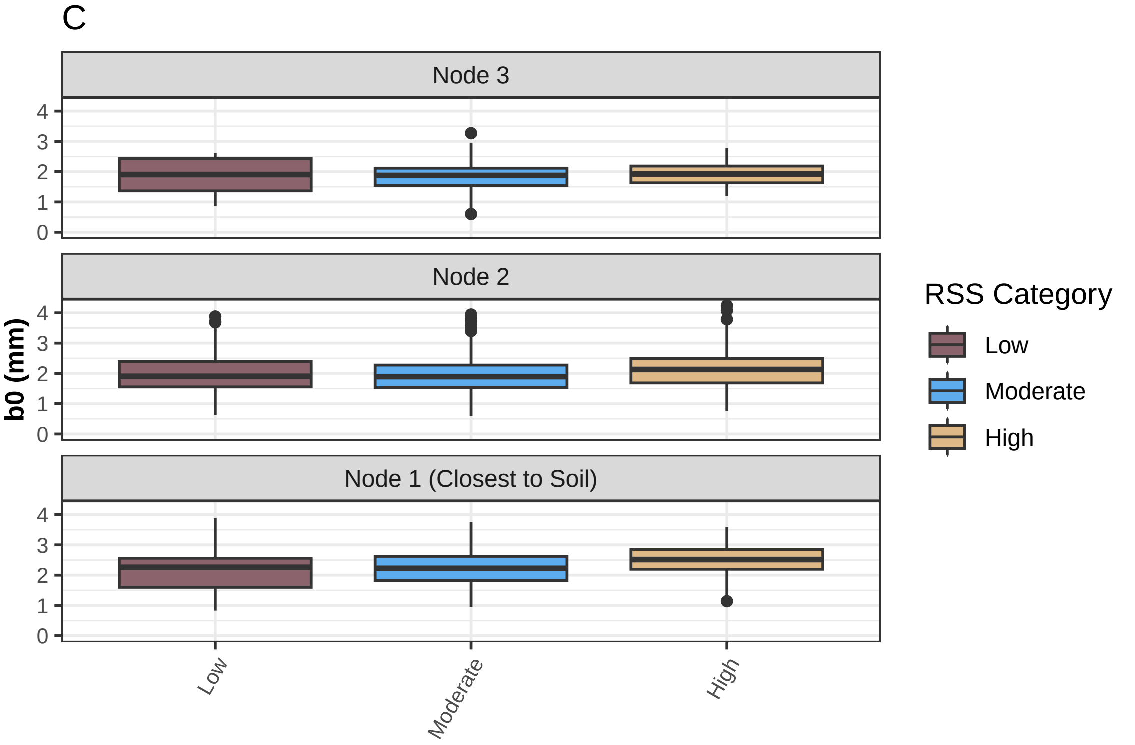

C
Node 3
Node 2
Node 1 (Closest to Soil)
b0 (mm)
0
1
2
3
4
Low
Moderate
High
RSS Category
Low
Moderate
High

D

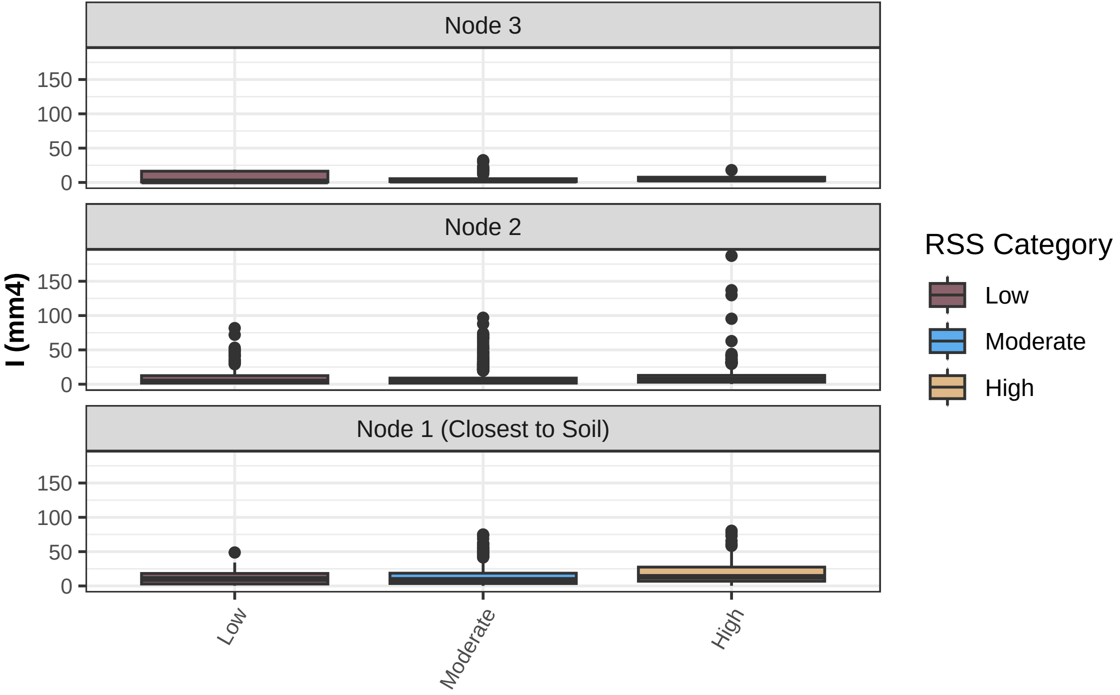

Node 3
Node 2
Node 1 (Closest to Soil)
I (mm4)
0
50
100
150
Low
Moderate
High
RSS Category
Low
Moderate
High

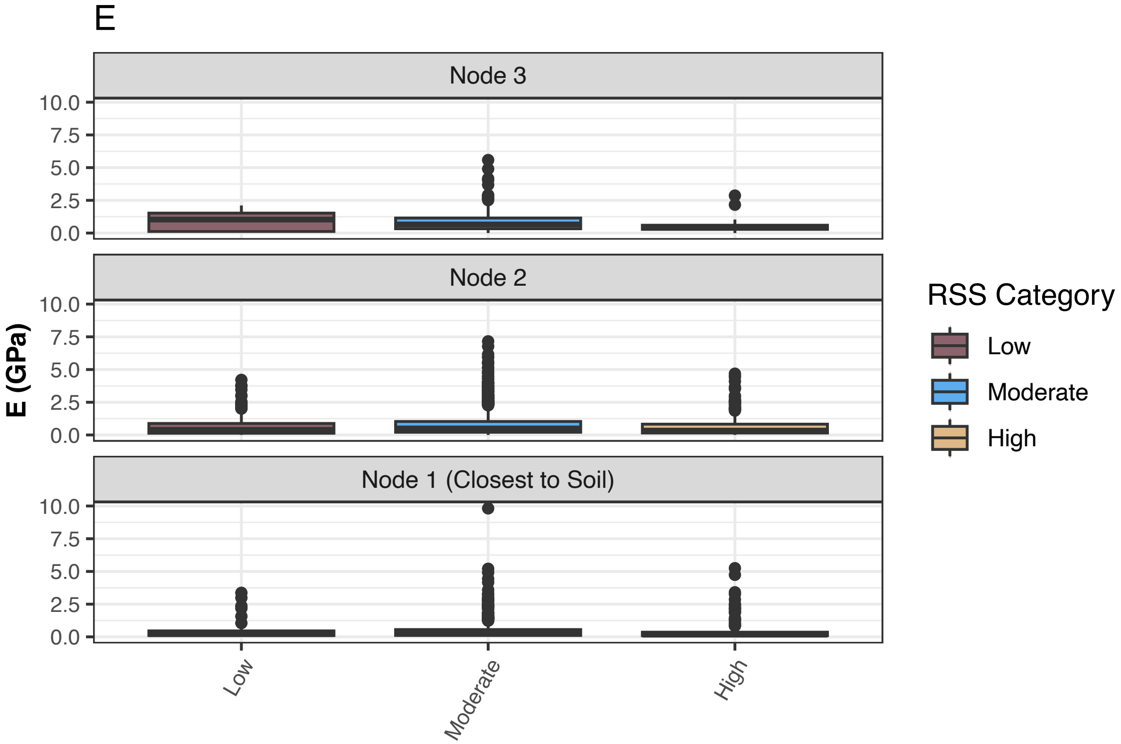


Figure S10. Node-specific variation in brace root mechanics by root system stiffness categories. (A) structural stiffness (K), (B) the minor radius axis parallel to the plane of displacement (a0), (C) the major radius axis perpendicular to the plane of displacement (b0), (D) the second moment of area (I), and (E) the bending modulus (E). Colors indicate root system stiffness categories.

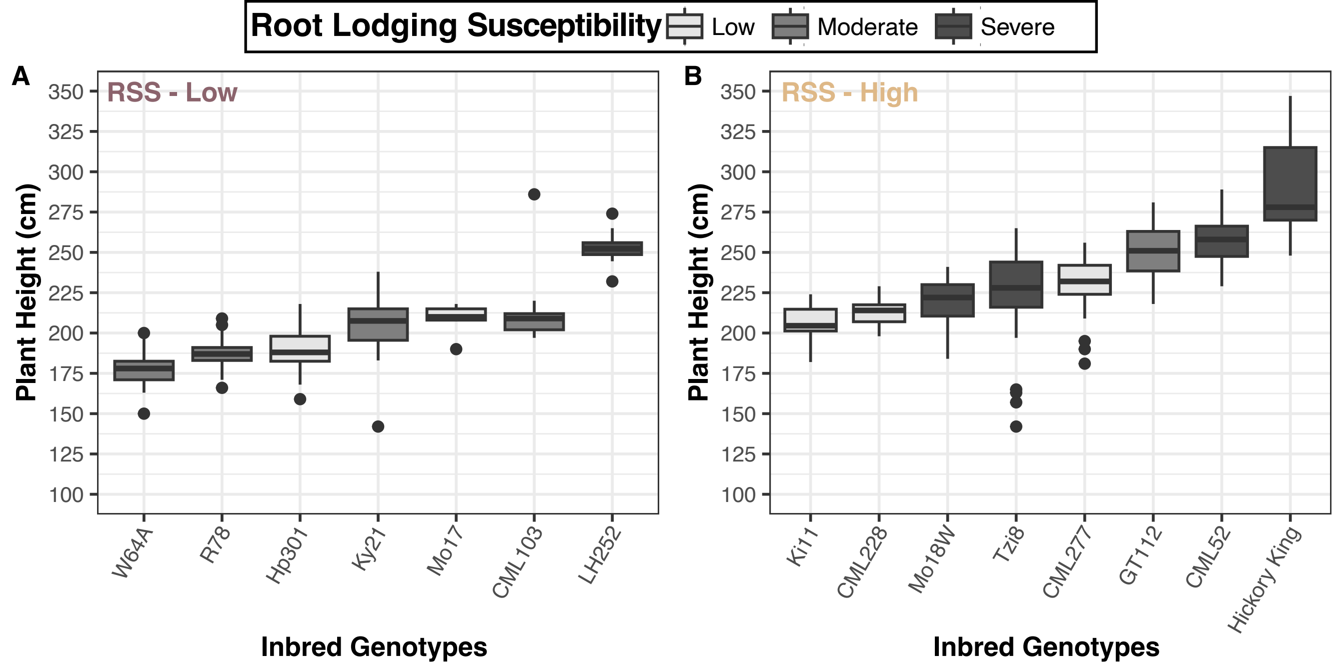


Figure S11. Plant height data for low and high root system stiffness categories. (A) Among the inbred genotypes in the low root system stiffness category, one is also categorized as high root lodging susceptible and is the tallest. (B) Among the inbred genotypes in the high root system stiffness category, those classified as low root lodging susceptible are among the shortest. RSS = root system stiffness.

**Supplementary Figure and Table Literature Cited**

Flint-Garcia, S. A., A.-C. Thuillet, J. Yu, G. Pressoir, S. M. Romero, S. E. Mitchell, J. Doebley, et al. 2005. Maize association population: a high-resolution platform for quantitative trait locus dissection. *The Plant Journal* 44: 1054–1064.

Liu, K., M. Goodman, S. Muse, J. S. Smith, E. Buckler, and J. Doebley. 2003. Genetic Structure and Diversity Among Maize Inbred Lines as Inferred From DNA Microsatellites. *Genetics* 165: 2117–2128.

Smith, Abraham George, Eusun Han, Jens Petersen, Niels Alvin Faircloth Olsen, Christian Giese, Miriam Athmann, Dorte Bodin Dresbøll, and Kristian Thorup-Kristensen. 2022. "RootPainter: Deep Learning Segmentation of Biological Images with Corrective Annotation." *The New Phytologist* 236 (2): 774–91.
